# Information Shielding Through Neural Deactivation: A Fundamental Mechanism of Human Information Processing

**DOI:** 10.1101/2025.02.04.636404

**Authors:** Zhenhong He, Yifan Du, Ziqi Fu, Youcun Zheng, Nils Muhlert, Barbara Sahakian, Rebecca Elliott

**Author notes:** Corresponding author. Division of Psychology and Mental Health, School of Health Sciences, University of Manchester, Manchester, M13 9PT, United Kingdom. The authors declare that there are no conflicts of interest in relation to this study.

## Abstract

Information processing is a core issue in cognitive neuroscience. The brain can either accept or shield information. However, information shielding has traditionally been regarded as a byproduct of other cognitive processes that prepare brain areas for activation, with the accompanying brain deactivation seen as merely supportive. Here, we offer an alternative perspective, demonstrating that people can directly perform information shielding without engaging other cognitive processes. Using information shielding as a novel investigative lens, we provide direct evidence that deactivation serves as a primary mechanism in its own right. We found that information shielding was characterized by a deactivation-dominant neural dynamic, with initial executive control region activation giving way to increased sensory region deactivation, especially after a 3-second boundary. Neural decoding results further confirmed this deactivation trend. This deactivation pattern was consistent across both auditory and visual modalities. Furthermore, non-invasive brain stimulation demonstrated a causal relationship between the deactivation pattern and information shielding, and we found that this deactivation mechanism extended to general unnecessary information processing, regardless of whether the necessity was determined objectively or subjectively. This challenges the conventional emphasis on activation as the primary mode of information processing and opens new fundamental avenues for understanding inhibitory cognitive processes.

## Introduction

Cognitive neuroscience fundamentally focuses on understanding how the human brain processes information^1^. A deep understanding of information processing is therefore crucial to addressing the core issues in this field.

Information processing includes two aspects: the additive direction, which focuses on information intake and enhancement, and the subtractive direction, which focuses on information rejection and attenuation. While research has predominantly explored the additive aspect (exemplified by key studies such as ^2,3^), the subtractive aspect has received comparatively less attention. In the limited number of relevant studies, information subtraction was often viewed as merely a byproduct of other cognitive processes (e.g., focusing on relevant stimulus A naturally leads to ignoring irrelevant stimulus B^4–6)^. However, information subtraction can also function as a primary process; in other words, we do not need to ignore B through focusing on A—we can directly shield B from processing, even if it is relevant to us. An illustrative example is when a person dining at a restaurant overhears an uncomfortable nearby conversation about themselves. Despite the conversation’s relevance, the individual can choose to block it out to maintain their dining experience and emotional balance. This active subtraction capability represents a fundamental aspect of neural information processing, enabling the brain to maintain cognitive autonomy independent of external information relevance.

We refer to this subtraction of information intake as information shielding (IS).

Unlike related concepts such as selective attention^7^, the “cocktail party” effect^8^, and biased competition^9^, which achieve information subtraction as a byproduct of attention enhancement, IS represents a more fundamental neural mechanism that directly targets information rejection. While attention-based mechanisms require a contrast between to-be-attended and to-be-ignored information^10^, IS operates independently as a pure subtraction process. This distinction reveals IS as a primary rather than subsidiary cognitive process - a basic neural operation as fundamental as information enhancement On the neural underpinnings, the brain exhibits three primary states of neural activity when responding to specific demands: responsive (activated), inactive, or suppressed (deactivated)^11–13^. While the first two states—reflecting active engagement and disengagement—have been extensively studied (e.g., in the default mode network and task-positive networks^14,15^), the third state of suppression or deactivation remains poorly understood. Existing interpretations often regard deactivation as a mechanism of neural efficiency, where the brain strategically downregulates activity to conserve resources. These interpretations typically cast deactivation in a supportive role, either to enhance relevant information processing or to prevent cognitive overload^14,16–19^. Our research aims to challenge this conventional view by exploring whether deactivation is not simply a supporting process, but rather a primary mechanism of information processing—especially in the context of reduced processing demands. IS provides an opportunity to study this deactivation-based mechanism directly, as it inherently involves diminishing the intake and processing of unwanted information.

To investigate this hypothesis, we developed an active IS paradigm targeting both auditory and visual modalities—the main channels through which humans process information—to delineate the neural substrates underlying IS processes. We used functional magnetic resonance imaging (fMRI) to scan human participants as they engaged in either natural processing (NP) of voice or picture stimuli, or applied specific cognitive strategies (inhibition, distraction, and distancing) to establish information shielding (IS). Additionally, through electroencephalography (EEG), we identified a temporal boundary around 3 seconds post-stimulus onset. Around this boundary, our findings revealed that IS-induced changes in brain activity were characterized by a deactivation-dominant dynamic. Moreover, we observed consistent cross-modal patterns in the neural representation of IS, suggesting that the brain’s response to IS is both robust and modality-independent. Finally, we confirmed the causal relationship between the deactivation pattern and IS by transcranial magnetic stimulation (TMS), and extend our findings beyond IS to the broader context of reduced processing needs, demonstrating that the brain consistently engages in a deactivation pattern whenever information is deemed “unnecessary”. Together, these findings constitute the first direct empirical evidence the mechanisms underlying IS in the human brain. The results establish a foundational framework for understanding the underlying deactivation- based neural processes.

### Information shielding: a deactivation-dominant neural dynamic in auditory and visual modalities

While studies on information processing have predominantly focused on the aspects of information intake and enhancement, such as improving efficiency and quality^2,3^, the attenuation and shielding of information—and especially their underlying mechanisms—have received far less attention. Limited research has often treated IS merely as a byproduct of other cognitive processes^10,20^. Our study demonstrates that IS is an independent, primary process governed by a fundamental deactivation mechanism that shows remarkable consistency across sensory modalities. We designed auditory and visual IS (AIS and VIS) tasks to illustrate the existence, neural representations, and temporal dynamics of IS across these modalities (Figure 1A; see “Methods-Behavioral task”).

**Figure. 1.**
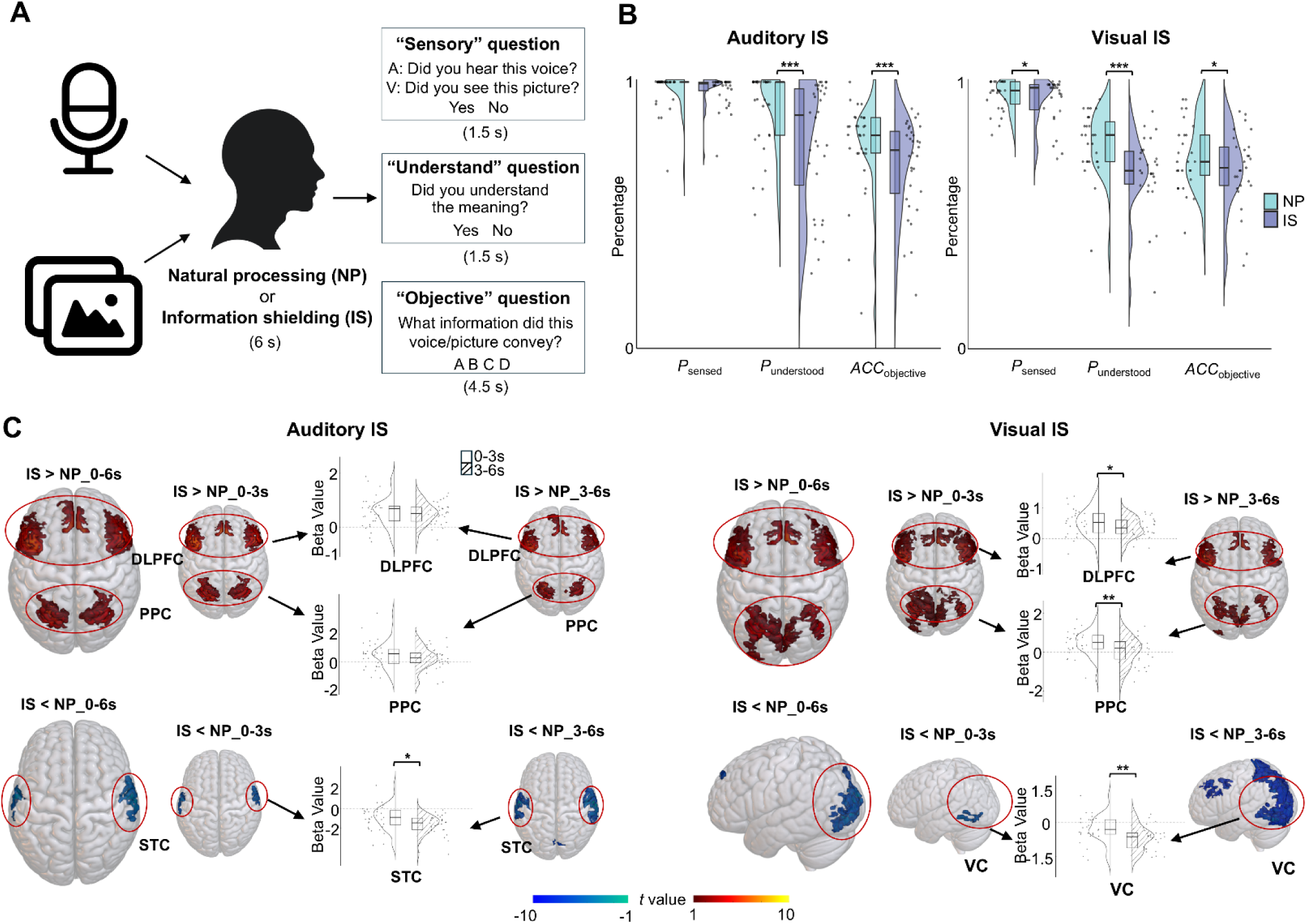
Experimental design and brain responses during information shielding (IS). **A**, Schematic of the auditory AIS and visual IS (VIS) experiments. Participants engaged in either natural processing (NP) or IS of voice or picture stimuli, followed by answering three questions to assess sensory perception, understanding, and objective information conveyed. **B.** Behavioral results of AIS (left panel) and VIS (right panel). Raincloud plots represent the distribution of responses rates of perceiving (*P*_sensed_) and understanding the stimuli (*P*_understood_), along with accuracy in identifying the conveyed information (*ACC*_objective_). Box plots represent median and interquartile ranges of these values. **C,** fMRI temporal dynamics of the AIS (left panel) and VIS (right panel). Brain maps show IS-induced activations in dorsolateral prefrontal cortex (DLPFC), posterior parietal cortex (PPC), superior temporal cortex (STC), and visual cortex (VC) during early (0-3s) and late (3-6s) periods. Color scale indicates *t*-values for activation (red, 1 to 10) and deactivation (blue, -1 to -10). Raincloud plots show IS-related activation/deactivation (beta values) in each region, segmenting into early and late periods. \**p* < 0.05, \*\**p* < 0.01, \*\*\**p* < 0.01.

We observed consistent IS phenomena and effectiveness across both auditory and visual modalities in behavioral measurements. Participants reported significantly lower rates of perceiving and comprehending the stimuli, along with reduced accuracy in identifying the conveyed information after applying IS strategies (Figure 1B; see “Results-Behavioral Results”). This cross-modal concordance in behavioral outcomes indicates that IS operates in a similar manner to disrupt information processing, irrespective of the sensory modality.

Investigation of neural representations revealed a consistent pattern of top-down control across auditory and visual modalities. During both AIS and VIS, we observed enhanced activation in executive control regions (dorsolateral prefrontal cortex, DLPFC; posterior parietal cortex, PPC) coupled with deactivation in the corresponding sensory regions (superior temporal cortex, STC/visual cortex, VC; Extended data 1- Figure 1A,B). Notably, successful IS implementation correlated with stronger executive control region activation and more pronounced sensory region deactivation in both modalities (Extended data 1- Figure 1C,D). This parallel neural architecture across sensory domains supports a supramodal top-down regulatory mechanism underlying IS. We further investigated the temporal dynamics of IS. We identified a critical boundary approximately 3 seconds post-stimulus onset, marking a significant transition in neural activity patterns that was consistently observed across modalities. The identification of this boundary was informed by previous literature, which posits that executive control processes can be initiated rapidly, while sensory and attentional disengagement processes are relatively slower^21–24^. This led to our hypothesis that during IS, the activation of executive control regions might reach its peak earlier, followed by the deactivation of sensory regions, with a potential temporal boundary segmenting the two phases—an effect we expected to observe consistently across modalities. This temporal boundary, identified around 3 seconds post-stimulus onset, was determined using high-temporal-resolution EEG data from the AIS experiment by constructing temporal curves of IS decoding accuracy. The boundary was evident only after a certain exposure duration (not observed in the 3-second stimuli), but once established, it was independent of the total stimulus length (observable across the 6- second and 10-second stimuli; Extended data 1- Figure 1F,G). This IS-related boundary was further validated in the fMRI data of the AIS and VIS experiment, which reveal consistent change points between 3 and 4.5 seconds post-stimulus onset in both executive control and sensory regions (Extended data 1- Figure 1H). These findings together highlight a critical time boundary where significant neural changes occur during IS, regardless of the sensory domain.

The temporal dynamics surrounding this boundary exhibited striking similarities between auditory and visual modalities. Both IS related activated frontoparietal regions and deactivated sensory areas demonstrated a consistent trend toward deactivation in the later stages of IS processing. Specifically, when segmenting neural responses into pre-3 and post-3 second periods, we observed parallel temporal patterns across modalities: frontal-parietal activation was significant in the early stage but diminished later, while sensory region deactivation became more pronounced after the 3-second mark (Figure 1C). Moreover, individuals who demonstrated more successful IS also showed a more pronounced trend of deactivation in sensory regions in both modalities (Extended data 1- Figure 1J,K). This highlights that successful IS is characterized not only by the magnitude of sensory deactivation but also by the dynamic progression of deactivation across time, a pattern consistent across auditory and visual domains.

Taken together, we identified a temporal deactivation pattern, where frontoparietal activation weakened while sensory region deactivation increased over time, suggesting that both executive and sensory regions trend towards deactivation in the later stages of IS. Importantly, this temporal deactivation pattern showed remarkable consistency across both auditory and visual modalities, suggesting a domain-general mechanism of IS where executive control initiates rapidly to facilitate subsequent sensory disengagement.

### The cross-modality consistency of information shielding

In both visual and auditory modalities, we observed similar brain deactivation patterns during active IS. This observation led us to systematically investigate the cross-modal nature of IS mechanisms at multiple levels of analysis.

First, we examined whether neural patterns associated with IS could transfer from one modality to another. Using cross-modal multivariate pattern analysis (MVPA; see “Methods-Data recording and analysis-MRI data-MVPA”), we found that IS exhibited cross-modal generalizability, whereas NP did not. Specifically, to assess whether there is a unified code for AIS and VIS, we trained a decoder to discriminate between IS and NP in the auditory modality and tested its performance on visual modality, and vice versa. The results showed that that neural coding during IS generalized across both modalities, indicating a shared, cross-sensory representation of IS. In contrast, no such generalization was observed during the NP condition. Furthermore, compared to the early 3-second period, neural coding during IS showed greater modality generalizability than NP during the late 3-second period, suggesting that the cross-modal shielding state becomes more established over time (Figure 2A).

**Figure 2.**
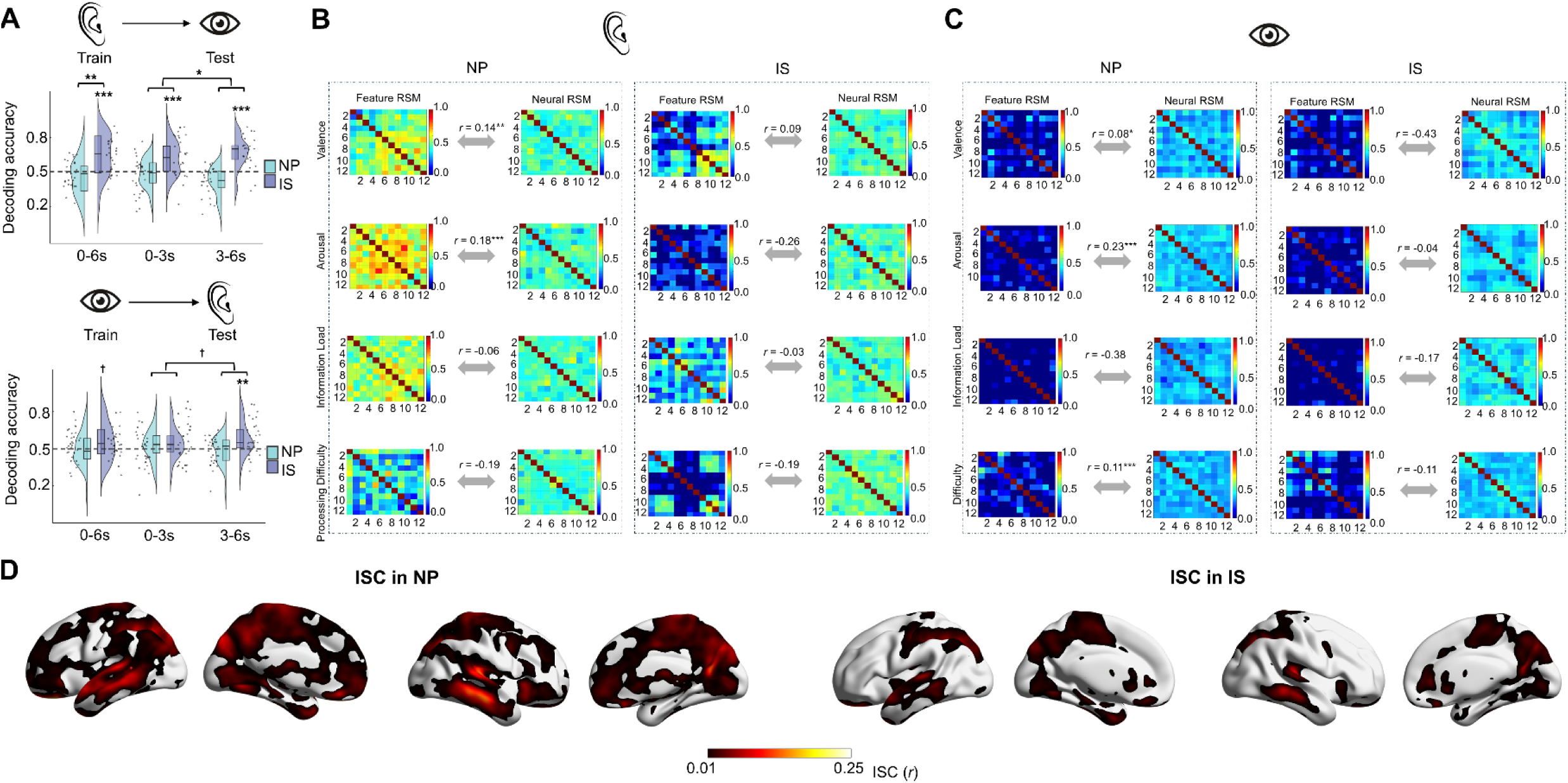
The cross-modality consistency of IS. **A**, Decoding performance for cross- modal decoding, trained on auditory modality and tested on visual modality (top panel) and vice versa (bottom panel). Raincloud plots show the distribution of decoding accuracy. Asterisks (*) indicate significant decodability in each region of interest (ROI). Connecting lines with * represent significant differences between conditions. **B-C**, Correlation analysis between the feature RSM and neural RSM under NP and IS conditions in auditory (B) and visual (C) modalities. **D**, Cross-modal ISC analysis results. Brain maps depict regions showing cross-modal synchronized activity, with the color scale indicating ISC-values (red, 0.01 to 0.25). \**p* < 0.05, \*\**p* < 0.01, \*\*\**p* < 0.01.

Second, we investigated how IS affects the brain processing of various information features and whether these effects show cross-modal consistency. We found that IS similarly reduces the processing of certain features across modalities. To assess this, we conducted representational similarity analysis (RSA; see “Methods-Data recording and analysis-MRI data-RSA”) to examine how subjective information features—such as valence, arousal, processing difficulty, and information load—related to neural patterns during both IS and NP conditions. By constructing representational similarity matrices (RSMs) for both the subjective features and neural activity, we analyzed the alignment between feature-based RSMs and neural RSMs to identify which features could be effectively shielded and which might resist shielding. Our analysis revealed that, compared to the NP condition, IS significantly reduced the impact of emotional features (valence and arousal) on neural activity. This diminished sensitivity to emotional features was consistent across both auditory and visual modalities (Figure 2B,C), indicating that IS applies a modality-consistent mechanism for degrading specific information features in the brain. Interestingly, IS also reduced the impact of processing difficulty on neural activity, but this effect was observed only in the visual modality.

Third, we explored why the IS-related findings appear to be modality-independent. We hypothesized that this may be due to the presence of cross-modal resonance brain regions. We conducted intersubject correlation (ISC; see “Methods-Data recording and analysis-MRI data-Cross modal ISC analysis”) analysis to examine neural synchronization between the auditory and visual groups. The results revealed distinct patterns of neural synchronization during IS compared to the NP condition. While both conditions showed synchronized activity in primary sensory regions (the visual cortex), which served as peak regions for cross-modal synchronization, the IS condition uniquely engaged executive control regions. Specifically, we observed enhanced synchronization in the right middle frontal gyrus (part of the DLPFC) and right supramarginal gyrus (part of the PPC) during IS, with limited involvement in these regions during the NP condition (Figure 2D). These regions of cross-modal neural synchrony could represent a mechanistic basis for the shared neural representations and information processing ways we observed across modalities during IS.

### Multivariate pattern analysis reveals the neural dynamics of information shielding

We next assessed the neural representation of IS across different brain regions through MVPA decoding analysis of fMRI data (Figure 3A; see “Methods-Data recording and analysis-MRI data-MVPA”). The ability to decode specific conditions above chance level indicates that the voxel patterns within a brain region contain sufficient information to distinguish between different cognitive states (IS or NP), demonstrating condition-specific neural coding.

**Figure 3.**
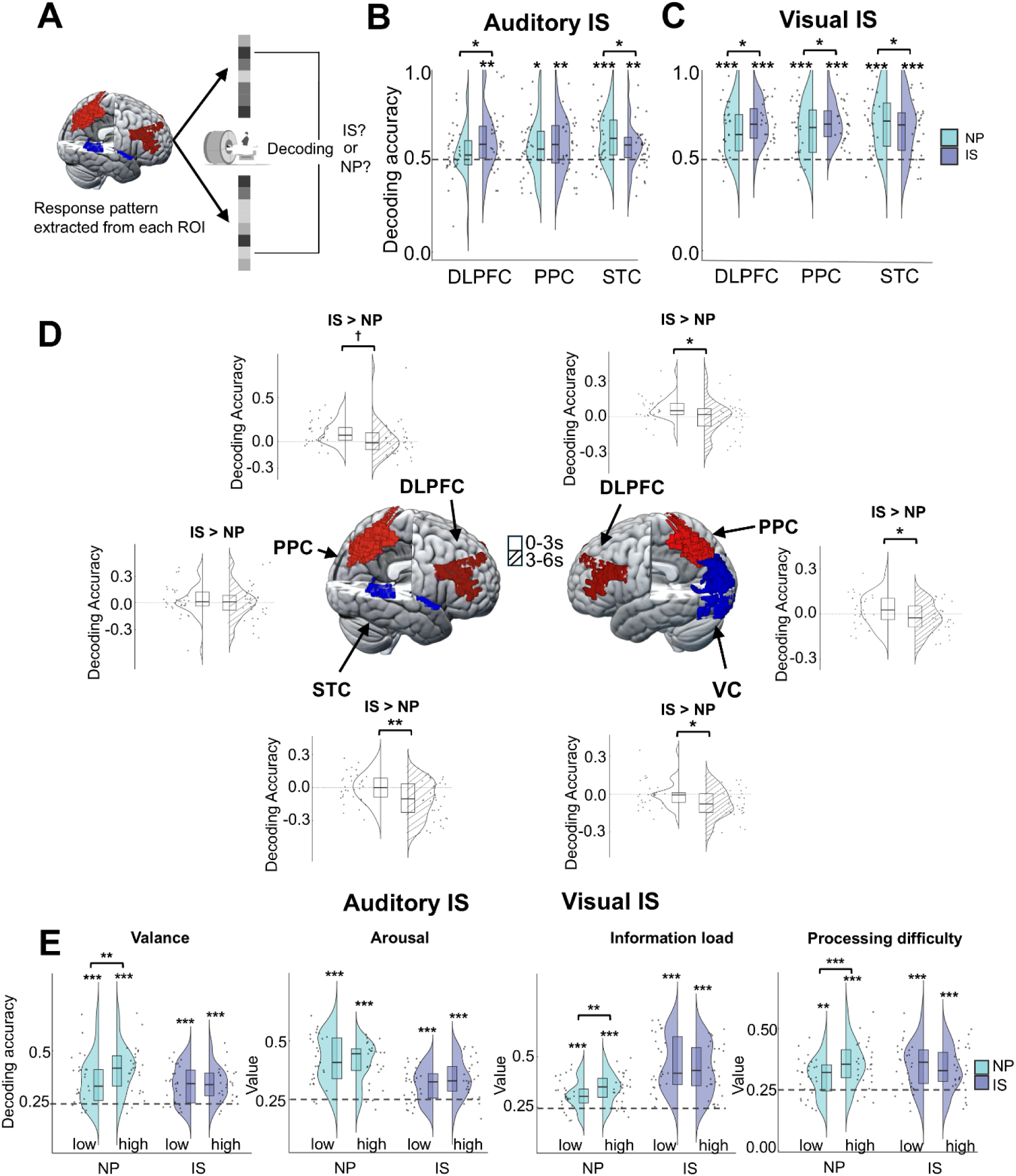
IS related neural coding. **A**, Schematic of fMRI decoding analysis in ROIs, showing cross-modal classification between NP and IS conditions. **B-C,** ROI decoding performance comparing NP and IS conditions in auditory (B) and visual (C) modalities. **D**, ROI decoding accuracy differences (IS > NP) during early (0-3s) and late (3-6s) periods for auditory (left panel) and visual (right panel) modalities. Brain maps show ROI locations in Montreal Neurological Institute (MNI) space: DLPFC (dark red), PPC (light red), and STC/VC (blue). Raincloud plots depict decoding accuracy differences (IS > NP), segmented into early and late periods. **E**, Decoding performance in the combined ROI to discriminate among the following four conditions: high and low intensity of information features under the IS state, and high and low intensity of information features under the NP state. Raincloud plots depict decoding accuracy. \**p* < 0.05, \*\**p* < 0.01, \*\*\**p* < 0.01.

Our analyses revealed three key findings about the neural coding of IS. First, comparing IS versus NP states showed differential decoding patterns that aligned with the activation results from Part 1, demonstrating a similar trend toward deactivation over time. Specifically, while both IS and NP states were robustly decoded from frontoparietal and sensory regions, IS exhibited a distinct pattern: enhanced decoding performance in frontoparietal regions but diminished decoding accuracy in sensory regions (Figure 3B,C). This pattern parallels the activation results, suggesting that regional activation levels correspond to neural coding capacity.

Second, temporal analysis revealed dynamic changes in regional coding capabilities. By segmenting the data into early (<3 seconds) and late (>3 seconds) periods, we found that the initially enhanced frontoparietal decoding during IS attenuated in the later stage. Simultaneously, the reduction in sensory region decoding became more pronounced over time (Figure 3D). These temporal dynamics in coding capabilities mirror the progressive deactivation pattern observed in both executive control and sensory regions during sustained IS.

Third, we examined how IS affects the neural representation of inherent information features. While MVPA could effectively distinguish between high and low levels of valence, arousal, and processing difficulty during NP, these distinctions became significantly less distinguishable during IS (Figure 3E). This finding indicates that IS not only involves deactivation but also systematically dampens the neural encoding of information features, suggesting a broad attenuation of information processing.

Together, these decoding results provide convergent evidence for a deactivation- dominant mechanism in IS, characterized by progressive reduction in both neural activation and coding capacity, particularly in sensory regions, and accompanied by diminished neural representations of information features.

### From information shielding to managing unnecessary information: the brain’s deactivation response

IS reduces the need for information processing. While this relationship between “unnecessary” information processing and neural deactivation is evident, its causal nature and specificity remain uncertain. We established that the deactivation pattern causally contributes to the processing of less-needed information through combined TMS and fMRI methodology. Furthermore, using RSA, we revealed that when the necessity for processing is subjectively high—as in cases where participants were instructed to concentrate on the stimuli—the brain does not similarly engage in the deactivation mode; instead, it exhibits a reversed activation mode. Conversely, under conditions of reduced necessity, whether subjectively induced (during active IS) or objectively determined (when information was not received), the brain similarly engaged in a deactivation mode. These results suggest a close link between the reduced necessity of information processing and the brain’s engagement of the deactivation mode.

Previous neural-behavioral correlation analyses support this hypothesis: individuals demonstrating superior IS performance exhibited more pronounced transitions from low to high deactivation in sensory regions (Extended data-Figure 1J,K), suggesting that decreased information necessity corresponds to enhanced deactivation patterns.

To explore causality, we employed neuromodulation techniques combined with fMRI assessments (see “Methods-Experimental Procedure-Experiment 3(TMS + fMRI)”). Excitatory repetitive TMS over the left DLPFC resulted in better IS modulation of the information processing (Figure 4A,B, and Extended data1-Figure 2A,B) and altered DLPFC activity dynamics. Specifically, the TMS group demonstrated a more pronounced temporal transitions from high to low activation in DLPFC, compared to the sham group (Figure 4C). These findings were corroborated by MVPA results (Extended data1-Figure 2C,D). Moreover, individuals exhibiting more pronounced temporal transitions from low to high deactivation in STC demonstrated enhanced IS capabilities—an effect that was significant in TMS groups but only marginally significant in the sham group (Figure 4D). These findings collectively establish the causal role of the deactivation mode, particularly the temporal dynamics of sensory region deactivation, in the processing of less-needed information. Further, we systematically manipulated the necessity of information processing.

**Figure 4.**
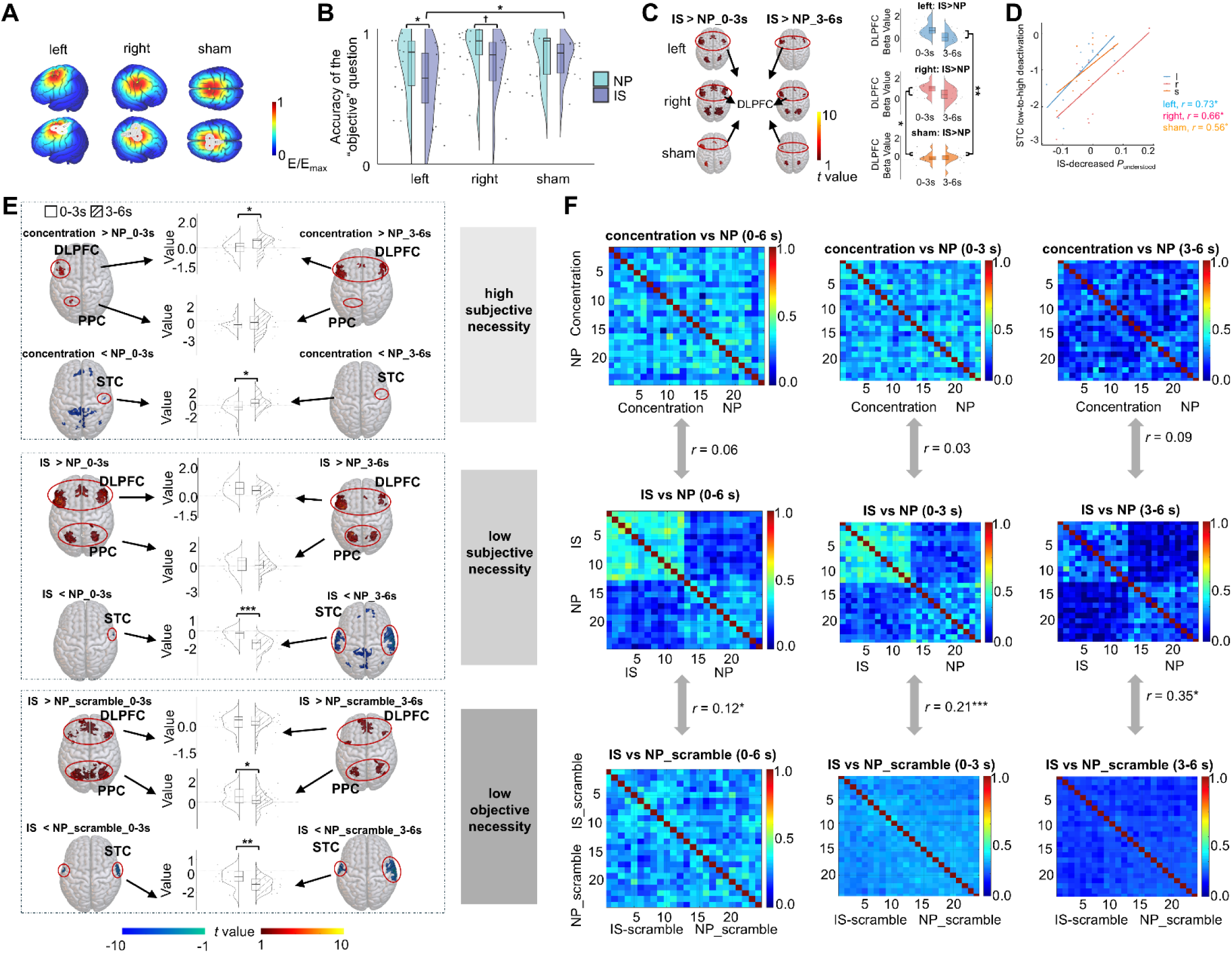
The link between IS and brain deactivation patterns. **A**, TMS electric fields in the three TMS groups visualized using SimNIBS software (www.simnibs.org). The color scale represents the electric field strength, ranging from 0 (blue) to the individual maximum (red). **B**, Behavioral results from the TMS experiment, showing the accuracy of responses in the “objective” question for the three TMS groups. **C,** Brain activation differences (IS > NP) during early (0-3s) and late (3-6s) periods for each TMS group. Raincloud plots show DLPFC activation changes (beta values). **D**, Correlation between IS-related changes in behavioral performance and brain activity patterns for the three TMS groups. **E,** Brain activation and deactivation differences for the following comparisons during early (0-3s) and late (3-6s) periods:: concentration vs. NP (top panel), IS vs. NP (middle panel), IS vs. NP-scramble (bottom panel). Brain maps display activation and deactivation, and raincloud plots depict beta values, segmented into early and late periods. **F,** Correlations between neural RSMs for the following contrasts across different time windows (0-6s, 0-3s, 3-6s) in AIS experiment: concentration vs. NP and IS vs. NP, as well as IS vs. NP and IS vs. NP-scramble.

Firstly, we examined subjective necessity. Under conditions of heightened subjective necessity—as in cases where participants were instructed to concentrate on the information (Extended data1-Figure2 E,F), the contrast analysis (concentration vs. NP conditions) revealed a reversed pattern of neural activity: a low-to-high activation pattern and a high-to-low deactivation pattern (Figure 4E). It demonstrates that for highly necessary information, the brain does not show the IS-related deactivation mode, but instead exhibits a reversed activation mode. The minimal neural representational similarity between concentration and IS conditions, as revealed by RSA (Figure 4F), further supports the distinction in neural patterns corresponding to high versus low subjective processing necessity.

Next, we examined objective necessity (Extended data1-Figure 2E,F). When processing necessity was reduced through the presentation of scrambled voice stimuli—creating conditions of objectively unnecessary information processing—we observed patterns mirroring those seen during IS with normal stimuli: high-to-low activation coupled with low-to-high deactivation (Figure 4E). The high representational similarity between normal and scrambled conditions, as revealed by RSA, suggests comparable neural states during active IS and when processing objectively absent information (Figure 4F). These findings were replicated in the visual domain, where active IS conditions demonstrated neural patterns similar to those observed during “close-eye” conditions, with high representational similarity between the two (Extended data1-Figure 2G). This convergence of evidence indicates that the brain engages in a deactivation mode when processing information deemed unnecessary, regardless of whether the reduced necessity is objectively or subjectively determined.

Taken together, our findings establish an intrinsic relationship between neural deactivation patterns and information processing necessity through bidirectional evidence. When we enhanced the neural deactivation pattern through DLPFC stimulation, participants demonstrated stronger IS effects, indicating a more decreased necessity for information processing. Conversely, when we manipulated processing necessity, we found that the brain constantly engages in a deactivation mode when processing unnecessary information, regardless of whether this reduced necessity stems from subjective strategy or objective absence of meaningful information. This convergent evidence from both neuromodulation and necessity manipulation firmly establishes the relationship between neural deactivation and reduced information processing.

## Discussion

This study addresses four fundamental questions regarding IS.

### 1. Can IS be considered a primary rather than subsidiary cognitive process?

Previous research has focused much more on information intake and efficiency enhancement but less on information attenuation and shielding. In the few relevant studies, IS was often viewed merely as a byproduct of other cognitive processes^25^. In contrast, our study validates IS as a primary cognitive process through the following aspects: **1) Conceptual distinction**: We clearly differentiate IS from related concepts such as selective attention, the cocktail party effect, and biased competition. Unlike these processes, which primarily focus on attending to relevant information while ignoring irrelevant inputs as a mean to enhance focus, IS is solely concerned with shielding information, irrespective of its relevance. This distinct functional goal— actively rejecting information rather than enhancing the reception of other stimuli—demonstrates IS’s role as a primary, instead of subsidiary, cognitive process; **2) Research framework**: We established a paradigm to directly study IS by integrating various potential shielding strategies, including those that solely shield B (inhibition, distancing) and those that shield B by focusing on A (distraction). Through conjunction analysis, we extracted the common features across these shielding strategies. This approach aims to uncover the core mechanisms of IS in general, moving beyond traditional methods that only consider indirect ignoring/inhibiting through selective focus or past expectation^20^. Therefore, this approach reveals that IS has its own complete operational mechanisms and can achieve its goals through multiple strategies, supporting its status as a primary process rather than a mere byproduct; **3) Cross-modal Validation**: Both auditory and visual IS show consistent phenomena and effectiveness, as evidenced by both subjective self-reports and objective neuroimaging measurements. This cross-modal consistency and effectiveness indicates that IS represents a universal cognitive process in its own right, rather than an incidental effect of other processes; **4) Mechanistic uniqueness**: Our findings show that IS involves predominantly deactivation-driven neural activities. The primary nature of the IS therefore lies in its distinct neural mechanism: while most cognitive processes primarily rely on the activation of specific brain regions to achieve their functions^26^, IS operates through the systematic deactivation of neural activities, representing a fundamentally different approach to information processing.

### 2. Can deactivation serve as a dominant neural mechanism?

The brain’s neural activity to specific demands manifests in three primary states: activation, inactivity, and deactivation^11–13^. While extensive research has examined activation and inactivation patterns, deactivation remains poorly understood and is typically viewed as a supportive mechanism—either for conserving neural resources or facilitating relevant information processing^14,16–19^. Using IS as a novel investigative lens, we provide direct evidence that deactivation serve as a primary mechanism in its own right.

Through neural investigations across auditory and visual IS tasks, we demonstrated that IS involves a deactivation-dominant neural dynamic, characterized by both brain activity patterns and neural coding capabilities. While initial frontoparietal activation and enhanced IS state decoding are observed in executive regions, these effects diminish over time. Concurrently, sensory regions show progressive deactivation and reduced IS decoding performance, becoming more pronounced particularly after the 3- second boundary. This temporal shift in both brain activity patterns and coding capabilities suggests a coordinated neural mechanism where initial executive control gives way to widespread deactivation in later stages of IS processing. In addition, more effective IS was linked to stronger deactivation dynamics and more pronounced reduction in decoding accuracy within sensory regions, suggesting that deactivation, especially its progressive increase over time, could be the key determinant of successful IS. More importantly, these patterns - including both the activation dynamics and neural coding characteristics - were observed consistently across both auditory and visual modalities, indicating a fundamental, domain-general mechanism rather than a modality-specific phenomenon.

The observed consistency in deactivation dynamics during IS underscores its potential as a fundamental neural strategy for efficient information management. It challenges the conventional understanding of deactivation as merely supportive^14,16–19^, instead positioning it as an independent and dominant process in the context of IS. This insight has broader implications for understanding inhibitory cognitive processes in general. It is reasonable to hypothesize that all inhibition-related cognitive processes might rely more on deactivation rather than on activation, highlighting the importance of this often-overlooked neural mechanism.

### 3. Is the neural representation underlying IS domain-general?

Through a series of cross-modal analyses, we provide evidence for the domain-general nature of IS. First, we identified a cross-modal transfer of the IS-related neural patterns: decoders trained on auditory IS could successfully identify visual IS, and vice versa, which was not observed during NP. This differential pattern indicates that IS induces a modality-independent neural state that is distinct from NP. While NP involves naturally receiving auditory or visual stimuli, leading to modality-specific brain activity that is challenging to generalize, IS engages executive control mechanisms to shield information, facilitating the generalization of neural patterns across modalities. In addition, the IS- increased cross-modal generalization became more pronounced during the late period (>3 seconds), suggesting the establishment of a stable, modality-independent shielding state.

Second, we found consistent patterns in how IS affects information feature processing across modalities. Using RSA, we demonstrated that IS consistently reduced the neural processing of emotional features (valence and arousal) across both auditory and visual domains. In contrast, non-emotional features (e.g., processing difficulty) showed modality-specific effects. This selectivity in cross-modal consistency is meaningful as emotional features typically are more attention-capturing and undergo preferential processing^27^. IS appears to employ a unified mechanism specifically targeted at disrupting highly salient information features.

Third, we identified specific regions showing synchronization between auditory and visual IS, which may explain the observed cross-modal consistency. Using ISC analysis, we found that while both IS and NP conditions exhibited synchronization peaks in primary sensory regions, IS uniquely engaged synchronization hot spots in executive control regions (DLPFC and PPC). These synchronized executive regions likely represent the neural substrate for a domain-general IS mechanism, with executive control systems orchestrating consistent shielding processes across different modalities.

### 4. Does the brain adopt a deactivation mode not only for IS, but for unnecessary information processing in general?

Through neuromodulation, we confirmed that this deactivation pattern causally contributes to the IS effect. Through necessity manipulation approaches, we further demonstrated that this deactivation mechanism extends beyond IS, representing a general neural strategy for handling unnecessary information processing.

First, our TMS-fMRI results demonstrated that deactivation plays a causal role in managing less-needed information. Specifically, excitatory rTMS over the left DLPFC enhanced IS performance and modulated the temporal dynamics of DLPFC activity, resulting in more pronounced transitions from high to low activation compared to the sham group. Moreover, individuals who exhibited stronger transitions from low to high deactivation in sensory regions also showed improved IS performance; the relation was prominent in TMS group. Therefore, we uncovered that the deactivate temporal dynamic in executive control and sensory regions are not a byproduct of IS; instead, this deactivation pattern intrinsically and causally contributes to the IS effect.

Second, by systematically manipulating the necessity for information processing, we revealed that the deactivation mode is not limited to IS but instead represents a general mechanism employed by the brain when processing is deemed unnecessary. This was observed regardless of whether the reduced necessity was subjectively induced (active IS) or objectively determined (absence of meaningful content). Notably, unlike the auditory scramble condition, which represents unnecessary information at the input level, the visual “eyes-closed” condition created a shielding scenario representing complete unnecessity at the reception level. Despite these different forms of unnecessity, both conditions led to similar deactivation patterns, suggesting that whenever unnecessity is present, the brain engages in IS-like neural activity. In contrast, when information was deemed highly necessary (as in the concentration condition), the brain adopted a reversed activation pattern instead of deactivation.

When faced with unwanted information, the brain could theoretically employ two strategies: (1) Investing cognitive resources to activate specific neural mechanisms that inhibit the reception of information; or (2) Reducing resource investment, thereby diminishing the processing of that information. Our research has yielded a surprising discovery: when shielding or facing unneeded information, the brain predominantly employs a deactivation pattern. This phenomenon is not unique to our study but has been observed in other contexts, such as repetition suppression in animal and human studies^28,29^ and memory suppression in human research^30^, where similar deactivation responses were found in relation to reduced processing demands. These findings suggest that deactivation is a widespread mechanism for minimizing information processing—a phenomenon that has not been given due attention or systematically summarized until now.

Deactivation, however, is entirely reasonable as it implies a reduction in cognitive resource allocation^16–19^, which aligns with decreased information processing demands and proves more energetically efficient and evolutionarily advantageous^31^. This also suggests that, in addition to the well-recognized mechanism of activating specific brain regions and pathways to fulfill certain functions^32^, deactivation also represents a crucial mode of information processing—one that has been largely overlooked. Our findings could serve as a starting point for further research into the essential role of deactivation in cognitive function, highlighting that the brain’s ability to “turn down” processing may be as fundamental as its capacity to “turn up” activation in managing information processing demands.

## Methods

Experiments 1-4 were approved by the Ethical Committee at the School of Psychology, Shenzhen University (Approval No. SZU_PSY_2024_067).

### Participants

In our study, participants self-identified their ethnicity as East Asian Chinese, which was subsequently verified against their government-issued identity cards for accuracy. A total of 218 participants were enrolled across four experiments. Specifically, Experiment 1 involved fMRI assessments and was subdivided into two parts: AIS (n = 35) and VIS (n = 28). Experiment 2 involved EEG assessments, further subdivided into three parts: 2.1 (n = 30), 2.2 (n = 40), and 2.3 (n = 40). Experiment 3 involved combined TMS and fMRI assessments (n = 45), with participants equally assigned to three TMS groups. Experiment 4 utilized fMRI and included participants from Experiment 1 who agreed to participate again. Specifically, the AIS experiment involved 15 participants from Experiment 1-AIS, while the VIS experiment included 27 participants from Experiment 1-VIS. All participants provided informed written consent, and the study procedures were approved by the Shenzhen University Institutional Review Board. Nine participants were lost to attrition (four from Experiment 1, two from Experiment 2, and three from Experiment 3), and an additional five participants from Experiment 3 voluntarily withdrew before completing the study. Data from the remaining 205 participants were analyzed: Experiment 1 (AIS, n = 32; VIS, n = 27), Experiment 2 (2.1: n = 30, 2.2: n = 40, 2.3: n = 38), Experiment 3 (n = 37, with 13, 12, and 12 participants in the left, right, and sham TMS groups, respectively), and Experiment 4 (AIS: n = 15, revisited participants from Experiment 1-AIS; VIS: n = 27, revisited participants from Experiment 1-VIS). Participants’ demographic and neuropsychological data are summarized in Table 1. All participants provided written informed consent and were monetarily compensated for their participation.

**Table 1.** Demographic information of participants in the final analysis (*M* ± *SD*).

| Experiment 1 | AIS (n = 32) | VIS (n = 27) |  |  |  |
| --- | --- | --- | --- | --- | --- |
| Age (years) | 21.4 ± 1.9 | 21.04 ± 2.46 | | | |
| Sex<br>(male/female) | 19/13 | 14/13 |  |  |  |
| Experiment 2 | Exp 2.1<br>(n = 30) | Exp 2.2<br>(n = 40) | Exp 2.3<br>(n = 38) |  |  |
| Age (years) | 20.27 ± 1.39 | 20.38 ± 1.7 | 21.55 ± 2.46 | | |
| Sex<br>(male/female) | 15/15 | 23/17 | 19/19 |  |  |
| Experiment 3 | Sham group<br>(n = 12) | rDLPFC-activated group<br>(n = 12) | IDL PFC-activated group<br>(n = 13) | Statistic | *p* value |
| Age (years) | 20.73 ± 2.22 | 20.58 ± 2.11 | 20.08 ± 1.75 | 0.35 (*F*) | 0.705 |
| Sex<br>(male/female) | 7/5 | 6/6 | 9/6 | 0.06 ($\chi^2$) | 0.972 |
| **Experiment 4** | **AIS (n = 15,**<br>revisited<br>participants from<br>Experiment 1) | **VIS (n = 27,**<br>revisited<br>participants from<br>Experiment 1) |  |  |  |
| Age (years) | 21.4 ± 2.13 | 21.04 ± 2.46 | | | |
| Sex<br>(male/female) | 8/7 | 14/13 |  |  |  |

### Experimental materials

The AIS experimental materials comprised a total of 400 human voice clips, divided into two sets. The normal set consisted of 300 voice clips, with 100 clips each of 3- second, 6-second, and 10-second durations. The scrambled set was generated by converting each of the 100 normal 6-second voice clips into scrambled counterparts, resulting in an additional 100 scrambled voice clips.

The original texts were in Chinese, and they were translated into English for readability. The normal voice clips were generated through the following process: First, ChatGPT-4 generated a sentence allowed for a natural speaking speed of 6 seconds (e.g., “This is a very complex problem. It is difficult to find a solution. It requires professional knowledge and skills”). Based on this 6-second sentence, GPT then created a shorter sentence with fewer information (3 second speaking speed allowed; e.g., “This problem is too difficult to solve”), and a longer sentence with more information (10 second speaking speed allowed; e.g., “This is a very complex problem, difficult to find a solution, requiring professional knowledge and skills. Many people have tried, but there is still no progress, needing more time and resources”). Second, the voice clips were recorded by AI speakers (half male, half female; generated from https://ttsmaker.cn) reading aloud the short or long sentences, at a normal speaking speed, volume, and pitch. Third, for each of the 6-second and the corresponding short and long sentences, GPT-4 generated a multiple-choice quiz with 4 options (e.g., “What did this voice say? A: There might be problems in the future; B: This is an easy problem.; C: This is a complex problem; D: The problem has been solved”). Only one option (in this case, option C) correctly matched the information conveyed in the corresponding voice clip. These quizzes were designed to assess participants’ understanding of the spoken content. The scrambled voice clips were created by taking the 100 normal 6-second voice clips and thoroughly scrambling the audio, rendering them meaningless.

The experimental materials for the VIS comprised 100 images randomly selected from the International Affective Picture System (IAPS^33^). Both the normal voice clips and the IAPS pictures were randomly assigned to four different experimental conditions: 1) NP - Participants listened to the voice clips or viewed the pictures in a natural manner, without any additional instructions or interventions; 2) Inhibition - Participants were instructed to inhibit or suppress their natural responses while listening to the voice clips or viewing the pictures; 3) Distraction - Participants were asked to perform a secondary task (e.g., imagining their home) while listening to the voice clips or viewing the pictures; 4) Distancing - participants were instructed to mentally distance themselves from the content of the voice clips or pictures, viewing them as if they were not personally involved. Detailed instruction texts for each condition can be found in Extended Data 3.

Balancing randomization and analytical uniformity, we selected an equal number of stimuli for each of the four experimental conditions, maintaining consistency across subsequent experiments. Specifically, each condition included an equal number (n = 25) of either IAPS images or voice clips. The voice clips were 6 seconds in duration or their corresponding 3- or 10-second versions. To assess potential confounds, we evaluated whether emotional valence, arousal, information load, and processing difficulty differed significantly between the assigned conditions. An independent sample, demographically matched to the main study participants, rated the stimuli using nine-point scales: valence (1 = most negative to 9 = most positive), arousal (1 = least arousing to 9 = most arousing), information load (1 = lowest to 9 = highest), and processing difficulty (1 = least difficult to 9 = most difficult). The results indicated that the stimuli were well-matched across conditions, minimizing potential confounds related to emotional content, information density, or processing difficulty. Comprehensive supplementary materials are provided in Extended Data 3, including the full text of all voice clips, the IAPS IDs of the pictures, the associated quizzes, descriptive statistics for average ratings and individual stimulus ratings, and all statistical results to ensure transparency and facilitate replication efforts.

For the EEG experiment, voice clips were presented via headphones (Sennheiser HD200 PRO) at a comfortable level (80–85 dB SPL). For the fMRI experiments, voice clips were presented using Media Control Functions (DigiVox, Montreal, Canada) via electrostatic headphones (NordicNeuroLab, Norway, or Sensimetrics, USA) at a comfortable level (80–85 dB SPL).

### Behavioral task

Prior to participating in the AIS/VIS experiment, participants received thorough briefings on the detailed instructions associated with each instruction trigger word: “NP”, “inhibition”, “distraction”, or “distancing”. Participants were trained to associate each trigger word with its corresponding cognitive strategy.

The experimental paradigm, illustrated in Figure 1A, consisted of the following sequence for each trial: (1) Presentation of the instruction trigger word (“NP”, “inhibition”, “distraction”, or “distancing”) for 1.5 seconds; (2) A fixation period lasting either 1.5 or 3 seconds; (3) Voice or picture presentation, during which participants applied the strategy indicated by the trigger word for 6 seconds; (4) A “sensory” question, either “Did you hear this voice? ” for the AIS experiment or “Did you see this picture?” for the VIS experiment, presented for 1.5 seconds; (5) An “understand” question, either “Did you understand the meaning of the voice?” for the AIS experiment or “Did you understand the meaning of the picture? ” for the VIS experiment, also presented for 1.5 seconds; (6) An “objective” question, either “What information did this voice convey? ” for the AIS experiment or “What information did this picture convey? ” for the VIS experiment, presented with four possible options, including one correct answer, for 4.5 seconds.

### Experimental Procedure

We conducted four experiments to investigate the neural mechanisms underlying IS. In Experiment 1, we utilized fMRI to assess the neural correlates of AIS and VIS, employing distinct participant samples for each modality. Experiment 2 employed EEG to capture the temporal neural dynamics associated with AIS. This experiment comprised one main study (2.1), which utilized 6-second duration voice stimuli, alongside two complementary studies (2.2 and 2.3), each using 3-second or 10-second voice stimuli, with different participant samples for each. In Experiment 3, we employed TMS combined with fMRI to explore brain-behavior causality related to IS. Finally, Experiment 4 utilized fMRI to verify the specificity of IS by comparing its brain activity patterns with those observed during “concentration” and “complete shielding” conditions. While Experiments 1, 2, and 3 each utilized different participant samples, Experiment 4 involved a subset of participants from Experiment 2 who returned for further testing.

### Experiment 1 (fMRI)

Both the AIS and VIS experiments followed a within-subject, event-related design with four task conditions (NP, inhibition, distraction, and distancing). Each voice clip or picture was repeated twice to ensure sufficient trials (n = 50) for each task condition.

### Experiment 2 (EEG)

Experiment 2 was the AIS experiment. The experimental design for this study was identical to that of Experiment 1. Experiment 2.1 used 6-second voice stimuli, while the complementary Experiments 2.2 and 2.3 employed 3-second and 10-second voice stimuli, respectively, to replicate the findings across varying stimulus lengths.

### Experiment 3 (TMS + fMRI)

Experiment 3 was the AIS experiment. The experimental design, voice stimuli, and timing for each screen in this study were identical to those used in Experiment 1. In this study, participants were divided into three TMS groups, resulting in a mixed-design experiment with a 4 (*task condition*: NP, inhibition, distraction, distancing) × 3 (*TMS group*: left DLPFC-activated, right DLPFC-activated, sham) structure. The *task condition* was a within-subject factor, while the *TMS group* was a between-subject factor.

The experimental procedure is illustrated in Extended Data1-Figure 2A. Participants underwent four TMS sessions, each followed by a task session (fMRI session). The duration of each TMS session was set to 15 minutes to ensure that the after-effects of TMS fully covered the subsequent fMRI task session (14.38 min)^34,35^. We used offline TMS, rather than concurrent TMS-fMRI, to avoid potential technical issues^36^ and minimize any side effects (e.g., muscle twitching) that could impact participants’ task performance.

### TMS protocol

A figure-eight-shaped coil was connected to the magnetic stimulator (M-100 Ultimate; Shenzhen Yingchi Technology Co., Ltd, Shenzhen, China). The TMS targets were the lDLPFC and rDLPFC for the two experimental groups, and the vertex to provide a similar scalp sensation in the sham TMS group^37^. To locate the lDLPFC and rDLPFC, we used the MNI coordinates (x = −28, y = 0, z = 58 for lDLPFC, and x = 48, y = 0, z = 52 for rDLPFC), which were derived from activation peaks identified in the contrast between the conjunction of three IS strategies and NP in Experiment 1. To locate the vertex, the coordinate (x = 0, y = 0, z = 80) was determined as the midpoint of a region halfway between the nasion and the inion, and equidistant from the left and right ear^38^. To locate the motor area used for the measurement of the resting motor threshold (rMT), we determined the coordinate of the left (x = −38.3, y = −15.2, z = 67.9) or right motor hand area (x = 38.3, y = −15.2, z = 67.9) which was found to be the optimal coil position for motor cortex stimulation^39^. Prior to stimulation, a T1- weighted structural MRI was obtained from each participant and normalized to the MNI space. The voxels corresponding to the location of the right VLPFC or the vertex was marked on the participant’s normalized T1 images. The location of TMS coil was guided by a frameless stereotactic neuronavigation system (Brainsight, Rogue Research, Montreal, Canada). During stimulation, the position and orientation of the coil was monitored continuously by the neuronavigation system.

Each participant’s resting motor threshold (rMT) was measured from their motor cortex. Three electrodes were fixed on their right palm to collect motor evoked potentials (MEPs). The rMT was defined as the lowest intensity evoking at least five MEP responses with amplitudes larger than 50 μV in 10 trials. The rTMS was applied at 90% of each participant’s rMT during the experiment^40^. The rTMS was administered at 10 Hz; this frequency could produce an excitatory effect on the target brain region according to the majority of TMS literature^41^. Each 15 min session contained 30 trains, each lasting 3.9 s with an inter-train interval of 26.1 s^42^. The TMS simulated electric field is illustrated on an adult brain model in Figure 4A (SimNIBS software, www.simnibs.org). To reduce the potential effect of body movement (especially walking) on TMS-induced neural plasticity, participants sat beside the scanner bed when receiving TMS pluses. They moved slowly from their chair to the MRI scanner bed immediately after each TMS session, and moved slowly from the scanner bed to the chair to receive TMS stimuli after the first MRI session. The time elapsed between the TMS and MRI sessions was 36.1 ± 13.7 s.

### Experiment 4 (fMRI)

#### AIS

A total of 15 participants from Experiment 1-AIS revisited the experiment one week later. The task procedure and experimental paradigm were identical to those of Experiment 1, with the following exceptions: First, this experiment utilized 25 normal voice stimuli, randomly selected from the original 100 voice clips, along with 100 scrambled voice stimuli. Second, since participants had previously engaged in shielding of normal voices in Experiment 1, this supplementary experiment instructed them either to concentrate on the normal voice (representing maximal attention) or to employ shielding strategies while processing the scrambled voice (representing minimal attention or near-complete shielding). Third, trials with scrambled voices included only the “sensory” and “understand” questions. Although the “understand” question for scrambled stimuli may seem unnecessary, it serves a technical purpose in confirming that the scrambled voices convey no meaningful information, thereby validating the intended function of the stimuli.

The design of this experiment allowed for a within-subject comparison between participants’ first-visit conditions (NP-normal, inhibition-normal, distraction-normal, distancing-normal) and their second-visit conditions (concentration-normal, NP- scramble, inhibition-scramble, distraction-scramble, distancing-scramble), resulting in a 9-task condition within-subject design.

#### VIS

A total of 27 participants from Experiment 1-VIS also revisited the experiment one week later. The task procedure remained identical to Experiment 1, with the following modifications: First, this experiment utilized 25 IAPS pictures, randomly selected from the original set of 100. Second, since participants had previously engaged in shielding of pictures in Experiment 1, this supplementary experiment required them to close their eyes during the picture presentation period.

This design allowed for a within-subject comparison of participants’ first-visit conditions (NP, inhibition, distraction, distancing) and their second-visit condition (close-eye), resulting in a 5-task condition within-subject design.

### Data recording and analysis

### Behavioral data

Behavioral data were recorded using E-prime 3.0 (PST, Pittsburgh, USA). Data were analyzed using R Statistical Software (v4.3.2). Descriptive data are presented as *M* ± *SD*, unless otherwise mentioned. Linear mixed-effect (logistic) models were constructed to examine trial-by-trial behavior. In the basic model, the within-subject predictor “strategy type” had two levels: trials in which participants employed one of the “distancing,” “distraction,” or “inhibition” strategies were labeled as IS, whereas trials in which participants processed to the stimuli naturally were labeled as NP. The NP condition served as the baseline. For each trial, responses to the “sensory,” “understanding,” and accuracy on “objective” questions were coded as 0 (negative response—did not hear/see, did not understand, or incorrect) or 1 (positive response— heard/saw, understood, or correct) and used as response variables in separate models.

Participants’ responses to the subjective “sensory” question were converted into the sensory rate (*P*_sensed_), representing the probability of having heard/saw the voice/picture stimulus. Similarly, responses to the subjective “understand” question were converted into the understanding rate (*P*_understood_), indicating the probability of having understood the voice stimulus. Participants’ performance on the objective multiple-choice quiz was converted into accuracy (*ACC*_objective_), representing the proportion of correct answers. Random intercepts were included for each participant to account for individual differences. The main basic models are as follows:

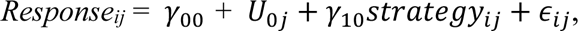

where:

*Response_ij_* is response in the *i*-th trial of the *j*-th individual participant;

*γ*_00_ is the fixed intercept at the reference level, i.e., in NP;

*γ*_10_ is the fixed effect coefficient, i.e., difference between IS and NP;

*U*_0*j*_ is the random intercept for the *j*-th participant;

*ɛ_ij_* is the residual.

The basic model was applied to each experiment. Particularly, in Experiment 3, we utilized TMS groups (lDLPFC-activated, rDLPFC-activated, and sham) as an additional between-subject predictor to probe difference in the effectiveness of IS strategies across different TMS manipulations, with the sham group as the reference condition. Cross-level interactions between TMS groups and strategy types were also included as a fixed effect.

We included participants who completed the study twice — that is, those who participated first in Experiment 1 and then in Experiment 4—to explore factors across different visits. We constructed three models: 1) The “concentration” model followed a 3 (*task condition*: concentration, NP, IS) within-subject design. We used NP as the baseline to examine whether the concentration and the IS condition differed from NP when participants were presented with normal-ordered stimuli; 2) The “scramble” model followed a 2 (*task condition*: NP, IS) × 2 (*stimuli type*: normal order, scrambled order) within-subject design. Here, we used NP as the baseline for *task condition* and normal order as the baseline for *stimuli type* to investigate differences in performance across *task condition* and *stimuli type* combinations.

For the “scramble” model, we included both *P*_sensed_ and *P*_understood_ as response variables. Additionally, we conducted a manipulation check to verify whether the scrambled stimuli effectively conveyed meaningless information. In this check, *P*_understood_ served as the response variable in a model that included stimulus type (normal order vs. scrambled order) as a fixed effect. We considered the stimulus manipulation successful if participants exhibited significantly lower *P*_understood_ for scrambled voices compared to normal voices.

### EEG data

EEG data were recorded using a 64-channel amplifier (BrainProducts, Gilching, Germany), with a sampling frequency of 250 Hz. Electrode impedances were kept below 10 kV. The reference electrode was placed at the CPz. No online filter was applied.

#### Preprocessing

Data analysis was performed in MATLAB using the EEGLAB toolbox (https://sccn.ucsd.edu/eeglab/index.php). Data were first re-referenced to the average of the left and right mastoids. Ocular artifacts were eliminated using the independent component analysis. Then, the EEG data were filtered using a 0.1 to 30 Hz bandpass filter. The filtered data were segmented beginning 200 ms before the onset of the voice stimuli and lasting for 800 ms. The baseline correction was based on the -200-0 ms pre-stimulus onset time window.

#### MVPA & Temporal decoding analysis

Data analysis was performed using MVPAlab toolbox^43^. The preprocessed EEG voltage (including filtering, baseline correction, and artifact removal) was extracted as the feature of interest. To balance and stabilize the dataset, we averaged every five trials. All electrodes were selected for the analysis, and data normalization was conducted across trials to minimize systemic biases between different experimental sessions. The analysis timing was set from -200 to 3000 ms, -200 to 6000 ms, -200 to 10000 ms relative to the stimulus onset, with a step size of 5 timepoints to provide a detailed temporal resolution.

MVPA was used to investigate the temporal dynamics of auditory processing under three different conditions: 3-second, 6-second, and 10-second voice stimuli. The MVPA analysis was conducted using the NP condition as the training set, and we examined the decoding success rate for the IS condition. A higher decoding rate (indicated by red regions) implies greater dissimilarity between the two conditions. We used a 5-fold cross-validation approach to train and test the support vector machine (SVM) with a linear kernel classifier to discriminate between two conditions. The model’s performance was thoroughly evaluated by calculating the mean accuracy, area under curve, and weights vector.

After that, we performed decoding analysis on time-resolved broadband responses across channels to discriminate the two conditions (IS and NP) at each time from -200 ms to 3,000 ms, -200 ms to 6,000 ms, -200 ms to 10,000 ms, separately for each condition. The obtained decoding time series for each condition were smoothed by the moving average algorithm (5 time points).

#### Change-point detection

We use Matlab’s built-in changepoints detection algorithm (findchangepts) to detect at which specific point in time the decoding rate curve mutates.

### MRI data

Brain images were collected using a 3T MR scanner (Siemens Trio). Functional images were collected using an EPI sequence (number of slices, 72; gap, 0.6 mm; slice thickness, 2.0 mm; TR, 1500ms; TE, 30ms; flip angle, 75°; voxel size, 2 mm × 2 mm × 2 mm; FOV, 192 mm × 192 mm). Structural images were acquired through 3D sagittal T1-weighted MPRAGE (224 slices; TR, 1900ms; TE, 2.23ms; voxel size, 1.1 mm × 1.1 mm × 1.1 mm; flip angle, 8°; inversion time, 904 ms; FOV, 220 mm × 220 mm).

Preprocessing, subject-level, group-level, and the temporal analysis of brain activity were applied to Experiment 1, 3-4. MVPA were applied to Experiment 1 and 3. RSA was applied to Experiment 1 and 4. All fMRI analysis were conducted on the pre- defined ROIs, unless otherwise mentioned.

#### Regions of interest definition

Given that IS involves top-down processes wherein higher-order cognitive regions exert control over sensory regions^44^, we focus on the DLPFC and PPC, which together constitute the executive control network^45^. Additionally, we included sensory regions specific to each modality—namely, the STC for auditory processing^46,47^ and the visual cortex (VC, including V1-V3) for visual processing^48^. Increased activation within the executive control network would thus serve as an indicator of the implementation and execution of IS strategies. Conversely, a decrease in STC or VC activation would indicate IS engagement, based on traditional models and empirical studies of attention suggesting that activations in stimulus-specific regions are reduced when participants ignore stimuli^49,50^. All ROIs were selected based on the anatomical definitions provided by the Automated Anatomical Labeling atlas 3 (AAL3)^51^. Specifically, the DLPFC was defined as Brodmann areas (BA) 9 and 46^52^; the PPC as BA 5 and 7^53^; the STC as BA 22^54^; and the VC as BA 17-19^55,56^. These ROIs were defined in MNI space, combined into a unified ROI mask specific to each experiment: for the AIS experiment, the ROIs included the DLPFC, PPC, and STC; for the VIS experiment, the ROIs comprised the DLPFC, PPC, and VC. These unified ROI masks were then transformed into individual participant space using the inverse normalization parameters estimated during preprocessing.

#### Preprocessing and subject-level analyses

Images were preprocessed and analyzed using Matlab (2020b) and Statistical Parametric Mapping (SPM12; Wellcome Trust Centre for Neuroimaging, London, UK). The first ten volumes were discarded to account for signal equilibration and participants’ adaptation to scanning noise. Functional data were first corrected for geometric distortion with the SPM FieldMap toolbox, and slice-time corrected and realigned for motion correction by registration to the mean image. Artifact detection was conducted using the Artifact Detection Tools (ART) software (https://www.nitrc.org/projects/artifact_detect); global mean intensity (> 2 standard deviations from mean image intensity for the entire scan) and motion (> 2 mm) outliers were identified and entered as regressors of no interest in the first-level general linear model (GLM)^57^. Then, functional images were co-registered with the T1-weighted 3D images, normalized to MNI space, and smoothed with an 8-mm full width at half-maximum isotropic Gaussian kernel. We regressed out six motion estimates (three translations and three rotations), one artifact (outlier scans) as identified by ART, and two physiological time series (cerebrospinal fluid and the white matter signals) with global signal regression For subject-level analyses, we specified GLMs with four regressors for the four conditions (NP, inhibition, distraction, and distancing). The duration for each regressor was defined as the voice/picture exposure period (6 s) per trial. All regressors were convolved with the canonical hemodynamic response function in SPM12. Each normalized image was then high-pass filtered using a cutoff time constant of 128 s. natural hearing, inhibition, distraction, and distancing. For each participant in Experiment 1 and 3, we generated seven contrast images: (1) NP, (2) inhibition, (3) distraction, (4) distancing, (5) IS, (6) IS > NP, and (7) IS < NP. Contrasts 1-4 were calculated against the implicit baseline as defined by SPM. To identify brain regions commonly activated across all IS strategies, we performed a conjunction analysis of inhibition, distraction, and distancing conditions, producing the contrast 5 (IS). Contrast 6 represented IS-related activation, obtained by the conjunction analysis of (inhibition > NP), (distraction > NP), and (distancing > NP). Conversely, contrast 7 represented IS-related deactivation, derived from a conjunction analysis of (NP > inhibition), (NP > distraction), and (NP > distancing).

For each participant in Experiment 4-AIS, besides the first-visit contrast images (the above seven), we generated ten additional contrast images: (1) concentration, (2) concentration > NP, (3) concentration < NP, (4) NP-scramble, (5) inhibition-scramble, (6) distraction-scramble, (7) distancing-scramble, (8) IS-scramble, (9) IS > NP- scramble, and (10) IS < NP-scramble. For each participant in Experiment 4-VIS, we generated one additional contrast image: (1) close-eye.

These subject-level contrast images were subsequently used as inputs for second-level random-effects analyses to assess group-level effects.

#### Group-level activity analysis

In Experiment 1, we performed one-sample *t*-tests on the IS > NP and IS < NP contrasts to detect group-level IS-related brain activation and deactivation. In Experiment 3, we performed two-sample *t*-tests on these same contrasts across the lDLPFC-activated group versus the sham group, as well as the rDLPFC-activated group versus the sham group. These analyses aimed to determine whether TMS over the left or right DLPFC differentially influences IS-related brain activity.

In Experiment 4, we developed two models: the “concentration model” incorporated the first-visit conditions (IS and NP) and the second-visit condition (concentration). We conducted one-sample *t*-tests on the IS > NP, IS < NP, concentration > NP, concentration < NP contrasts respectively, to detect group-level IS-related or concentration-related brain activation and deactivation, and pair-sample *t*-tests to compared the activation and deactivation differences between IS and concentration. The “scramble model” combined the first-visit conditions (IS-normal and NP-normal) with the second-visit conditions (IS-scramble and NP-scramble). We conducted one-sample *t*-tests on the IS > NP-normal, IS < NP-normal, IS > NP-scramble, IS < NP-scramble contrasts respectively, to detect group-level “active IS”-related or “complete IS”-related brain activation and deactivation, and pair-sample *t*-tests to compared the activation and deactivation differences between active IS and complete IS.

#### Change-point detection

We constructed temporal curves of IS-related brain activation differences by calculating the difference between the IS and NP conditions at each TR (0–1.5 s, 1.5–3 s, 3–4.5 s, 4.5–6 s) over the entire 6-second stimulus exposure period. Specifically, we extracted the time series data for both the IS and NP conditions and computed the difference (IS minus NP) at each TR. This provided a temporal profile of activation differences associated with IS relative to NP.

We also constructed temporal curves of IS-related decoding rates at each TR. Specifically, we performed MVPA at each TR to classify between IS and NP conditions. Details of the MVPA procedures can be found in the section “MVPA—Unimodal MVPA.” This allowed us to construct temporal curves of IS-related decoding accuracies over the stimulus exposure period.

Both types of temporal curves—the activation differences and the decoding accuracies—were constructed using combined ROI masks. Focusing on these ROIs allowed us to target regions implicated in executive control and sensory processing relevant to IS.

To identify specific time points at which significant changes occurred in the temporal curves, we employed MATLAB’s built-in changepoint detection algorithm (findchangepts). This algorithm detects points where the statistical properties of a sequence change significantly, enabling us to pinpoint when sudden shifts in decoding rates or activation differences occurred. To assess the reliability and stability of the changepoint detection, we conducted 1,000 bootstrap analyses. In each bootstrap iteration, we sampled the data with replacement and repeated the changepoint detection process. This approach allowed us to obtain a distribution of changepoint times and calculate the 95% confidence interval, providing statistical confidence in the timing of the detected changepoints.

#### Activity temporal analysis

To investigate potential temporal dissociations in brain activity as a result of the IS manipulation, GLM analyses were conducted to compare neural responses across two time frames for the voice/picture stimuli. Specifically, the models examined brain activity during the 0-3 s, and 3-6 s time periods following the onset of the 6-second voice/picture. For both IS > NP and NP > IS contrasts, Experiment 1 performed one- sample *t*-tests on the 0-3s and 3-6s respectively, and a pair-sample *t*-test to compared the temporal difference. Experiment 3 used a 3 (*TMS groups*: lDLPFC-activated, rDLPFC-activated, and sham) × 2 (*time*: pre-3s, post-3s) flexible factorial design to test whether TMS may modulate the IS-activation and deactivation across different time periods. In Experiment 4, the “concentration model” used a 3 (*task conditions*: concentration, IS, and NP) × 2 (*time*: pre-3s, post-3s) flexible factorial design to test whether concentration and IS may differ in the activation and deactivation across different time periods. The “scramble model” used a 2 (*task conditions*: IS, NP) × 2 (*time*: pre-3s, post-3s) × 2 (*voice type*: normal, scramble) flexible factorial design to test IS-normal and IS-scramble may differ in the activation and deactivation across different time periods.

#### Beta value extraction and visualization

For the brain activity analysis, a voxel-wise analysis was conducted within the unify ROI, and the surviving voxels (after performing false discovery rate (FDR) correction across all the voxels within the ROI) are displayed on the brain activation maps. To further illustrate the results, we extracted the averaged BOLD signals (parameter estimates) from all voxels within each individual ROI for each participant using the MarsBaR toolbox (https://marsbar-toolbox.github.io). These averaged parameter estimates were then subjected to *post-hoc* statistical tests and visualization.

#### MVPA

The analysis was conducted using the CoSMoMVPA toolbox (https://www.cosmomvpa.org) and LIBSVM (https://www.csie.ntu.edu.tw/~cjlin/libsvm). The analysis included three components: 1) a **unimodal decoding analysis** to assess the discriminability of brain activity patterns between IS and NP conditions within the same sensory modality, 2) a **feature decoding analysis** to assess how IS affects the discriminability of information features (high versus low valence, arousal, information load, and processing difficulty) in neural activity patterns, and 3) a **cross-modal decoding analysis** to test whether IS-related neural patterns generalize across auditory and visual modalities.

### Data preparation

Following data preprocessing and GLM analysis, we generated beta values corresponding to each condition (IS or NP) across three time periods: 0-6 seconds, 0-3 seconds, and 3-6 seconds post-stimulus onset. However, the initially generated beta values were not suitable for MVPA, which requires trial-specific data. Therefore, we restored beta values for each single trial (https://github.com/ritcheym/fmri_misc/blob/master/generate_spm_singletrial.m). These restored single-trial beta values were then used in the following MVPA analyses.

### Unimodal MVPA

We evaluated the model’s ability to discriminate between IS and NP conditions within each modality (using data from Experiment 1’s AIS and VIS tasks, and Experiment 3’s AIS task). Higher decoding accuracy indicates more distinct neural representations compared to the chance level (0.5). We focused on the previously defined ROIs (DLPFC, PPC, STC/VC) and used the beta values across all voxels within each ROI to decode between NP and IS. A leave-one-run-out cross-validation procedure was employed to validate the classifier’s ability to distinguish between conditions. In each classification iteration, the training set consisted of data from all runs except one (40 trials: 20 trials per condition), and the testing set included data from the left-out run (10 trials: 5 trials per condition). To reduce variability and enhance the reliability of the results, we performed 100 bootstrap iterations with random resampling of trials. Principal component analysis (PCA) was applied to the training data for dimensionality reduction before training the SVM classifier.

### Feature MVPA

We evaluated the model’s ability to discriminate among the following four conditions: high and low intensity of information features under the IS state, and high and low intensity of information features under the NP state (using data from Experiment 1’s VIS task). The analyzed features included valence, arousal, information load, and processing difficulty, which were derived from ratings by an independent, demographically-matched sample (see “Methods - Experimental materials” for details). To ensure balanced trial numbers across conditions, features were binarized based on median splits: ratings above the median were classified as high intensity, while those below were classified as low intensity. Higher decoding accuracy indicates more distinct neural representations compared to the chance level (0.25). To facilitate holistic comparisons across different conditions, we focused on the combined ROI, which included the DLPFC, PPC, and VC. Beta values from all voxels within this ROI were used to decode among the four conditions. A leave-one-run-out cross-validation procedure was employed to validate the classifier’s ability to distinguish between conditions. In each classification iteration, the training set consisted of data from all runs except one (16 trials: 8 trials per condition), and the testing set included data from the left-out run (4 trials: 2 trials per condition). The bootstrap iterations and PCA settings were identical to those used in the unimodal MVPA.

### Cross-modal MVPA

We evaluated the model’s ability to generalize the IS or NP conditions across modalities (Experiment 1, AIS and VIS experiments), with higher accuracy indicating greater generalization of neural representations between modalities. To facilitate cross-modal analysis, we defined a CEN ROI by combining the regions (DLPFC and PPC) that were consistently involved in both AIS and VIS.

The cross-modal MVPA analyses were conducted under two configurations: **1) auditory modality as training set, visual modality as testing set, and 2) visual modality as training set, auditory modality as testing set**. Unlike the unimodal MVPA using the leave-one-run-out method, for the cross-modal MVPA, the inherent cross-modal testing procedure served as validation. The selection of visual trials matched the sample sizes of the auditory dataset and ensured a balanced number of trials for both NP and IS conditions. The bootstrap iteration settings were identical to those used in the unimodal MVPA. Features extracted via PCA from one modality training data to extract shared, modality-invariant features, which were then applied to another modality testing data for classification.

### Statistical Analysis

For both unimodal and cross-modal MVPA, one-sample *t*-tests were conducted to determine whether the classification accuracy for each condition significantly exceeded the chance level (0.5). For the feature MVPA, one-sample t-tests were conducted to determine whether the classification accuracy for each condition significantly exceeded the chance level (0.25). Paired- or two-sample *t*-tests were performed to compare the accuracy differences between the IS and NP conditions within the same modality (unimodal MVPA) or between the two modalities (cross-modal MVPA). Effect sizes for the *t*-tests were calculated using *Cohen’s d*. For the feature MVPA, a repeated-measures ANOVA with *task condition* (NP, IS) and *feature intensity* (high, low) as within-subject factors was used to determine whether IS affects the discriminability of information features. Effect sizes were calculated using *η²*.

### RSA

The analysis was performed using MATLAB, SPM, The Decoding Toolbox (https://sites.google.com/site/tdtdecodingtoolbox), and custom scripts. It included two main components: a **neural-neural RSA** to assess the similarity of brain representations between different experimental conditions, and a **feature-neural RSA** to examined whether the brain’s neural activity patterns represent specific characteristics inherent in the stimuli. To obtain a holistic representation and facilitate comparisons across different conditions, RSA was performed on a combined ROI (AIS: DLPFC, PPC, STC; VIS: DLPFC, PPC, VC).

### Data Preparation

The analysis used the single-trial beta values corresponding to each condition (e.g., IS or NP). We computed pairwise Pearson correlation coefficients between all single-trial beta values of the ROI across the relevant conditions, resulting in a representational dissimilarity matrix (RDM) for each participant. It reflects the dissimilarity of neural patterns between trials, where lower correlation indicates greater dissimilarity. To achieve a group-level representation, we averaged the RDMs across all participants, producing an average RDM. We then compared this average RDM with a conceptual model RDM—an idealized binary classification model with values of 0 and 1 representing expected dissimilarities between conditions—using Spearman’s rank correlation coefficient. This comparison assessed how well the neural patterns aligned with the predicted patterns based on the conceptual model, thereby evaluating the ROI’s ability to distinguish between NP and IS conditions. A permutation test with 5,000 iterations was conducted to determine the statistical significance of these correlations. Finally, we transformed the RDMs into representational similarity matrices (RSMs) by subtracting each value in the RDM from 1 (i.e., RSM=1−RDM). This transformation converts dissimilarity measures into similarity measures, facilitating direct comparison of neural patterns.

### Neural-neural RSA

The analysis was performed across three time windows: 0-6 seconds, 0-3 seconds, and 3-6 seconds post-stimulus onset. For the “Concentration” model, we generated RSMs for the concentration vs. NP conditions and separately for the IS vs. NP conditions. This resulted in two RSMs. We then calculated the Pearson correlation coefficients between these two RSMs. This analysis evaluates whether the neural patterns associated with the concentration vs. NP contrast were similar to those associated with the IS vs. NP contrast.

For the “Scramble” model, we generated RSMs for the IS vs. NP (normal) conditions and separately for the IS vs. NP (scramble) conditions. This resulted in two RSMs for each time window. We then calculated the Pearson correlation coefficients between these two RSMs. This analysis evaluates whether the neural patterns associated with the IS vs. NP (normal) contrast were similar to those associated with the IS vs. NP (scramble) contrast.

For the “Close-eye” model, we generated RSMs for the close-eye vs. NP conditions and separately for the IS vs. NP conditions. This resulted in two RSMs for each time window. We then calculated the Pearson correlation coefficients between these two RSMs. This analysis evaluates whether the neural patterns associated with the close-eye vs. NP contrast were similar to those associated with the IS vs. NP contrast.

### Feature-neural RSA

The analysis was only performed on the 0-6 s time window to shrink the analysis and presentation load. This analysis was conducted in the AIS and VIS experiments. We first created neural RDMs separately for the individual IS and NP conditions and converted to RSMs. Next, we constructed feature RDMs based on four subjective perceptual dimensions: valence, arousal, information load, and processing difficulty. These perceptual dimensions were derived from ratings provided by an independent sample that was demographically matched to the main study participants (further details can be found in the “Experimental materials” section). For each perceptual dimension, we calculated the absolute difference between the ratings of each pair of stimuli (for example, the valence RDM(i,j) = |valence_values(i) − valence_values(j)|). This produced a symmetric feature RDM for each dimension, where larger values represent greater dissimilarity between stimuli on that perceptual feature. The created feature RDMs were then converted to RSMs.

We ensured that the neural RSMs and feature RSMs were properly aligned, so that each element (i, j) in the matrices corresponded to the same pair of stimuli. We then calculated the Spearman correlation coefficient between each feature RSM and the corresponding neural RSM. The Spearman correlation was chosen for its robustness to non-parametric data and its ability to capture monotonic relationships. This correlation quantified the degree to which neural similarity patterns reflected the similarities in subjective perceptual features, providing insights into how well neural representations align with participants’ perceptual experiences.

### Cross-modal ISC analysis

ISC analysis was conducted with two groups of participants exposed to auditory and visual stimuli. For both groups, preprocessed time series of brain activity were extracted for two conditions: IS and NP. The Pearson correlation coefficient was calculated for each pair of participants within the auditory and visual groups, resulting in a cross-modal ISC matrix. Subsequently, ISC values were averaged separately for the IS and NP conditions, yielding cross-modal ISC means across the whole brain. This process involved evaluating the correlation of voxel-level time series between each subject pair in both the auditory and visual groups on a voxel-by-voxel basis. To ensure a standardized reference, only the ISC values corresponding to non-zero voxels in a standard MNI space whole-brain mask were retained. The masked cross-modal ISC results were then saved as a new NIfTI file, preserving the spatial reference information of the input data, which facilitated further statistical analysis and visualization.

### Multiple comparison correction

Brain activation was reported for clusters containing more than 10 voxels if they survived an FDR correction at *p* < 0.05. For all statistical analyses involving multiple comparisons, the *p*-values were further adjusted using an additional FDR correction.

### Neural-behavioral correlation analysis

The correlation analysis was applied to Experiment 1 and 3.

We calculated two-tailed Pearson correlations between IS-related changes in behavioral performance (difference of *P*_sensed_, *P*_understood_, and *ACC*_objective_ between the IS and NP conditions) and fMRI brain activity (difference of activation between the IS and NP conditions on each ROI) across the 0-6, 0-3, and 3–6-time windows.

Statistical significance was set at *p*(FDR) < 0.05.

### Data availability

The data used in this study are not publicly available owing to concerns regarding participant privacy; however, the first or corresponding author will provide deidentified primary data upon request.

### Code availability

The first or corresponding author will provide the code used in this study upon request.

## Results

### Behavioral results

Descriptive statistics for average ratings of each condition are provided in Extended Data 4.

We determined an appropriate sample size *a priori* based on prior experience. To verify whether we had sufficient power to detect a significant behavioral difference between task conditions, *post-hoc* power analyses using the R package SIMR (https://cran.r-project.org/web/packages/simr/index.html) were conducted for all predictors in the best-performing models across the experiments. The analysis revealed that the models from Experiments 1(AIS/VIS), 2.1, 2.2, 2.3, and 4 exhibited robust power, ranging from 87% to 100%. However, the best-performing model for Experiment 3 was underpowered, with predictor power ranging between 9% and 43%. When the model for Experiment 3 was adjusted to repeated measures ANOVAs (within-between interaction) without accounting for random effects, the *post-hoc* power calculation using G*Power software (https://www.psychologie.hhu.de/arbeitsgruppen/allgemeine-psychologie-und-arbeitspsychologie/gpower) indicated that, with a total sample size of 37 participants, there was > 99% power to detect a medium effect size (*f* = 0.5) at *α* = 0.05.

#### Experiment 1

**AIS:** Compared to NP, IS did not produce in different *P*_sensed_ compared to NP [*log-odds* = -0.09, *z* = -0.29*, p* = 0.769*, d* = -0.05, 95% *CI* = (-0.72, 0.53)], but resulted in lower *P*_understood_ [*log-odds* = -1.33, *z* = -7.25*, p* < 0.001*, d* = -0.73, 95% *CI* = (-1.69, -0.97)] and lower *ACC*_objective_ [*log-odds* = -0.44, *z* = -3.53, *p* < 0.001, *d* = -0.24, 95% *CI* = (- 0.68, -0.19)] (Figure 1B).

**VIS:** Compared to NP, IS led to a decrease in *P*_sensed_ [*log-odds* = -0.39, *z* = -2.13*, p* = 0.033*, d* = -0.22, *95% CI* = (-0.75, -0.03)], *P*_understood_ [*log-odds* = -0.57, *z* = -6.03*, p* < 0.001*, d* = -0.32, *95% CI* = (-0.76, -0.38)], and *ACC*_objective_ (*log-odds* = -0.24, *t* = -2.76*, p* = 0.010*, d* = -0.13, *95% CI* = (-0.42, -0.06)] (Figure 1B).

#### Experiment 2

**EXP 2.1 (6-second duration stimuli):** Compared to NP, IS did not significantly affect the *P*_sensed_ [*log-odds* = 0.02, *z* = 0.08, *p* = 0.934, *d* = -0.01, 95% *CI* = (-0.45, 0.49)], but reduced *P*_understood_ [*log-odds* = -0.75, *z* = -4.48, *p* < 0.001, *d* = -0.41, 95% *CI* = (-1.08, - 0.42)], and reduced the *ACC*_objective_ [*log-odds* = -0.24, *z* = -5.04, *p* = 0.002, *d =* -0.13, 95% *CI* = (-0.39, -0.09)] (Extended data1-Figure 1E).

**EXP 2.2 (3-second duration stimuli):** Compared to NP, IS did not significantly affect *P*_sensed_ [*log-odds* = -0.57, *z* = -0.93, *p* = 0.352, *d* = -0.31, 95% *CI* = (-1.77, 0.63)], but resulted in higher *P*_understood_ [*log-odds* = 0.84, *z* = 3.88, *p* < 0.001, *d* = 0.46, 95% *CI* = (0.42, 1.27)], and lower *ACC*_objective_ [*log-odds* = -0.26, *z* = -3.58, *p* < 0.001, *d* = -0.14, 95% *CI* = (-0.40, -0.12)] (Extended data1-Figure 1E)..

**EXP 2.3 (10-second duration stimuli):** Compared to NP, IS entailed lower *P*_sensed_ [*log- odds* = -1.23, *z* = -5.81, *p* < 0.001, *d* = -0.68, 95% *CI* = (-1.65, -0.82)], lower *P*_understood_ [*log-odds* = -1.61, *z* = -17.96, *p* < 0.001, *d* = -0.89, 95% *CI* = (-1.78, -1.43)], as well as lower *ACC*_objective_ [*log-odds* = -0.65, *z* = -9.88, *p* < 0.001, *d* = -0.36, 95% *CI* = (-0.78, - 0.52)] (Extended data1-Figure 1E).

#### Experiment 3

In the lDLPFC-activated group, compared to NP, IS did not significantly affect *P*_sensed_ [*log-odds* = 14.11, *z* = 0.61, *p* = 0.541, *d* = 7.78, 95% *CI* = (-31.08, 59.30)] or *P*_understood_ [*log-odds* = -0.17, *z* = -0.25, *p* = 0.801, *d* = -0.09, 95% *CI* = (-1.47, 1.14)], but resulted in lower *ACC*_objective_ [*log-odds* = -0.66, *z* = -2.00, *p* = 0.045, *d* = -0.36, 95% *CI* = (-1.30, -0.01)]. In the rDLPFC-activated group, IS did not significantly affect *P*_sensed_ [*log-odds* = 14.94, *z* = 0.65, *p* = 0.517, *d* = 8.24, 95% *CI* = (-30.26, 60.15)] or *P*_understood_ [*log-odds* = 0.46, *z* = 0.72, *p* = 0.472, d = 0.25, 95% *CI* = (-0.80, 1.73)], whereas it resulted in marginally lower *ACC*_objective_ in IS compared to NP [*log-odds* = -0.64, *z* = -1.76, *p* = 0.078, *d* = -0.35, 95% *CI* = (-1.35, 0.07)] (Fig. 3C-E). In the sham group, compared to NP, IS did not significantly affect *P*_sensed_ [*log-odds* = -14.93, *z* = -0.65, *p* = 0.517, *d =* - 8.23, 95% *CI* = (-60.08, 30.23)], *P*_understood_ [*log-odds* = -0.42, *z* = -0.81, *p* = 0.422, *d* = - 0.23, 95% *CI* = (-1.45, 0.61)], and *ACC*_objective_ [*log-odds* = -0.23, *z* = -0.97, *p* = 0.331, *d* = -0.12, 95% *CI* = (-0.70, 0.24)].

Regarding group difference in the IS condition, neither of the lDLPFC-activated or rDLPFC-activated group was different from the sham group on the difference in *P*_sensed_ (lDLPFC-activated: *log-odds* = -1.26, *z* = -1.84, *p* = 0.066, *d* = -0.69, *95% CI* = [-2.60, 0.09]; rDLPFC-activated: *log-odds* = -0.46, *z* = -0.61, *p* = 0.541, *d* = -0.25, *95% CI* = [-1.93, 1.01]), *P*_understood_ (lDLPFC-activated: *log-odds* = -0.41, *z* = -0.74, *p* = 0.462, *d* = -0.23, *95% CI* = [-1.49, 0.68]; rDLPFC-activated: *log-odds* = -0.31, *z* = -0.55, *p* = 0.581, *d* = -0.17, *95% CI* = [-1.42, 0.80]). However, lDLPFC-activated group showed lower *ACC*_objective_ than the sham group (*log-odds* = -0.90, *z* = -1.99, *p* = 0.047, *d* = -0.50, *95% CI* = [-1.78, -0.01]), while rDLPFC-activated group was not different from the sham group (*log-odds* = -0.19, *z* = -0.42, *p* = 0.678, *d* = -0.10, *95% CI* = [-1.10, 0.71]).

Regarding group difference in the NP condition, neither of the lDLPFC-activated or rDLPFC-activated group was different from the sham group on the difference in *P*_sensed_ [lDLPFC-activated: *log-odds* = -15.37, *z = -0.67, p* = 0.505*, d* = -8.47, 95% *CI* = (- 60.56, 29.82); rDLPFC-activated: *log-odds* = -15.40, *z* = -0.67, *p* = 0.504, *d* = -8.49, 95% *CI* = (-60.60, 29.79)], *P*_understood_ [lDLPFC-activated: *log-odds* = -0.24, z = -0.29, *p* = 0.769, *d* = -0.13, 95% *CI* = (-1.84, 1.36); rDLPFC-activated: *log-odds* = -0.78, *z* = - 0.97, p = 0.332, *d* = -0.43, 95% *CI* = (-2.35, 0.79)], or *ACC*_objective_ [lDLPFC-activated: *log-odds* = -0.24, *z* = -0.45, *p* = 0.654, *d* = -0.13, 95% *CI* = (-1.29, 0.81); rDLPFC- activated: *log-odds* = 0.45, *z* = 0.80, *p* = 0.426, *d* = 0.25, 95% *CI* = (-0.66, 1.55)].

#### Experiment 4

##### “Concentration” model for AIS

Compared to NP, neither IS [*log-odds* = -0.49, *z* = - 0.64, *p* = 0.520, *d* = -0.27, 95% *CI* = (-1.98, 1.00)] or concentration [*log-odds* = -0.46, *z* = -0.56, *p* = 0.576, *d* = -0.25, 95% *CI* = (-2.07, 1.15)] differed in *P*_sensed_. Concentration did not yield a significant difference in *P*_understood_ compared to NP [*log-odds* = -0.24, *z* = -0.63, *p* = 0.532, *d* = -0.13, 95% *CI* = (-0.99, 0.51)], while IS significantly reduced *P*_understood_ [*log-odds* = -1.74, *z* = -6.93, *p* <0.001, *d* = -0.96, 95% *CI* = (-2.23, -1.25)].

The difference in *ACC*_objective_ between concentration and NP condition was not significant [*log-odds* = 0.21, *z* = 0.88, *p* = 0.378, *d* = 0.12, 95% *CI* = (-0.25, 0.66)], while IS showed significantly lower *ACC*_objective_ compared to NP [*log-odds* = -0.68, *z* = -3.42, *p* = 0.001, *d* = -0.37, 95% *CI* = (-1.07, -0.29)] (Extended data1-Figure 2E).

#### “Scramble” model for AIS

Compared to normal voice, scramble voice resulted in lower *P*_understood_ [*log-odds* = -3.69, *z =* -27.72*, p* < 0.001, *d* = -2.04, 95% CI = (-3.96, 3.43)], demonstrating that the scrambled stimuli conveyed meaningless information.

In normal voices, IS did not entail different *P*_sensed_ [*log-odds* = -0.44, *z =* -0.58, p = 0.561, *d* = -0.24, 95% *CI* = (-1.93, 1.05), but it did result in lower *P*_understood_ [*log-odds* = -1.33, *z =* -4.35, p < 0.001, *d* = -0.73, 95% *CI* = (-1.93, 0.73)] as compared to NP. In scrambled voices, IS did not produce significant differences in either *P*_sensed_ [*log-odds* = -0.63, *z* = -1.00*, p* = 0.317, *d* = -0.35, 95% *CI* = (-1.87, 0.61)] or *P*_understood_ [*log-odds* = -0.19, *z* = -0.20*, p* = 0.843, *d* = -0.10, 95% CI = (-2.12, 1.73)] compared to NP (Fig. 4B). It should be noted that the effect of IS on *P*_understood_ in the scrambled voice condition is not interpretable, as the scrambled stimuli lack semantic content. These findings suggest that when information cannot be received—such as with scrambled voices devoid of meaningful content—active IS is ineffective (Extended data1-Figure 2E).

##### “Close-eye” model for VIS

Compared to NP, close-eye led to a decrease in all of *P*_sensed_ (*log-odds* = -3.79, *z* = 18.08*, p* < 0.001*, d* = -2.09, *95% CI* = [-4.20, -3.38]), *P*_understood_ (*log-odds* = -1.71, *z* = 14.74*, p* < 0.001*, d* = -0.95, *95% CI* = [-1.94, -1.49]), and *ACC*_objective_ (*log-odds* = -1.18, *z* = 10.41*, p* < 0.001*, d* = -0.50, *95% CI* = [-1.40, - 0.96]), demonstrating that the “close-eye” instruction was effective (Extended data1- Figure 2E).

#### EEG results

##### MVPA & Temporal decoding analysis

The analysis was conducted using the EEG datasets of NP condition as the training set, and we examined the decoding success rate for the IS condition EEG datasets over time. A higher decoding rate (indicated by red regions) implies greater dissimilarity between the two conditions.

**3-second duration stimuli:** No significant changes in decoding accuracy were observed. The classification accuracy fluctuates between 0.4 and 0.6, with no distinct patterns indicating sudden changes in brain activity within the 3-second window. The predominance of blue regions suggests that the NP condition and IS condition are generally dissimilar across this timeframe (Extended data1-Figure 1F).

**6-second duration stimuli:** A noticeable shift in decoding accuracy around the 3.4 s mark was observed. The accuracy values range from 0.3 to 0.7, with clear patterns of increased accuracy (red regions) and decreased accuracy (blue regions) around the 3.4 s mark. The more prominent red regions after 3.4 s indicate that the NP condition and IS condition are less similar during this period (Extended data1-Figure 1F).

**10-second duration stimuli:** A distinct change in decoding accuracy around the 3 s mark was observed. The accuracy values range from 0.4 to 0.6, with prominent increases (red regions) and decreases (blue regions) in accuracy around the 3 s mark. The significant red regions after 3 s again suggest greater dissimilarity between the NP condition and IS condition (Extended data1-Figure 1F).

##### Change-point detection

**3-second voice stimuli:** No change point was detected (Extended data1-Figure 1G).

**6-second voice stimuli:** A change point was detected at 3.412 s. The mean decoding rate before the change point (0.56 ± 0.05) was significantly lower than after the change point (0.59 ± 0.04; *t*(498) = -2.22, *p* = 0.027, *d* = 0.98) (Extended data1-Figure 1G).

**10-second voice stimuli:** A change point was detected at 3.003 s. The mean decoding rate before the change point (0.51 ± 0.02) was significantly lower than after the change point (0.54 ± 0.03; *t*(498) = -11.44, *p* < 0.001, *d* = 1.11) (Fig. 2E) (Extended data1- Figure 1G).

#### fMRI results

##### Change-point detection

**AIS:** For both the temporal curves of IS-related brain activation differences and the IS-related decoding rates, a change point was detected at the third TR, corresponding to an average changepoint time of 3–4.5 seconds post-stimulus onset. Furthermore, the average changepoint scan indices observed across all 32 participants fell within the 95% confidence interval for the changepoint time, and the interval itself was very narrow. This indicates that our observed average changepoint demonstrates high stability and reliability (Extended data1-Figure 1H).

**VIS:** Similarly, both the temporal curves of IS-related brain activation differences and the IS-related decoding rates revealed a change point at the third TR, corresponding to an average changepoint time of 3 –4.5 seconds post-stimulus onset. The average changepoint scan indices observed among all 27 participants also fell within the 95% confidence interval for the changepoint time, with a very narrow confidence interval. This consistency indicates that the average changepoint observed is stable and reliable (Extended data1-Figure 1H).

##### Brain activity

The full results of ROI analyses are presented in Extended data 5.

#### Experiment 1

**AIS:** During the 0-6 second period, compared to NP, IS induced greater activation in DLPFC [*post-hoc t*(31) = 9.11, *p*(FDR) < 0.001, *d* = 1.61] and PPC [*post-hoc t*(31) = 4.33, *p*(FDR) < 0.001, *d* = 0.77], and greater deactivation in STC [*post-hoc t*(31) = - 5.35, *p*(FDR) < 0.001, *d* = -0.95] (Extended data1-Figure 1A, and Table 2a).

**Table 2.** *Post*-*hoc* beta values (*M* ± *SD*) for comparisons in Experiment 1: Analysis of voice exposure periods (a) 0-6 seconds, and (b) difference of post-3s vs pre-3s period.

| Items | IS | NP | $t$ (degrees of freedom) | $p$ | $p(\text{FDR})^a$ | Cohen's $d$ |
| --- | --- | --- | --- | --- | --- | --- |
| **AIS** |  |  |  |  |  |  |
| DLPFC | 0.12 ± 0.16 | -0.34 ± 0.27 | 9.11 (31) | < 0.001 | < 0.001*** | 1.61 |
| PPC | 0.26 ± 0.29 | -0.17 ± 0.52 | 4.33 (31) | < 0.001 | < 0.001*** | 0.77 |
| STC | 0.78 ± 0.41 | 1.27 ± 0.62 | -5.35 (31) | < 0.001 | < 0.001*** | -0.95 |
| **VIS** |  |  |  |  |  |  |
| DLPFC | 0.19 ± 0.20 | -0.23 ± 0.20 | 7.99 (26) | <0.001 | <0.001*** | 1.54 |
| PPC | 0.22 ± 0.16 | -0.05 ± 0.42 | 3.28 (26) | 0.003 | 0.005** | 0.63 |
| VC | 0.34 ± 0.11 | 0.49 ± 0.36 | -2.28 (26) | 0.031 | 0.031* | -0.44 |

<sup>a</sup> Corrected using false discovery rate correction method; \*\*\* $p(\text{FDR}) < 0.001$ .

**b**
| Items | 0-3s | 3-6s | *t* (degrees of freedom) | *p* | *p*(FDR)<sup>a</sup> | *Cohen's d* |
| --- | --- | --- | --- | --- | --- | --- |
| **AIS** |  |  |  |  |  |  |
| IS-increased DLPFC | 0.57 ± 0.54 | 0.50 ± 0.38 | -0.79 (31) | 0.437 | 0.437 | -0.14 |
| IS-increased PPC | 0.56 ± 0.96 | 0.28 ± 0.76 | -1.69 (31) | 0.102 | 0.153 | -0.30 |
| IS-decreased STC | -0.71 ± 0.93 | -1.22 ± 0.70 | -3.15 (31) | 0.004 | 0.012* | -0.56 |
| **VIS** |  |  |  |  |  |  |
| IS-increased DLPFC | 0.53 ± 0.43 | 0.36 ± 0.32 | -2.32 (26) | 0.028 | 0.028* | -0.45 |
| IS-increased PPC | 0.46 ± 0.63 | 0.08 ± 0.71 | -3.23 (26) | 0.003 | 0.005** | -0.62 |
| IS-decreased VC | -0.27 ± 0.61 | -0.68 ± 0.51 | -3.91 (26) | 0.001 | 0.003** | -0.75 |

<sup>a</sup> Corrected using false discovery rate correction method; \* $p(\text{FDR}) < 0.05$ .

During the 0-3 and 3-6 second periods, similar patterns (DLPFC and PPC activation, and STC deactivation) were observed. When comparing the IS-related activation or deactivation across the two periods, it was found that the IS-related increase in activity in the DLPFC and PPC were less pronounced in the post-3s period compared to the pre- 3s period, though this difference did not reach significance. However, the IS-related decrease in activity in the STC became more pronounced in the post-3s period and was statistically significant [*post-hoc t*(31) = -3.15, *p*(FDR) = 0.012, *d* = -0.56] (Figure 1C, and Table 2b).

**VIS:** During the 0-6 second period, compared to NP, IS induced greater activation in DLPFC [*post-hoc t*(26) = 7.99, *p*(FDR) < 0.001, *d* = 1.54] and PPC [*post-hoc t*(26) = 3.28, *p*(FDR) = 0.005, *d* = 0.63], and greater deactivation in VC [*post-hoc t*(26) = -2.28, *p*(FDR) = 0.031, *d* = -0.44] (Extended data1-Figure 1B, and Table 2a).

During the 0-3 and 3-6 second periods, similar patterns (DLPFC and PPC activation, and VC deactivation) were observed. Notably significant activation of the DLPFC and PPC was not observed, but prominent deactivation in these regions was evident. When comparing the IS-related activation or deactivation across the two periods, it was found that the IS-related increase in activity in the DLPFC and PPC were less pronounced in the post-3s period compared to the pre-3s period [DLPFC: *post-hoc t*(26) = -2.32, *p*(FDR) = 0.028, *d* = -0.45; PPC: *post-hoc t*(26) = -3.23, *p*(FDR) = 0.005, *d* = -0.62]. In contrast, the IS-related decrease in activity in the VC became more pronounced in the post-3s period compared to the pre-3s period [*post-hoc t*(26) = -3.91, *p*(FDR) = 0.003, *d* = -0.75] (Figure 1C, and Table 2b).

#### Experiment 3

During the 0-6 second period, compared to the sham group, lDLPFC- activated group had more prominent IS-related activation in DLPFC, largely in the left potion, while rDLPFC-activated group had more prominent IS-related activation in DLPFC, largely in the right potion, proved the effectiveness of the TMS intervention (Extended data1-Figure 2B). According to the *post-hoc* ANOVA of mean parameter estimates for each ROI, the IS-related increase in DLPFC activation differed among the three TMS groups [*F*(2, 34) = 4.70, *p* = 0.048, *η²* = 0.22]. Specifically, the lDLPFC group exhibiting a more prominent IS-related increase DLPFC activation compared to the sham group (*p* = 0.033), and the rDLPFC group also showing a more prominent IS-related increase DLPFC activation compared to the sham group (*p* = 0.006; Table 3a).

**Table 3.** Post-hoc beta values (M ± SD) under IS-NP differences across all TMS groups for comparisons in Experiment 3: Analysis of voice exposure periods (a) 0-6 seconds, and (b) difference of post-3s vs pre-3s period.

| Items | lDLPFC-activation group | rDLPFC-activation group | Sham group | *F* (degrees<br>of freedom) | *p* | *p*(FDR)<sup>a</sup> | $\eta^2$ |
| --- | --- | --- | --- | --- | --- | --- | --- |
| IS-increased DLPFC | 0.47 ± 0.39 | 0.58 ± 0.34 | 0.16 ± 0.31 | 4.70(2,34) | 0.016 | 0.048* | 0.22 |
| IS-increased PPC | 0.16 ± 0.55 | 0.57 ± 0.86 | 0.09 ± 0.82 | 1.46(2,34) | 0.246 | 0.369 | 0.08 |
| IS-decreased STC | -0.31 ± 0.58 | -0.45 ± 0.53 | -0.62 ± 0.52 | -0.99(2,34) | 0.382 | 0.382 | 0.06 |

<sup>a</sup>Corrected using false discovery rate correction method; \* $p < 0.05$ .

b
| Items | lDLPFC-activation<br>group | rDLPFC<br>activation<br>group | sham group | *F* (degrees<br>of<br>freedom) | *p* | *p*(FDR)<sup>a</sup> | $\eta^2$ |
| --- | --- | --- | --- | --- | --- | --- | --- |
| DLPFC activation (post3s-pre3s) | -0.58 ± 0.27 | -0.52 ± 0.44 | -0.06 ± 0.61 | 4.73(2,34) | 0.015 | 0.045* | 0.22 |
| PPC activation (post3s-pre3s) | -0.46 ± 0.77 | -0.18 ± 1.34 | -0.01 ± 0.93 | 0.60(2,34) | 0.557 | 0.557 | 0.03 |
| STC activation (post3s-pre3s) | -0.98 ± 0.81 | -1.37 ± 0.92 | -0.94 ± 0.64 | 1.07(2,34) | 0.355 | 0.533 | 0.06 |

<sup>a</sup>Corrected using false discovery rate correction method; \* $p(\text{FDR}) < 0.05$ .

During the 0-3 and 3-6 second periods, the flexible factorial analysis demonstrated significant interaction effects between *TMS group* × *time* in the DLPFC and PPC. According to the *post-hoc* ANOVA of mean parameter estimates for each ROI, these interactions were explained by the fact that, the IS-related increase in DLPFC activation during the post-3s period compared to the pre-3s period [i.e., post 3s (IS-NP) vs. pre 3s (IS-NP)] differed among the three TMS groups [*F*(2, 34) = 4.73, *p* = 0.045, *η²* = 0.22]. Specifically, the lDLPFC-activated group exhibited a more prominent post-pre difference in IS-related increase in DLPFC activation compared to the sham group (*p* = 0.008), and the rDLPFC-activated group also showed a more prominent post-pre difference in IS-related increase in DLPFC activation compared to the sham group (*p* = 0.018; Table 3b).

#### Experiment 4

**“Concentration” model:** During the 0-6 second periods, IS vs. NP contrast observed similar brain activity patterns (higher DLPFC and PPC activation, and STC deactivation). The concentration vs. NP contrast only observed higher DLPFC and PPC activation. When comparing the two contrasts, it was found that IS vs. NP induced higher DLPFC and PPC activation [*post-hoc* DLPFC: *t*(14) = -3.96, *p*(FDR) = 0.003, *d* = -1.02; PPC: *t*(14) = -3.26, *p*(FDR) = 0.008, *d* = -0.84], and STC deactivation [*post-hoc t*(31) = 3.11, *p*(FDR) = 0.008, *d* = 0.80] (Extended data1-Figure2F and Table 4a).

**Table 4.** *Post-hoc* beta values (*M* ± *SD*) for concentration vs. NP and IS vs. NP conditions for comparisons in the “Concentration” model of Experiment 4: Analysis of voice exposure periods (a) 0-6 seconds, and (b) difference of post-3s vs. pre-3s period.

| Items | Concentration vs. NP | IS vs. NP | $t$ (degrees | $p$ | $p(\text{FDR})^a$ | Cohen's $d$ |
| --- | --- | --- | --- | --- | --- | --- |

|  |  |  | of freedom) |  |  |  |
| --- | --- | --- | --- | --- | --- | --- |
| DLPFC | $0.18 \pm 0.41$ | $0.48 \pm 0.34$ | -3.96 (14) | 0.001 | 0.003** | -1.02 |
| PPC | $-0.18 \pm 0.84$ | $0.31 \pm 0.64$ | -3.26 (14) | 0.006 | 0.008** | -0.84 |
| STC | $-0.08 \pm 1.01$ | $-0.60 \pm 0.56$ | 3.11 (14) | 0.008 | 0.008** | 0.80 |

<sup>a</sup>Corrected using false discovery rate correction method; \*\* $p < 0.01$ .

b
| Items | Post-3s(concentration vs. NP) -<br>Pre-3s(concentration vs. NP) | Post-3s(IS vs. NP) -<br>Pre-3s(IS vs. NP) | $t$ (degrees of<br>freedom) | $p$ | $p(\text{FDR})^a$ | *Cohen's d* |
| --- | --- | --- | --- | --- | --- | --- |
| DLPFC | $0.33 \pm 0.51$ | $-0.24 \pm 0.49$ | 2.38(14) | 0.032 | 0.048* | 0.62 |
| PPC | $0.14 \pm 1.16$ | $-0.09 \pm 0.85$ | 0.46(14) | 0.652 | 0.652 | 0.12 |
| STC | $0.96 \pm 1.17$ | $-1.24 \pm 0.81$ | 4.60(14) | 0.001 | 0.003** | 1.19 |

<sup>a</sup>Corrected using false discovery rate correction method; \* $p(\text{FDR}) < 0.05$ , \*\* $p(\text{FDR}) < 0.01$ .

During the 0-3 and 3-6 second periods, the flexible factorial analysis demonstrated significant interaction effects between *task condition* × *time* in the DLPFC and STC. According to the *post-hoc* ANOVA of mean parameter estimates for each ROI, these interactions were explained by the fact that, the post3s – pre3s difference in DLPFC and STC activation in IS was prominent compared to the concentration condition [*post- hoc* DLPFC: *t*(14) = 2.38, *p*(FDR) = 0.048, *d* = 0.62; STC: *t*(14) = 4.60, *p*(FDR) = 0.003, *d* = 1.19] (Table 4b).

**“Scramble” model:** During the 0-6 second periods, both IS vs. NP normal, and IS vs. NP scramble contrasts observed similar brain activity patterns (higher DLPFC and PPC activation, and STC deactivation). When comparing the two contrasts, it was found that IS vs. NP normal induced higher DLPFC activation [*post-hoc t*(14) = -2.85, *p*(FDR) = 0.039, *d* = -0.74] (Extended data1-Figure 2F and Table 5a).

**Table 5.**
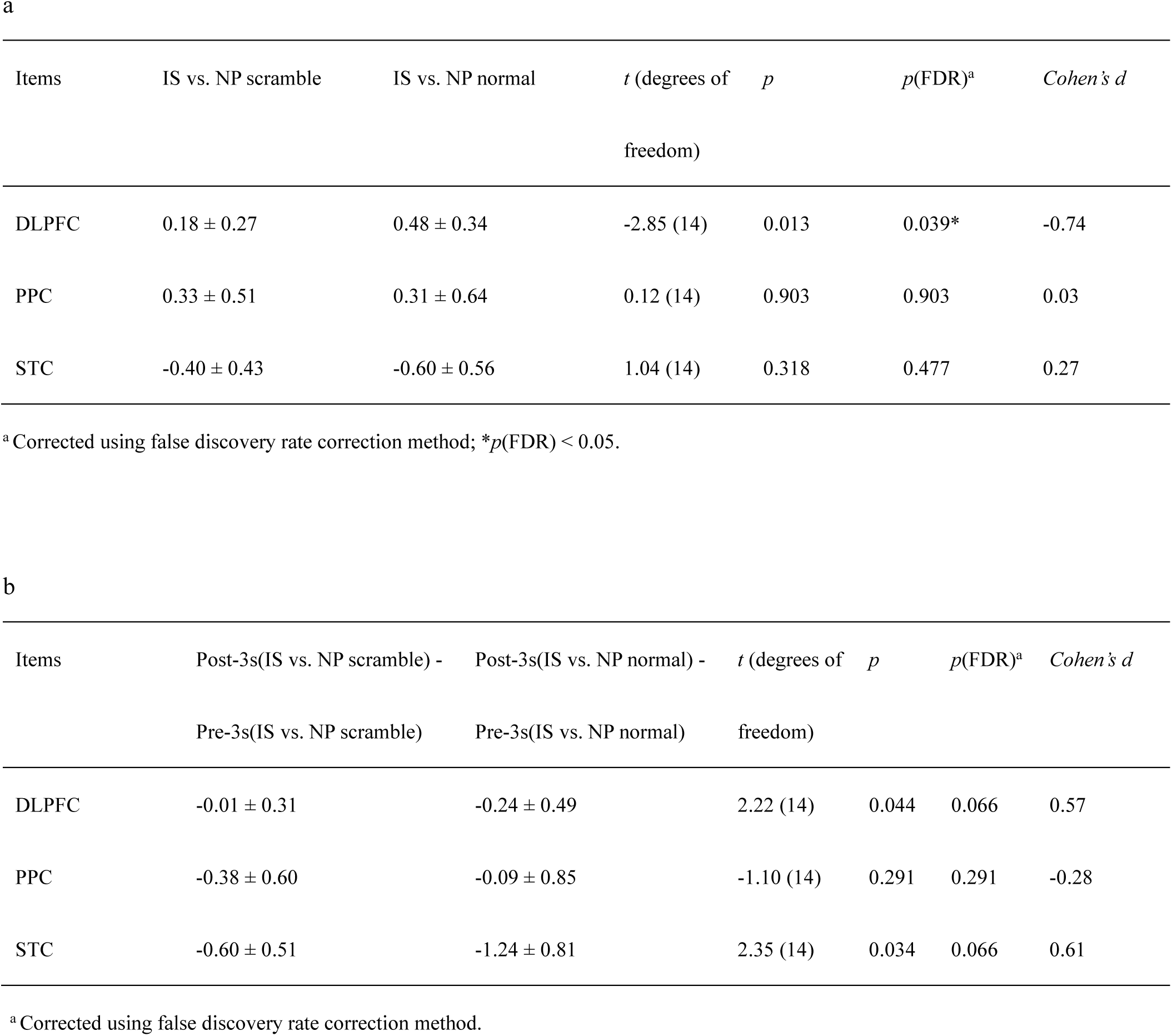
*Post-hoc* beta values (*M* ± *SD*) for IS vs. NP scramble and IS vs. NP normal conditions for comparisons in the “Scramble” model of Experiment 4: Analysis of voice exposure periods (a) 0-6 seconds, and (b) difference of post-3s vs. pre-3s period.

During the 0-3 and 3-6 second periods, the flexible factorial analysis did not reveal significant interaction effects between *task condition* × *time* × *voice type* in all ROIs (Table 5b).

##### Unimodal MVPA

Descriptive statistics and statistical tests against chance level (0.5) using one-sample *t*- tests for the decoding accuracy of each condition are provided in Extended Data 6 - Table 8.

#### Experiment 2

**AIS:** During the 0-6 second period, both NP and IS conditions were decodable in each of the three ROIs [all *t*(31) > 2.55, *p*(FDR) < 0.019, *d* > 0.64], with the exception of NP in the DLPFC, which was not decodable. Notably, IS was better decoded than NP in the DLPFC [*t*(31) = 2.30, *p*(FDR) = 0.042, *d* = 0.41] while NP was better decoded than IS in the STC [*t*(31) = -2.66, *p*(FDR) = 0.036, *d* = -0.47]. This suggests that compared to NP, IS increased decoding accuracy in the DLPFC and PPC but decreased it in the STC (Table 6a).

**Table 6.**
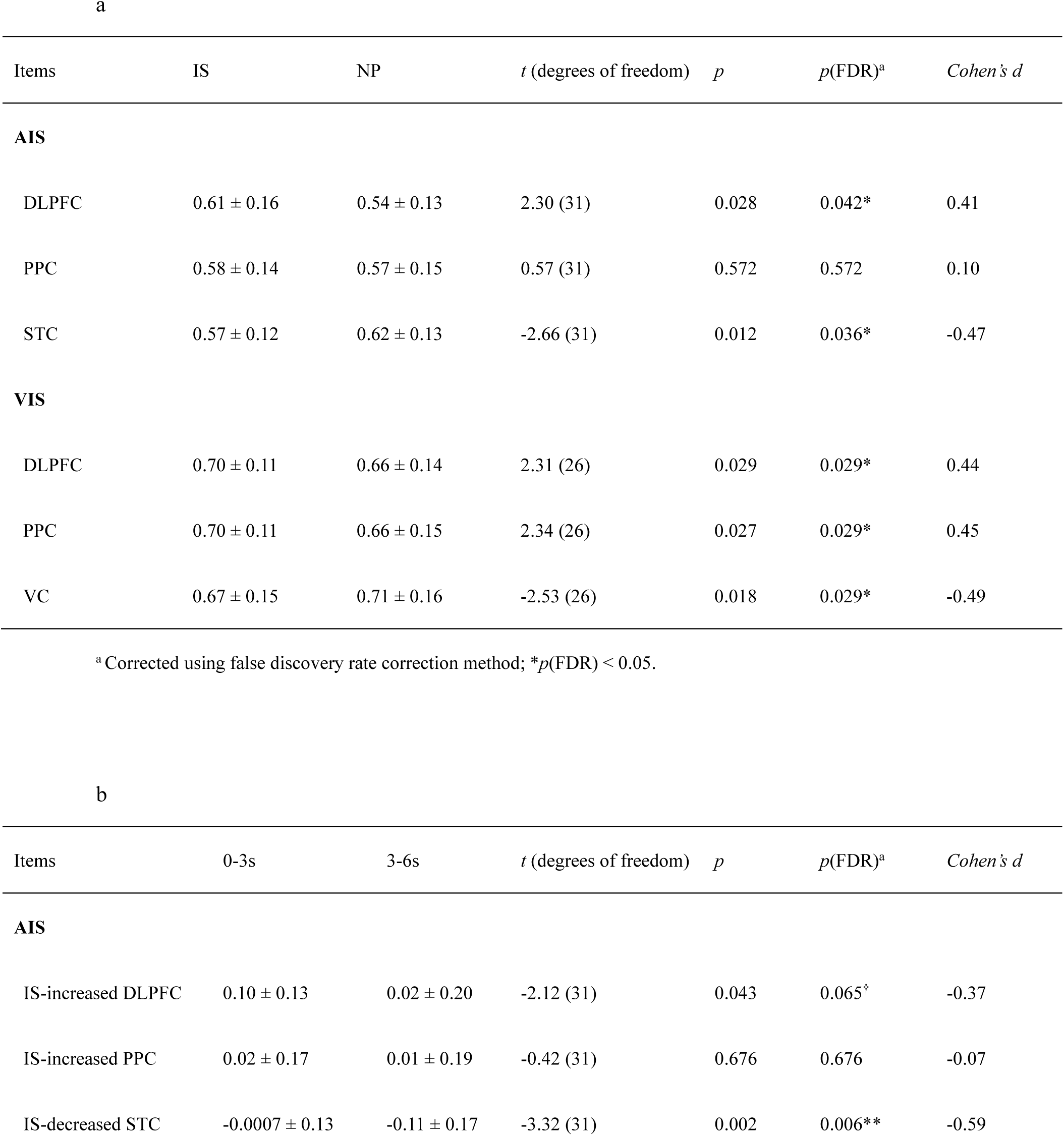
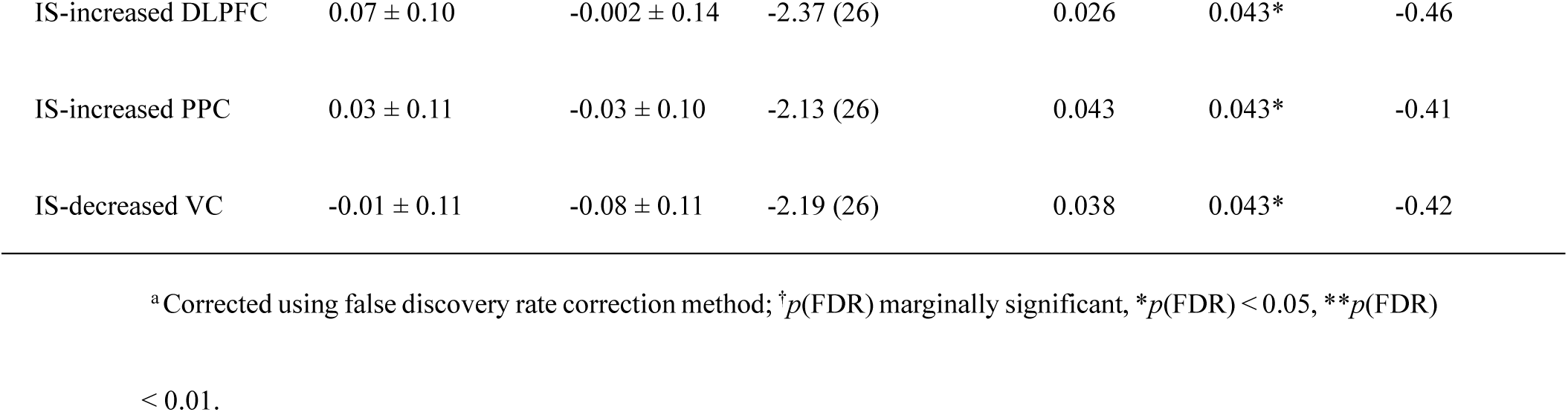
Results of unimodal MVPA decoding accuracies (*M* ± *SD*) in Experiment 1: Analysis of voice exposure periods (a) 0-6 seconds, and (b) difference of post-3s vs. pre-3s period.

| Items | IS | NP | $t$ (degrees of freedom) | $p$ | $p(\text{FDR})^a$ | Cohen's $d$ |
| --- | --- | --- | --- | --- | --- | --- |
| **AIS** |  |  |  |  |  |  |
| DLPFC | $0.61 \pm 0.16$ | $0.54 \pm 0.13$ | 2.30 (31) | 0.028 | 0.042* | 0.41 |
| PPC | $0.58 \pm 0.14$ | $0.57 \pm 0.15$ | 0.57 (31) | 0.572 | 0.572 | 0.10 |
| STC | $0.57 \pm 0.12$ | $0.62 \pm 0.13$ | -2.66 (31) | 0.012 | 0.036* | -0.47 |
| **VIS** |  |  |  |  |  |  |
| DLPFC | $0.70 \pm 0.11$ | $0.66 \pm 0.14$ | 2.31 (26) | 0.029 | 0.029* | 0.44 |
| PPC | $0.70 \pm 0.11$ | $0.66 \pm 0.15$ | 2.34 (26) | 0.027 | 0.029* | 0.45 |
| VC | $0.67 \pm 0.15$ | $0.71 \pm 0.16$ | -2.53 (26) | 0.018 | 0.029* | -0.49 |

<sup>a</sup>Corrected using false discovery rate correction method; \* $p(\text{FDR}) < 0.05$ .

**b**
| Items | 0-3s | 3-6s | $t$ (degrees of freedom) | $p$ | $p(\text{FDR})^a$ | Cohen's $d$ |
| --- | --- | --- | --- | --- | --- | --- |
| **AIS** |  |  |  |  |  |  |
| IS-increased DLPFC | $0.10 \pm 0.13$ | $0.02 \pm 0.20$ | -2.12 (31) | 0.043 | 0.065<sup>†</sup> | -0.37 |
| IS-increased PPC | $0.02 \pm 0.17$ | $0.01 \pm 0.19$ | -0.42 (31) | 0.676 | 0.676 | -0.07 |
| IS-decreased STC | $-0.0007 \pm 0.13$ | $-0.11 \pm 0.17$ | -3.32 (31) | 0.002 | 0.006** | -0.59 |
| VIS |  |  |  |  |  |  |
| IS-increased DLPFC | 0.07 ± 0.10 | -0.002 ± 0.14 | -2.37 (26) | 0.026 | 0.043* | -0.46 |
| IS-increased PPC | 0.03 ± 0.11 | -0.03 ± 0.10 | -2.13 (26) | 0.043 | 0.043* | -0.41 |
| IS-decreased VC | -0.01 ± 0.11 | -0.08 ± 0.11 | -2.19 (26) | 0.038 | 0.043* | -0.42 |

<sup>a</sup>Corrected using false discovery rate correction method; <sup>†</sup> $p(\text{FDR})$ marginally significant, $*p(\text{FDR}) < 0.05$ , $**p(\text{FDR})$

During the 0-3 second period, similar patterns were observed, with both NP and IS conditions being decodable in all three ROIs [all *t*(31) > 3.31, *p*(FDR) < 0.005, *d* > 0.83], except for NP in the DLPFC. The same trend persisted in the 3-6 second period [all *t*(31) > 2.19, *p*(FDR) < 0.048, *d* > 0.55], except for NP in the DLPFC and IS in the STC. When comparing the decoding accuracy differences between IS and NP across time periods, it was found that the IS-related increase in decoding accuracy in the DLPFC and PPC was less pronounced in the post-3s period compared to the pre-3s period, though this difference did not reach significance. However, the IS-related decrease in decoding accuracy in the STC became more pronounced in the post-3s period compared to the pre-3s period, and was statistically significant [*t*(31) = -3.32, *p*(FDR) = 0.006, *d* = -0.59] (Table 6b).

**VIS:** During the 0-6 second period, both NP and IS conditions were decodable in each of the three ROIs, with the exception of NP in the DLPFC [all *t*(26) > 5.74, *p*(FDR) < 0.001, *d* > 1.07]. Notably, IS was better decoded than NP in the DLPFC [*t*(26) = 2.31, *p*(FDR) = 0.029, *d* = 0.44] and PPC[*t*(26) = 2.34, *p*(FDR) = 0.029, *d* = 0.45] while NH was better decoded than IS in the VC [*t*(26) = -2.53, *p*(FDR) = 0.029, *d* = -0.49]. This suggests that compared to NP, IS increased decoding accuracy in the DLPFC and PPC but decreased it in the VC (Table 6a).

During the 0-3 second period, similar patterns were observed, with both NP and IS conditions being decodable in all three ROIs [all *t*(26) > 3.76, *p*(FDR) < 0.001, *d* > 0.72]. The same trend persisted in the 3-6 second period [all *t*(26) > 4.25, *p*(FDR) < 0.001, *d* > 0.81]. When comparing the decoding accuracy differences between IS and NP across time periods, it was found that the IS-related increase in decoding accuracy in the all three ROIs was less pronounced in the post-3s period compared to the pre-3s period [DLPFC: *t*(26) = -2.37, *p*(FDR) = 0.043, *d* = -0.46; PPC: *t*(26) = -2.13, *p*(FDR) = 0.043, *d* = -0.41]. In contrast, the IS-related decrease in decoding accuracy in the VC became more pronounced in the post-3s period compared to the pre-3s period [*t*(26) = -2.19, *p*(FDR) = 0.043, *d* = -0.42] (Table 6b).

#### Experiment 3

During the 0-6 second period, the IS-related increase in decoding accuracy in the DLPFC differed among the three TMS groups [*F*(2, 34) = 5.15, *p* = 0.011, *η²* = 0.23]. This effect was driven by the lDLPFC group showing a higher IS- related increase in decoding accuracy in the DLPFC compared to the sham group (*p* = 0.048), as well as the rDLPFC group displaying a higher IS-related increase in decoding accuracy in the DLPFC compared to the sham group (Table 7a).

**Table 7.** Results of unimodal MVPA decoding accuracies (*M* ± *SD*) under IS-NP differences across all TMS groups for comparisons in Experiment 3: Analysis of voice exposure periods (a) 0-6 seconds, and (b) difference of post-3s vs pre-3s period.

| Items | lDLPFC-activation group | rDLPFC-activation group | sham group | $F$ (degrees<br>of freedom) | $p$ | $p(\text{FDR})^a$ | $\eta^2$ |
| --- | --- | --- | --- | --- | --- | --- | --- |
| IS-increased DLPFC | $0.08 \pm 0.08$ | $0.11 \pm 0.08$ | $0.01 \pm 0.08$ | 5.15(2,34) | 0.011 | 0.033* | 0.23 |
| IS-increased PPC | $-0.01 \pm 0.07$ | $0.02 \pm 0.07$ | $-0.02 \pm 0.18$ | 0.35(2,34) | 0.705 | 0.705 | 0.02 |
| IS-decreased STC | $-0.06 \pm 0.09$ | $-0.01 \pm 0.10$ | $-0.01 \pm 0.13$ | 0.84(2,34) | 0.440 | 0.660 | 0.05 |

<sup>a</sup> Corrected using false discovery rate correction method; \* $p(\text{FDR}) < 0.05$ .

**b**
| Items | lDLPFC-activation<br>group | rDLPFC-activation<br>group | sham group | $F$ (degrees<br>of freedom) | $p$ | $p(\text{FDR})^a$ | $\eta^2$ |
| --- | --- | --- | --- | --- | --- | --- | --- |
| DLPFC activation (post3s - pre3s) | $-0.19 \pm 0.11$ | $-0.15 \pm 0.08$ | $-0.04 \pm 0.17$ | 4.70(2,34) | 0.016 | 0.048* | 0.22 |
| PPC activation (post3s - pre3s) | $-0.06 \pm 0.12$ | $-0.08 \pm 0.13$ | $-0.04 \pm 0.14$ | 0.32(2,34) | 0.727 | 0.727 | 0.02 |
| STC activation (post3s - pre3s) | $-0.02 \pm 0.13$ | $-0.11 \pm 0.12$ | $-0.07 \pm 0.16$ | 1.38(2,34) | 0.265 | 0.398 | 0.07 |

<sup>a</sup> Corrected using false discovery rate correction method; \* $p(\text{FDR}) < 0.05$ .

**Table 8.** Descriptive statistics of cross-modal MVPA decoding accuracies (*M* ± *SD*) in Experiment 1: Analysis of stimuli exposure periods (a) 0-6 seconds, and (b) difference of post-3s vs. pre-3s period.

| Items | IS | NP | $t$ (degrees of freedom) | $p$ | $p(\text{FDR})^a$ | Cohen's $d$ |
| --- | --- | --- | --- | --- | --- | --- |
| **Auditory modality as training set, visual modality as testing set** |  |  |  |  |  |  |
| CEN | $0.65 \pm 0.18$ | $0.45 \pm 0.16$ | 3.11 (26) | 0.004 | 0.008** | 0.60 |
| **Visual modality as training set, auditory modality as testing set** |  |  |  |  |  |  |
| CEN | $0.56 \pm 0.16$ | $0.49 \pm 0.14$ | 1.32 (31) | 0.197 | 0.197 | 0.23 |

| Items | 0-3s | 3-6s | $t$ (degrees of freedom) | $p$ | $p(\text{FDR})^a$ | Cohen's $d$ |
| --- | --- | --- | --- | --- | --- | --- |
| **Auditory modality as training set, visual modality as testing set** |  |  |  |  |  |  |
| CEN (IS>NP) | $0.13 \pm 0.23$ | $0.25 \pm 0.23$ | 2.61 (26) | 0.015 | 0.030* | 0.50 |
| **Visual modality as training set, auditory modality as testing set** |  |  |  |  |  |  |
| CEN (IS>NP) | $0.01 \pm 0.21$ | $0.10 \pm 0.21$ | 1.97 (31) | 0.057 | 0.057<sup>†</sup> | 0.35 |

<sup>a</sup>Corrected using false discovery rate correction method; <sup>†</sup> $p(\text{FDR})$ marginally significant, \* $p(\text{FDR}) < 0.05$ .

Similarly, the IS-related increase in decoding accuracy in the DLPFC during the post- 3s period compared to the pre-3s period [i.e., post 3s (IS-NP) vs. pre 3s (IS-NP)] differed among the three TMS groups [*F*(2, 34) = 4.70, *p* = 0.016, *η²* = 0.22]. This was driven by the lDLPFC group exhibiting a more prominent post-pre difference in IS- related increase in decoding accuracy in the DLPFC compared to the sham group (*p* = 0.005), and the rDLPFC group also showing a more prominent post-pre difference in IS-related increase in decoding accuracy in the DLPFC compared to the sham group (*p* = 0.040; Table 7b).

##### Feature MVPA

Descriptive statistics and statistical tests against chance level (0.25) using one-sample *t*-tests for the decoding accuracy of each condition are provided in Extended data 6- Table10.

**Valence:** The accuracy of all conditions was greater than the chance level of 0.25, which indicates that all conditions are decodable [all *t*(26) > 4.70, *p*(FDR) < 0.001, *d* > 0.90]. There was a significant interaction (*task condition*\**feature intensity*) on decoding accuracy [*F*(1, 26) = 6.06, *p* = 0.021, *η^2^* = 0.003]. A simple effects analysis indicated that, high and low arousal can be distinguished in NP [*F*(1, 26) = 6.06, *p* = 0.021, *η²* = 0.28], but not in IS [*F*(1,26) = 0.08, *p* = 0.786, *η²* = 0.003].

**Arousal:** The accuracy of all conditions was greater than the chance level of 0.25, which indicates that all conditions are decodable [all *t*(26) > 4.48, *p*(FDR) < 0.001, *d* > 0.86]. The interaction (task condition*feature intensity) on decoding accuracy was not significant [*F*(1, 26) = 3.07, *p* = 0.091, *η²* = 0.11].

**Information load:** The accuracy of all conditions was greater than the chance level of 0.25, which indicates that all conditions are decodable [all *t*(26) > 4.35, *p*(FDR) < 0.001, *d* > 0.84]. There was a significant interaction (*task condition*\**feature intensity*) on decoding accuracy [*F*(1, 26) = 9.73, *p* = 0.004, *η²* = 0.27]. A simple effects analysis indicated that, high and low arousal can be distinguished in NP [*F*(1, 26) = 7.74, *p* = 0.010, *η²* = 0.23], but not in IS [*F*(1, 26) = 3.50, *p* = 0.073, *η²* = 0.12].

**Processing difficulty**: The accuracy of all conditions was greater than the chance level of 0.25, which indicates that all conditions are decodable [all *t*(26) > 3.49, *p*(FDR) < 0.002, *d* > 0.67]. There was a significant interaction (task condition*feature intensity) on decoding accuracy [*F*(1, 26) = 9.55, *p* = 0.005, *η²* = 0.27]. A simple effects analysis indicated that, high and low arousal can be distinguished in NP [*F*(1,26) = 27.26, *p* < 0.001, *η²* = 0.51], but not in IS [F(1, 26) = 0.94, *p* = 0.341, *η²* = 0.06].

##### Cross-modal MVPA

Descriptive statistics and statistical tests against chance level (0.5) using one-sample *t*- tests for the decoding accuracy of each condition are provided in Extended Data 5 - Table 9.

**Table 9.** Correlations between participants’ IS-related behavioral efficacy and IS- related brain activation or deactivation (from Experiment 1).

| Items | $r$ | $p$ | $p(\text{FDR})^a$ |
| --- | --- | --- | --- |
| **AIS** |  |  |  |
| IS-decreased $P_{\text{sensed}}$ & IS-increased DLPFC activity | 0.118 | 0.522 | 0.671 |
| IS-decreased $P_{\text{sensed}}$ & IS-increased PPC activity | -0.063 | 0.732 | 0.824 |
| IS-decreased $P_{\text{sensed}}$ & IS-decreased STC activity | -0.190 | 0.297 | 0.446 |
| IS-decreased $P_{\text{understood}}$ & IS-increased DLPFC activity | -0.464 | 0.007 | 0.032* |
| IS-decreased $P_{\text{understood}}$ & IS-increased PPC activity | -0.238 | 0.189 | 0.382 |
| IS-decreased $P_{\text{understood}}$ & IS-decreased STC activity | 0.440 | 0.012 | 0.036* |
| IS-decreased $ACC_{\text{objective}}$ & IS-increased DLPFC activity | -0.227 | 0.212 | 0.382 |
| IS-decreased $ACC_{\text{objective}}$ & IS-increased PPC activity | -0.024 | 0.897 | 0.897 |
| IS-decreased $ACC_{\text{objective}}$ & IS-decreased STC activity | 0.510 | 0.003 | 0.027* |
| **VIS** |  |  |  |
| IS-decreased $P_{\text{sensed}}$ & IS-increased DLPFC activity | -0.434 | 0.024 | 0.054 |
| IS-decreased $P_{\text{sensed}}$ & IS-increased PPC activity | -0.417 | 0.030 | 0.054 |
| IS-decreased $P_{\text{sensed}}$ & IS-decreased VC activity | -0.154 | 0.443 | 0.498 |
| IS-decreased $P_{\text{understood}}$ & IS-increased DLPFC activity | -0.369 | 0.058 | 0.085 |
| IS-decreased $P_{\text{understood}}$ & IS-increased PPC activity | -0.359 | 0.066 | 0.085 |
| IS-decreased $P_{\text{understood}}$ & IS-decreased VC activity | 0.458 | 0.016 | 0.048* |
| IS-decreased $ACC_{\text{objective}}$ & IS-increased DLPFC activity | -0.565 | 0.002 | 0.009** |
| IS-decreased $ACC_{\text{objective}}$ & IS-increased PPC activity | -0.617 | 0.001 | 0.009** |
| IS-decreased $ACC_{\text{objective}}$ & IS-decreased VC activity | 0.090 | 0.655 | 0.655 |

<sup>a</sup> Corrected using false discovery rate correction method; \* $p(\text{FDR}) < 0.05$ , \*\* $p(\text{FDR}) < 0.01$ .

**Auditory modality as training set, visual modality as testing set:** During the 0-6 second period, the IS condition was modality-generalizable [*t*(26) = 4.25, *p*(FDR) < 0.001, *d* = 0.82], whereas the NP condition was not. Additionally, IS exhibited better modality-generalization compared to NP [*t*(26) = 3.11, *p*(FDR) = 0.008, *d* = 0.60] (Table 8a).

During the 0-3 second period, similar patterns were observed, the IS condition was modality-generalizable [*t*(26) = 4.36, *p*(FDR) < 0.001, *d* = 0.84], whereas the NP condition was not. During the 3-6 second period, the IS condition remained modality-generalizable [*t*(26) = 6.51, *p*(FDR) < 0.001, *d* = 1.25], whereas the NP condition continued to lack modality-generalizability. When comparing the differences in generalization accuracy between IS and NP across time periods, it was found that the IS-related increase in generalization accuracy was more pronounced in the post-3s period compared to the pre-3s period [*t*(26) = 2.61, *p*(FDR) = 0.030, *d* = 0.50]. This suggest that, compare to the early 3-second period, IS demonstrated greater modality- generalizability than NP during the late 3-second period (Table 8b).

**Visual modality as training set, auditory modality as testing set:** During the 0-6 second period, the IS condition demonstrated cross-modal generalizability [*t*(31) = 2.12, *p* = 0.042], though this effect did not survive FDR correction [*p*(FDR) = 0.085, *d* = 0.085], whereas the NP condition did not show cross-modal generalizability. Additionally, there was no significant difference in generalization accuracy between the two conditions (Table 8a).

During the 0-3 second period, the IS condition was modality-generalizable [*t*(31) = 3.57, *p*(FDR) = 0.005, *d* = 0.63], whereas the NP condition was not. However, during the 3-6 second period, neither the NP nor IS condition exhibited modality- generalizability. When comparing the differences in generalization accuracy between IS and NP across time periods, it was observed that the IS-related increase in generalization accuracy was more pronounced in the post-3s period compared to the pre-3s period, although this effect only reached marginal significance [*t*(31) = 1.97, *p*(FDR) = 0.057, *d* = 0.35] (Table 8b).

##### Neural-neural RSA

**“Concentration” model for AIS:** During the 0-6 second period, the similarity between the RSMs for concentration vs. NP and IS vs. NP was relatively low (*r* = 0.06, *p*(FDR) = 0.496). During the 0-3 second period, the similarity between the RSMs for concentration vs. NP and IS vs. NP remained low (*r* = 0.03, *p*(FDR) = 0.627). During the 3-6 second period, the similarity between the RSMs for concentration vs. NP and IS vs. NP was low (*r* = 0.09, *p*(FDR) = 0.350; Figure 4F).

**“Scramble” model for AIS:** During the 0-6 second period, the similarity between the RSMs for IS vs. NP normal and IS vs. NP scramble was relatively high (*r* = 0.12, *p*(FDR) < 0.044). During the 0-3 second period, the similarity between the RSMs for IS vs. NP normal and IS vs. NP scramble remained high (*r* = 0.21, *p*(FDR) < 0.001). During the 3-6 second period, the similarity between the RSMs for IS vs. NP normal and IS vs. NP scramble decreased, though it was still significant (*r* = 0.35, *p*(FDR) < 0.001, Figure 4F).

**“Close-eye” model for VIS:** During the 0-6 second period, the similarity between the RSMs for IS vs. NP and close-eye vs. NP scramble was relatively high (*r* = 0.92, *p*(FDR) < 0.001). During the 0-3 second period, the similarity between the RSMs for IS vs. NP normal and IS vs. NP scramble remained high (*r* = 0.87, *p*(FDR) < 0.001). During the 3-6 second period, the similarity between the RSMs for IS vs. NP normal and IS vs. NP scramble decreased, though it was still significant (*r* = 0.90, *p*(FDR) < 0.001, Extended data1-Figure 2G).

##### Feature-neural RSA

In the AIS experiment, the feature RSMs for valence and arousal were significantly correlated with the neural RSM of the NP condition [valence: *r* = 0.14, *p*(FDR) = 0.007; arousal: *r* = 0.18, *p*(FDR) < 0.001] but not with that of the IS condition [valence: *r* = 0.09, *p*(FDR) = 0.243; arousal: *r* = -0.26, *p*(FDR) = 1]. Fisher *r*-*z* transformation^58^ revealed that the correlation (arousal RSM and neural RSM) was weaker in IS than in NP conditions [*z* = -2.55, *p*(FDR) = 0.010]. The feature RSMs for information load and processing difficulty did not show significant correlations with the neural RSMs of either the NP or IS conditions (Figure 2B).

In the VIS experiment, the feature RSMs for valence, arousal, and processing difficulty were all significantly correlated with the neural RSM of the NP condition [valence: *r* = 0.08, *p*(FDR) = 0.028; arousal: *r* = 0.23, *p*(FDR) < 0.001; processing difficulty: *r* = 0.11, *p*(FDR) < 0.001] but not with that of the IS condition [valence: *r* = -0.43, *p*(FDR) = 1; arousal: *r* = -0.04, *p*(FDR) = 1; processing difficulty: *r* = -0.11, *p*(FDR) = 1]. Fisher *r*-*z* transformation revealed that the correlation (valence RSM and neural RSM) was weaker in IS than in NP conditions [*z* = -2.99, *p*(FDR) = 0.003]. The correlation (arousal RSM and neural RSM) was marginally weaker in IS than in NP conditions [*z* = -1.57, *p*(FDR) = 0.0582]. The feature RSM for information load did not exhibit significant correlations with the neural RSMs of either the NP or IS conditions (Figure 2C).

##### Cross-modal ISC

As shown in Fig. 2D, the results highlighted a set of regions across a large extent of the cortex. Significant synchronized regions under the NP condition included primary sensory areas, such as the VC. The ISC peak was observed in the VC (peak r = 0.2248; MNIxyz 26.0, -64.0, 14.0).

Under the IS condition, significant synchronization was observed not only in primary sensory regions, such as the VC, but also in higher-order cortical areas, including the right supramarginal gyrus (located in PPC) and right middle frontal gyrus (located in DLPFC). The ISC peak for this condition was located in the VC (peak r = 0.1521; MNIxyz 26.0, -62.0, 20.0).

While both conditions exhibited the VC as the site of the ISC peak, the IS condition additionally involved significant synchronization in regions located in the DLPFC and PPC, which were not observed during the NP condition Neural-behavioral correlations

#### Experiment 2

**AIS:** Three significant correlations were observed. The IS-induced decrease in *P*_understood_ was negatively correlated with the IS-induced increase in DLPFC activity (Fig. 7A; *r* = -0.464, *p*(FDR) = 0.032), and positively correlated with the IS-induced decrease in STC activity (*r* = 0.440, *p*(FDR) = 0.036). In addition, the IS-induced decrease in *ACC*_objective_ was positively correlated with the IS-induced decrease in STC activity (Fig. 7A; *r* = 0.510, *p*(FDR) = 0.027). This indicates that participants who experienced a greater IS-induced decrease in *P*_understood_ (indicating a more effective IS) also exhibited a greater IS-induced increase in DLPFC activity and a greater IS-induced decrease in STC activity. Participants who experienced a greater IS-induced decrease in *ACC*_objective_ (indicating a more effective IS) also exhibited a greater IS-induced decrease in STC activity (Extended data1-Figure1C, Table 9).

We also computed two-tailed Pearson correlations between IS-related changes in behavioral performance, and the post-3s vs. pre-3s difference of IS-related activation or deactivation. The IS-induced decrease in *P*_understood_ was positively correlated with post-3s vs. pre-3s difference in IS-induced STC deactivation (*r* = 0.528, *p*(FDR) = 0.018). This indicates that participants who experienced a greater IS-induced decrease in *P*_understood_ (indicating a more effective IS) exhibited a more prominent STC low-to-high deactivation pattern (Extended data1-Figure1J, Table 10).

**Table 10.** Correlations between participants’ IS-related behavioral efficacy and the post-3s vs. pre-3s differences of IS-related activation or deactivation (from Experiment 1).

| Items | *r* | *p* | *p*(FDR)<sup>a</sup> |
| --- | --- | --- | --- |
| **AIS** |  |  |  |
| IS-decreased $P_{\text{sensed}}$ & DLPFC high-to-low activation | 0.125 | 0.495 | 0.742 |
| IS-decreased $P_{\text{sensed}}$ & PPC high-to-low activation | 0.055 | 0.763 | 0.957 |
| IS-decreased $P_{\text{sensed}}$ & STC low-to-high deactivation | 0.373 | 0.036 | 0.162 |
| IS-decreased $P_{\text{understood}}$ & DLPFC high-to-low activation | 0.156 | 0.393 | 0.707 |
| IS-decreased $P_{\text{understood}}$ & PPC high-to-low activation | -0.010 | 0.957 | 0.957 |
| IS-decreased $P_{\text{understood}}$ & STC low-to-high deactivation | 0.528 | 0.002 | 0.018* |
| IS-decreased $ACC_{\text{objective}}$ & DLPFC high-to-low activation | -0.020 | 0.915 | 0.957 |
| IS-decreased $ACC_{\text{objective}}$ & PPC high-to-low activation | -0.191 | 0.294 | 0.662 |
| IS-decreased $ACC_{\text{objective}}$ & STC low-to-high deactivation | 0.225 | 0.216 | 0.648 |
| **VIS** |  |  |  |
| IS-decreased $P_{\text{sensed}}$ & DLPFC high-to-low activation | 0.034 | 0.868 | 0.975 |
| IS-decreased $P_{\text{sensed}}$ & PPC high-to-low activation | 0.247 | 0.214 | 0.482 |
| IS-decreased $P_{\text{sensed}}$ & VC low-to-high deactivation | 0.278 | 0.161 | 0.482 |
| IS-decreased $P_{\text{understood}}$ & DLPFC high-to-low activation | 0.006 | 0.975 | 0.975 |
| IS-decreased $P_{\text{understood}}$ & PPC high-to-low activation | -0.093 | 0.645 | 0.829 |
| IS-decreased $P_{\text{understood}}$ & VC low-to-high deactivation | 0.530 | 0.004 | 0.018* |
| IS-decreased $ACC_{\text{objective}}$ & DLPFC high-to-low activation | -0.113 | 0.573 | 0.829 |
| IS-decreased $ACC_{\text{objective}}$ & PPC high-to-low activation | 0.149 | 0.458 | 0.824 |
| IS-decreased $ACC_{\text{objective}}$ & VC low-to-high deactivation | 0.569 | 0.002 | 0.018* |

<sup>a</sup> Corrected using false discovery rate correction method; \* $p(\text{FDR}) < 0.05$ .

**VIS:** Three significant correlations were observed. the IS-induced decrease in *ACC*_objective_ was negatively correlated with the IS-induced increase in DLPFC and PPC activity (DLPFC: *r* = -0.565, *p*(FDR) = 0.009; PPC: *r* = -0.617, *p*(FDR) = 0.009). In addition, the IS-induced decrease in *P*_understood_ was positively correlated with the IS-induced decrease in VC activity (*r* = 0.458, *p*(FDR) = 0.048). This indicates that participants who experienced a greater IS-induced decrease in *ACC*_objective_ (indicating a more effective IS) also exhibited a greater IS-induced increase in DLPFC and PPC activity, and that participants who experienced a greater IS-induced decrease in *P*_understood_ (indicating a more effective IS) also exhibited a greater IS-induced decrease in VC activity (Extended data1-Figure1D, Table 9).

In addition, we also computed two-tailed Pearson correlations between IS-related changes in behavioral performance, and the post-3s vs. pre-3s difference of IS-related activation or deactivation. Both the IS-induced decreases in *P*_understood_ and *ACC*_objective_ were positively correlated with post-3s vs. pre-3s difference in IS-induced VC deactivation (*P*_understood_: Fig. 7A; *r* = 0.530, *p*(FDR) = 0.018; *ACC*_objective_: Fig. 7A; *r* = 0.569, *p*(FDR) = 0.018). This indicates that participants who experienced a greater IS-induced decrease in *P*_understood_ and *ACC*_objective_ (indicating a more effective IS) exhibited a more prominent VC low-to-high deactivation pattern (Extended data1- Figure1K, Table 10).

#### Experiment 3

Based on the robust finding of the relationship between IS-related behavioral changes and the difference in IS-related sensory region deactivation between post-3s and pre-3s across modalities, we specifically tested this correlation in Experiment 3. Positive correlations were observed across all three TMS groups, but only the lDLPFC-activated (*r* = 0.730, *p*(FDR) = 0.015) and rDLPFC-activated (*r* = 0.662, *p*(FDR) = 0.028) groups reached significance, while the sham group showed a marginally significant correlation (*r* = 0.556, *p*(FDR) = 0.061). Fisher’s *r*-to-*z* transformation revealed no significant difference in the correlation coefficients between the lDLPFC-activated group and the sham group (*z* = 0.66, *p* = 0.255), as well as between the rDLPFC-activated group and the sham group (*z* = 0.36, *p* = 0.359; Figure 4D, Table 11).

**Table 11.** Correlations between participants’ IS-related behavioral efficacy and the late vs. early differences of IS-related activation or deactivation, across the three TMS groups.

|  |  |  |  |
| --- | --- | --- | --- |
| Items: IS-decreased $P_{\text{understood}}$ & STC low-to-high deactivation | $r$ | $p$ | $p(\text{FDR})^a$ |
| Left DLPFC-activated group | 0.730 | 0.005 | 0.015* |
| Right DLPFC-activated group | 0.662 | 0.019 | 0.028* |
| Sham group | 0.556 | 0.061 | 0.061 |

<sup>a</sup> Corrected using false discovery rate correction method; \* $p(\text{FDR}) < 0.05$ .

## Acknowledgements

The authors greatly and sincerely thank Professor Trevor Robbins for his guidance on the idea formation, Conceptual clarification and research logic of this research; The author thank Baoxun Chen and Rui Wang for help with data collection, Yunkai Yang and Anhui Kong for help with the picture draft and data processing. This research was supported by supported by the Key Program of the National Natural Science Foundation of China (31920103009) and the Young Scientists Fund of the National Natural Science Foundation of China (32100855).

## Author contributions

Z. H. conceived the study and designed the experiments; Y. Z. conducted the research; Z. H., R. E., Y. D., Z. F., Y. Z. performed data analyses; Z. H. wrote the initial draft of the manuscript; N. M., R. E. and B. S. edited and reviewed the final manuscript. Z. H. acquired funding; B. S. and R. E. supervised the project.

## Completing Interest

The authors declare no competing interests.

## Extended data 1: Figures

**Extended data Figure.1.**
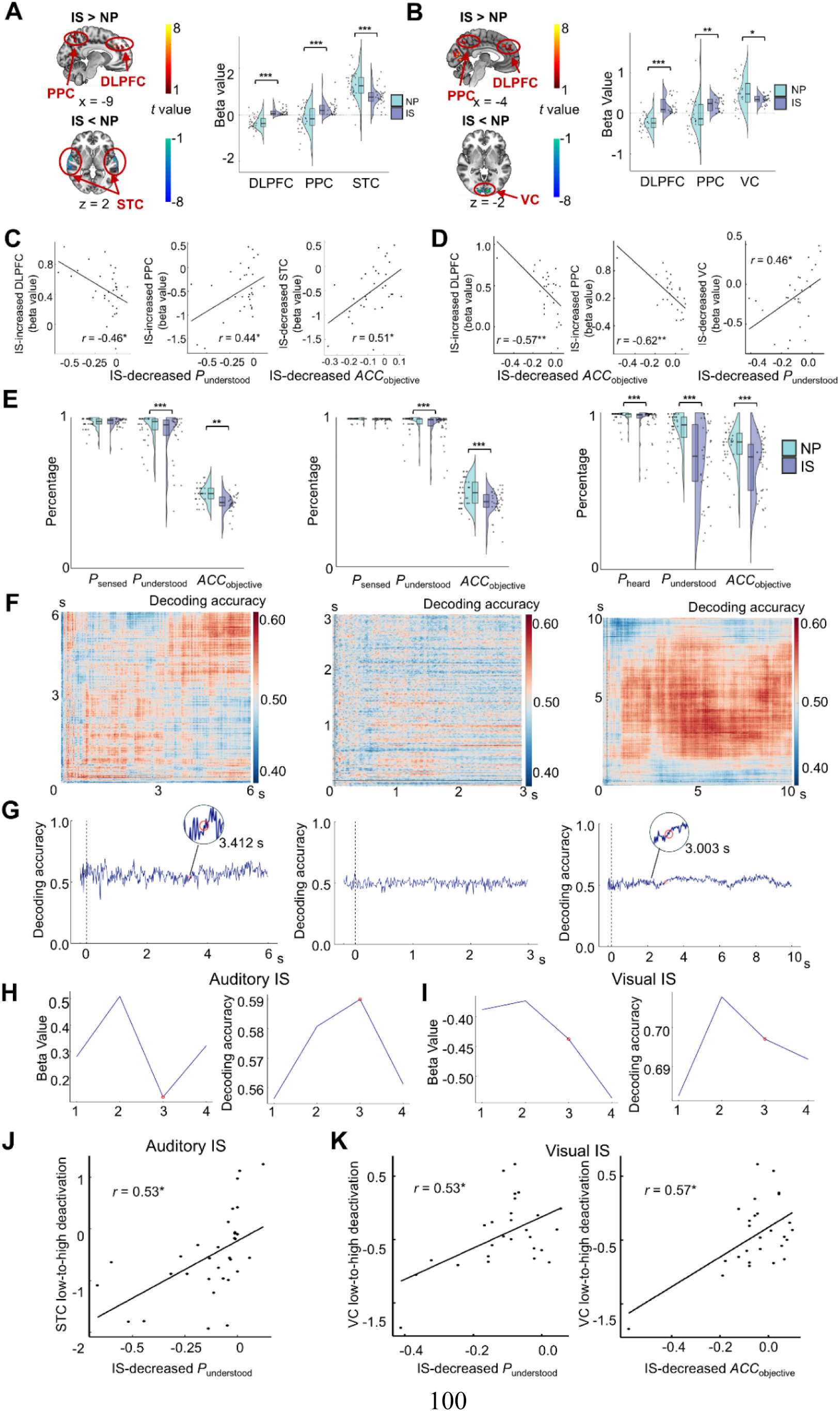
Additional results of brain responses during IS. **A**, fMRI results of the AIS experiment. Brain activation and deactivation maps showing the differences between IS and NP conditions in the predefined ROIs (DLPFC, PPC, STC). Rain cloud plots represent the distribution of beta values. **B,** fMRI results of the VIS experiment. Brain activation and deactivation maps showing the differences between IS and NP conditions in the predefined ROIs (DLPFC, PPC, VC). Rain cloud plots represent the distribution of beta values. **C-D,** Correlations between IS-related changes in behavioral performance and fMRI activation in AIS experiment (C) and VIS experiment (D). **E**, Behavioral performance across 6-second (left panel), 3-second (middle panel), and 10-second (right panel) of voice stimuli. Raincloud plots represent the distribution of responses for *P*_sensed_, *P*_understood_ and *ACC*_objective_, comparing NP with IS. **F,** Temporal decoding result for the IS condition across 6-second (left panel), 3- second (middle panel), and 10-second (right panel) voice stimuli. The classifier was trained on the NP condition and tested for its decodability to the IS condition over time. A higher decoding rate (indicated by red regions) implies greater dissimilarity between the two conditions. **G,** Temporal curve of IS decoding accuracy from the whole-brain EEG data, with the change-point highlighted by a red dot. This curve represents the classifier’s performance in distinguishing IS from NP across 6-second (left panel), 3- second (middle panel), and 10-second (right panel) voice stimuli. **H,** Temporal curve of IS-related brain activation differences (left panel) and the IS-related decoding rates (right panel) in AIS experiment. The temporal curve was derived from the fMRI data using a combined ROI mask (DLPFC, PPC, STC). **I,** Temporal curve of IS-related brain activation differences (left panel) and the IS-related decoding rates (right panel) in VIS experiment. The temporal curve was derived from the fMRI data using a combined ROI mask (DLPFC, PPC, VC). **J-K**, The correlation between IS-related changes in behavioral performance and fMRI activation differences (post3s > pre 3s) in AIS experiment (J) and VIS experiment (K).

**Extended data Figure.2.**
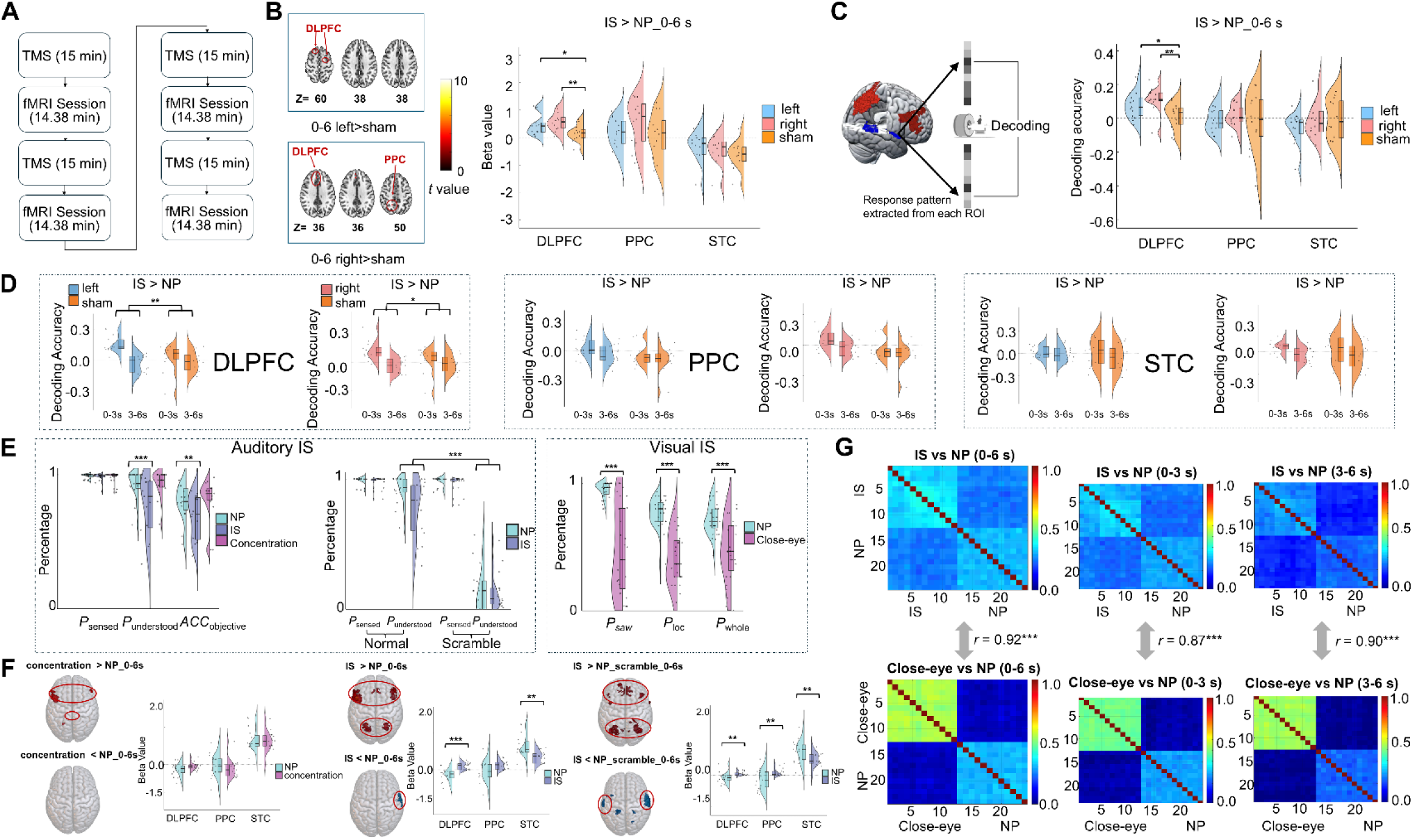
Additional results of the link between IS and brain deactivation patterns. A, TMS procedure. **B**, Brain activation differences (IS > NP) during the 6-second period, comparing lDLPFC-activated vs. sham (top panel) and rDLPFC-activated vs. sham groups (bottom panel). Brain maps display activation and deactivation. Raincloud plots depict beta values comparing IS and NP conditions in each ROI during the 6-second period, across the three TMS groups. **C**, Differences in decoding performance (IS > NP) during the 6-second period across three TMS groups. Raincloud plots depict beta values comparing IS and NP conditions in each ROI during the 6-second period, across the three TMS groups. **D,** ROI decoding accuracy differences (IS > NP) during early (0-3s) and late (3-6s) periods across the three TMS groups. Raincloud plots depict decoding accuracy differences (IS > NP) of each ROI, segmented into early and late periods. **E,** Behavioral results of the “Concentration” model for AIS (left panel), the “Scramble” model for AIS (middle panel), and “Close-eye” model for VIS (right panel). **F,** Brain activation and deactivation differences for the following comparisons during the whole 0-6s periods: concentration vs. NP (left panel), IS vs. NP (middle panel), IS vs. NP-scramble (right panel). Brain maps display activation and deactivation, and raincloud plots depict beta values, segmented into early and late periods. **G,** Correlations between neural RSMs for the following contrasts across different time windows (0-6s, 0-3s, 3-6s) in VIS experiment: IS vs. NP and close-eye vs. NP.

## Extended data 2: Shielding strategy instructions

**Concentration:** Voice/Picture will be presented shortly. Please concentration carefully and attentively to the sounds/pictures that follow.

**Natural processing**^59^: Voice/Picture will be presented shortly. Please relax and respond naturally.

**Inhibition**^60^: Voice/Picture will be presented shortly. Please do not think about any thoughts, feelings, or sensations you might have about this voice/picture. You should not think about any experiences mentioned in the voice/picture. Try to avoid thinking about or feeling anything to do with this voice/picture or the entire process. Even if you start to notice the voice/picture, exclude those thoughts from your mind and stop thinking about them. Not thinking about the voice/picture is your primary task. It is important that you do not think about the voice/picture, or anything related to this task. **Distraction**^60^: Voice/Picture will be presented shortly. Your task is to distract your attention away from any thoughts, feelings, or sensations about the voice/picture. You should do this by thinking about the different rooms in your home. Imagine your home, room by room, as much as you can. Picture the colors of the walls, furniture, photos on the wall, making each room as vivid as possible. Imagine yourself being there, occupy your thoughts, and imagine as many images, scenes, sounds, and activities as you can. As vividly and in as much detail as possible. Even if your thoughts drift elsewhere, bring them immediately back to thoughts about your home. Forming a mental image of your home is your primary task. It is important that you continue imagining your home in vivid detail.

**Distancing**^61^: Voice/Picture will be presented shortly. Imagine your soul leaving your body and observing everything objectively. Please take a third-person perspective of standing aloof from this matter, and observe it like a bystander.

## Extended data 3: Experimental materials

To facilitate direct comparisons between the ratings of stimuli across the four task conditions—in terms of valence, arousal, processing difficulty, d information load—we conducted mixed-effects ANOVAs, treating participants as random effects. For the AIS experiment, the assigned *task ndition* (NP, inhibition, distraction, distancing) and *voice duration* (3 seconds, 6 seconds, and 10 seconds) were included as within-subject factors. or the VIS experiment, the assigned *task condition* was included as the within-subject factor. For the AIS experiment, an independent sample of 28 college students (16 females; 20.04 ± 1.77, *M* ± *SD*), demographically matched to the ain study participants, rated the stimuli using nine-point scales. Valence was rated from 1 (most negative) to 9 (most positive); arousal from 1 east arousing) to 9 (most arousing); information load from 1 (lowest) to 9 (highest); and processing difficulty from 1 (least difficult) to 9 (most fficult). Results revealed no significant effects of task condition on any dimension for either 3-second [valence: *F*(3, 81) = 0.65, *p* = 0.588, *η²* = 018; arousal: *F*(3, 81) = 0.31, *p* = 0.818, *η²* = 0.009; information load: *F*(3, 81) = 0.04, *p* = 0.988, *η²* = 0.001; processing difficulty: *F*(3, 81) = 26, *p* = 0.857, *η²* = 0.007], 6-second [valence: *F*(3, 81) = 0.55, *p* = 0.647, *η²* = 0.015; arousal: *F*(3, 81) = 1.24, *p* = 0.298, *η²* = 0.034; information ad: *F*(3, 81) = 0.46, *p* = 0.708, *η²* = 0.013; processing difficulty: *F*(3, 81) = 1.02, *p* = 0.388, *η²* = 0.028], or 10-second [valence: *F*(3, 81) = 0.36, = 0.781, *η²* = 0.010; arousal: *F*(3, 81) = 1.53, *p* = 0.210, *η²* = 0.041; information load: *F*(3, 81) = 0.03, *p* = 0.994, *η²* = 0.001; processing difficulty: (3, 81) = 1.59, *p* = 0.197, *η²* = 0.043] clips. Nevertheless, a main difference in all dimension ratings was found across the 3-, 6-, and 10- second ration stimulus (valence: 3-second < 6-second < 10-second, *F*(2, 297) = 68.258, *p* < 0.001, *η²* = 0.31; arousal: 3-second > 6-second > 10-second, (2, 297) = 68.26, *p* < 0.001, *η²* = 0.27; information load: 3-second < 6-second < 10-second, *F*(2, 297) = 66.57, *p* < 0.001, *η²* = 0.31; processing fficulty: 3-second < 6-second < 10-second, *F*(2, 297) = 29.94, *p* < 0.001, *η²* = 0.17).

For the VIS experiment, an independent sample of 30 college students (15 females; 21.77 ± 2.01), demographically matched to the main study rticipants, rated the stimuli using nine-point scales (including valence, arousal, information load, and processing difficulty). Results revealed no gnificant effects of *task condition* on any dimension [valence: *F*(3, 87) = 1.51, *p* = 0.219, *η²* = 0.050; arousal: *F*(3, 87) = 0.44, p = 0.728, *η²* = 010; information load: *F*(3, 87) = 0.053, *p* = 0.984, *η²* = 0.001; processing difficulty: *F*(3, 87) = 0.217, *p* = 0.884, *η²* = 0.010] All word texts of the video clips, the ID of the IAPS pictures, quizzes, descriptive statistics for average ratings as well as average rating for each dividual stimulus can be found in the **Extended data** Table 1 **and 2**.

**Extended data Table 1.** Participants’ ratings of study materials (*M* ± *SD*) grouped by evaluation dimensions for **(a)** AIS experiment, including ch assigned task condition and voice length, and **(b)** VIS experiment.

|  | 3-second voice |  |  |  | 6-second voice |  |  |  | 10-second voice |  |  |  |
| --- | --- | --- | --- | --- | --- | --- | --- | --- | --- | --- | --- | --- |
|  | NP | Inhibition | Distraction | Distancing | NP | Inhibition | Distraction | Distancing | NP | Inhibition | Distraction | Distancing |
| Valence | 3.39 ± 0.54 | 3.22 ± 0.49 | 3.27 ± 0.52 | 3.37 ± 0.56 | 3.52 ± 0.65 | 3.57 ± 0.66 | 3.71 ± 0.56 | 3.66 ± 0.57 | 4.03 ± 0.72 | 3.90 ± 0.72 | 4.00 ± 0.67 | 4.08 ± 0.68 |
| Arousal | 6.03 ± 0.75 | 6.02 ± 0.72 | 5.93 ± 0.60 | 5.88 ± 0.73 | 5.62 ± 0.74 | 5.79 ± 0.73 | 5.45 ± 0.66 | 5.87 ± 0.54 | 4.85 ± 0.99 | 5.36 ± 0.86 | 4.99 ± 1.05 | 4.96 ± 0.92 |
| Information load | 2.94 ± 1.52 | 2.89 ± 1.48 | 2.87 ± 1.56 | 3.00 ± 1.55 | 3.90 ± 1.14 | 3.66 ± 1.10 | 3.66 ± 1.15 | 3.91 ± 1.13 | 4.47 ± 1.42 | 4.51 ± 1.29 | 4.54 ± 1.28 | 4.56 ± 1.31 |
| Processing difficulty | 2.63 ± 1.23 | 2.85 ± 1.11 | 2.86 ± 1.10 | 2.87 ± 1.24 | 2.63 ± 1.23 | 2.94 ± 1.13 | 3.48 ± 1.28 | 3.09 ± 1.27 | 3.57 ± 1.19 | 3.31 ± 1.29 | 3.98 ± 1.12 | 3.63 ± 1.27 |

|  | NP | Inhibition | Distraction | Distancing |
| --- | --- | --- | --- | --- |
| Valence | $5.56 \pm 0.40$ | $5.48 \pm 0.40$ | $5.54 \pm 0.43$ | $5.62 \pm 0.32$ |
| Arousal | $5.09 \pm 0.59$ | $5.08 \pm 0.62$ | $4.99 \pm 0.61$ | $5.04 \pm 0.64$ |
| Information load | $4.14 \pm 1.06$ | $4.14 \pm 1.03$ | $4.16 \pm 1.04$ | $4.16 \pm 1.11$ |
| Processing difficulty | $2.89 \pm 1.13$ | $2.906 \pm 1.08$ | $2.893 \pm 1.08$ | $2.937 \pm 1.11$ |

Extended data Table 2. All word texts of the video clips, the IDs of the IAPS pictures, quizzes, and the average rating for each individual stimulus ecorded in a separated file).

## Extended data 4: Descriptive statistics of the Behavioural results

Descriptive statistics for *P*_sensed_, *P*_understood_, and *ACC*_objective_ are presented in Extended data Table 3.

**Extended data Table 3.**
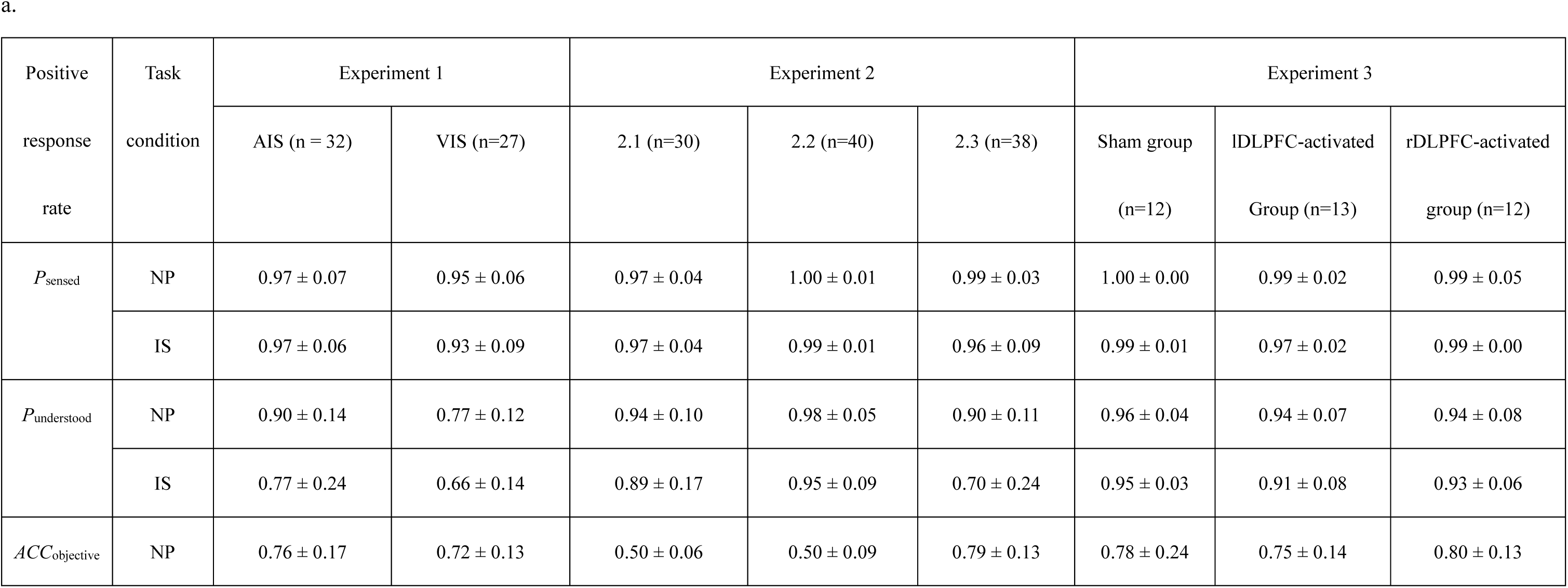
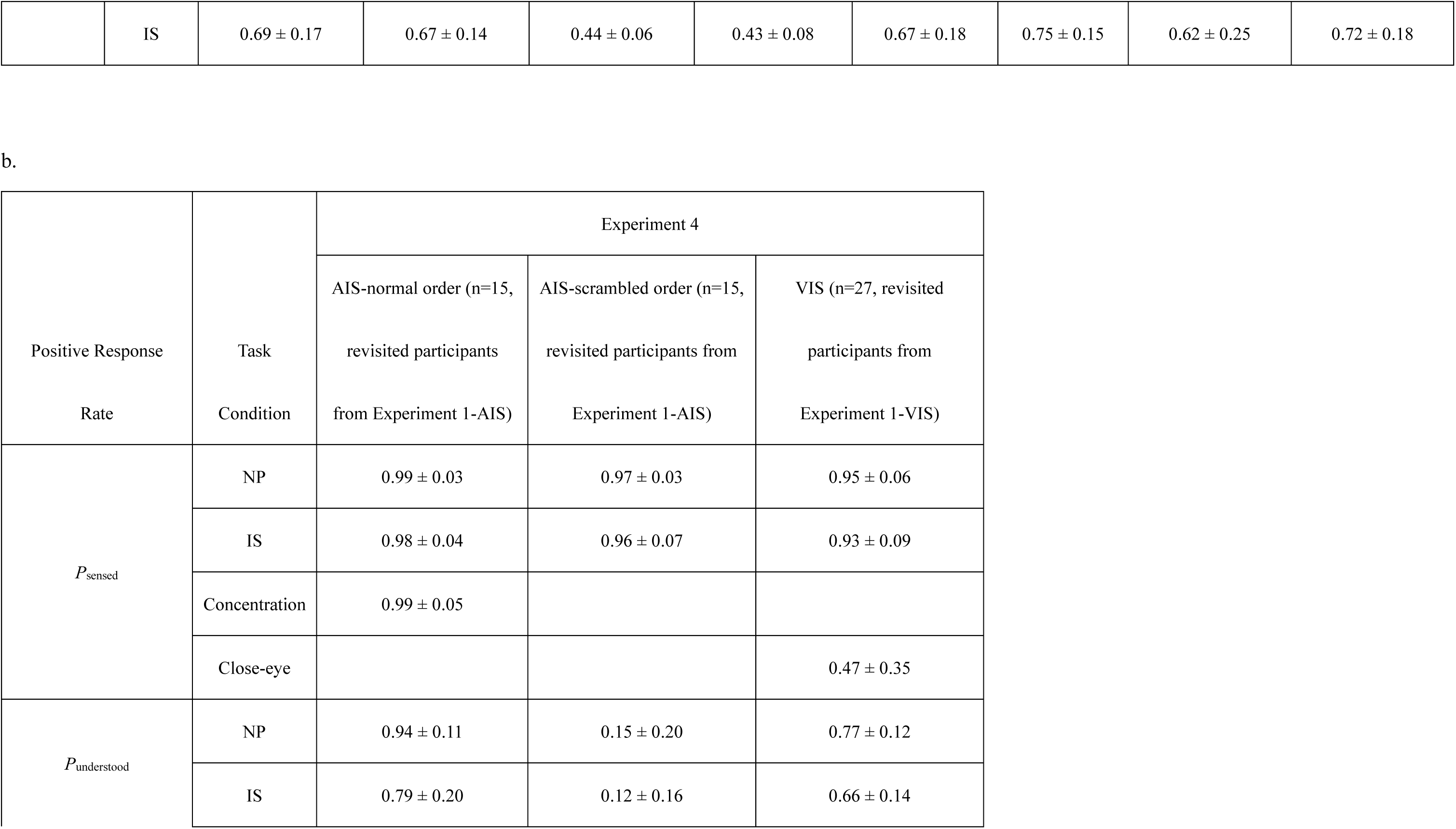
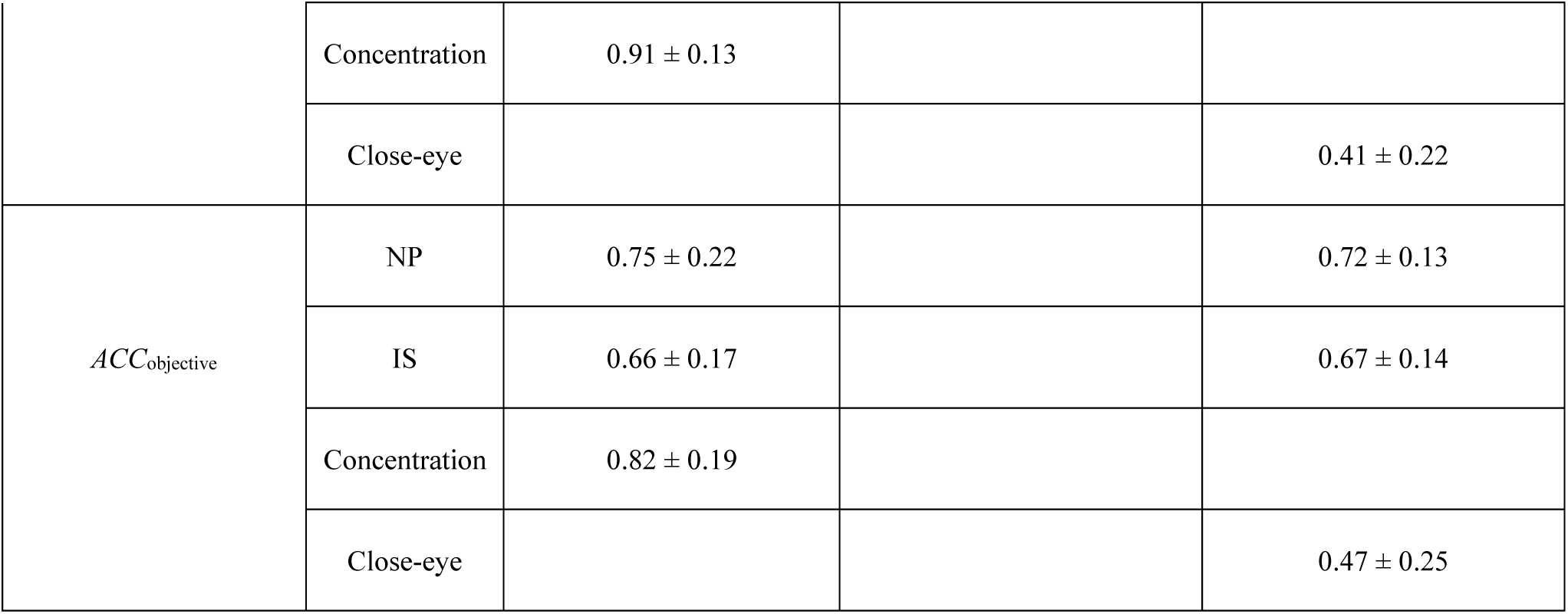
Descriptive statistics (M ± SD) of each type of responses under different conditions across all participants in Experiments 2, 3 **(a)** and Experiment 4 **(b)**. All rates were taken as the average positive response under each condition. Note: Response rate data of NP and under normal semantic order condition in Experiment 4 was extracted from the same group of participants’ first visit in Experiment 1.

## Extended data 5: Clusters showing the significant brain activations within the ROI

**Extended data Table 4.** Significant activation clusters within the combined ROI (DLPFC, PPC, STC/VC) in Experiment 1 (including the AIS and VIS parts): Analysis of voice exposure periods **(a)** 0-6 seconds, and **(b)** difference of post-3s vs pre-3s period.

| Region | Laterality | Cluster size<br>(Voxels) | T | Z | $p(\text{FDR})$ | Peak MNI coordinates | | |
| --- | --- | --- | --- | --- | --- | --- | --- | --- |
|  |  |  |  |  |  | x | y | z |
| AIS, Contrast: IS > NP |  |  |  |  |  |  |  |  |
| DLPFC | L | 709 | 11.10 | 7.00 | < 0.001 | -46 | 4 | 30 |
|  | R | 181 | 8.49 | 6.06 | < 0.001 | 8 | 30 | 38 |
|  | L | 289 | 8.06 | 5.87 | < 0.001 | -8 | 26 | 36 |
|  | R | 773 | 7.56 | 5.65 | < 0.001 | 44 | 10 | 30 |
|  | L | 16 | 5.45 | 4.53 | < 0.001 | -50 | 38 | 2 |
| PPC | L | 693 | 9.70 | 6.53 | < 0.001 | -24 | -62 | 44 |
|  | R | 667 | 8.03 | 5.86 | < 0.001 | 32 | -62 | 44 |
|  | L | 11 | 5.50 | 4.56 | < 0.001 | -26 | -70 | 36 |
| AIS, Contrast: IS < NP |  |  |  |  |  |  |  |  |
| STC | R | 289 | 9.03 | 6.28 | < 0.001 | 62 | -4 | 0 |
|  | L | 100 | 7.40 | 5.58 | < 0.001 | -60 | -8 | 2 |

**VIS, Contrast: IS > NP**

|  |  |  |  |  |  |  |  |  |
| --- | --- | --- | --- | --- | --- | --- | --- | --- |
| **DLPFC** | L | 782 | 10.24 | 6.43 | < 0.001 | -44 | 18 | 26 |
|  | R | 192 | 8.70 | 5.90 | < 0.001 | 8 | 30 | 38 |
|  | R | 750 | 7.16 | 5.28 | < 0.001 | 44 | 20 | 34 |
|  | L | 191 | 5.56 | 4.47 | < 0.001 | -2 | 32 | 38 |
| **PPC** | L | 810 | 7.98 | 5.63 | < 0.001 | -26 | -54 | 44 |
|  | R | 388 | 5.74 | 4.57 | < 0.001 | 36 | -62 | 46 |
|  | R | 116 | 4.13 | 3.59 | 0.002 | 10 | -66 | 48 |
|  | L | 15 | 2.55 | 2.39 | 0.035 | -14 | -70 | 32 |
| **VC** | L | 669 | 11.15 | 6.70 | < 0.001 | 4 | -70 | 2 |
|  | L | 73 | 6.33 | 4.88 | < 0.001 | -32 | -96 | -12 |
|  | L | 58 | 4.20 | 3.63 | 0.002 | -36 | -88 | -14 |
|  | L | 10 | 2.98 | 2.74 | 0.017 | -20 | -58 | 2 |

**VIS, Contrast: IS < NP**

|  |  |  |  |  |  |  |  |  |
| --- | --- | --- | --- | --- | --- | --- | --- | --- |
| **VC** | L | 104 | 10.13 | 6.39 | < 0.001 | -10 | -94 | 0 |
|  | L | 262 | 9.45 | 6.17 | < 0.001 | -16 | -94 | 14 |
|  | R | 446 | 9.45 | 6.17 | < 0.001 | 4 | -86 | -8 |
|  | L | 118 | 7.74 | 5.41 | < 0.001 | -22 | -78 | -16 |
|  | R | 124 | 7.28 | 5.33 | < 0.001 | 30 | -90 | 16 |
|  | R | 28 | 6.63 | 5.03 | < 0.001 | 34 | -86 | 4 |
|  | R | 11 | 5.69 | 4.54 | < 0.001 | 34 | -74 | -18 |
|  | R | 13 | 4.96 | 4.13 | 0.001 | 52 | -60 | -10 |
|  | L | 13 | 3.74 | 3.32 | 0.007 | -26 | -86 | 26 |
|  | R | 10 | 3.55 | 3.17 | 0.010 | 42 | -80 | 4 |
| **DLPFC** | L | 34 | 5.76 | 4.58 | < 0.001 | -22 | 52 | 42 |

| Region | Laterality | Cluster size<br>(Voxels) | T | Z | $p(\text{FDR})$ | Peak MNI coordinates | | |
| --- | --- | --- | --- | --- | --- | --- | --- | --- |
|  |  |  |  |  |  | x | y | z |
| AIS, Contrast: IS > NP 0-3s |  |  |  |  |  |  |  |  |
| DLPFC | L | 648 | 10.74 | Inf | < 0.001 | -50 | 24 | 26 |
|  | R | 147 | 8.91 | 7.81 | < 0.001 | 4 | 30 | 38 |
|  | L | 144 | 8.78 | 7.72 | < 0.001 | -2 | 32 | 38 |
|  | R | 705 | 8.07 | 7.22 | < 0.001 | 52 | 28 | 34 |
| PPC | L | 762 | 7.60 | 6.87 | < 0.001 | -24 | -62 | 44 |
|  | R | 638 | 6.61 | 6.11 | < 0.001 | 28 | -58 | 44 |
|  | L | 15 | 5.67 | 5.34 | < 0.001 | -26 | -70 | 36 |
| AIS, Contrast: IS > NP 3-6s |  |  |  |  |  |  |  |  |
| DLPFC | L | 546 | 10.14 | Inf | < 0.001 | -42 | 2 | 30 |
|  | R | 602 | 7.66 | 6.92 | < 0.001 | 44 | 10 | 30 |
|  | R | 113 | 7.48 | 6.78 | < 0.001 | 8 | 30 | 38 |
|  | L | 302 | 6.02 | 5.62 | < 0.001 | -8 | 26 | 36 |
|  | L | 11 | 4.52 | 4.34 | < 0.001 | -50 | 38 | 2 |
| **PPC** | L | 468 | 9.68 | Inf | < 0.001 | -24 | -60 | 44 |
|  | R | 438 | 7.08 | 6.48 | < 0.001 | 28 | -58 | 44 |
|  | L | 11 | 5.55 | 5.23 | < 0.001 | -26 | -70 | 36 |
|  | R | 13 | 3.20 | 3.13 | 0.005 | 12 | -66 | 46 |

**AIS, Contrast: Post 3s (IS > NP) – Pre 3s (IS > NP)**

**AIS, Contrast: IS < NP 0-3s**
|  |  |  |  |  |  |  |  |  |
| --- | --- | --- | --- | --- | --- | --- | --- | --- |
| **STC** | R | 177 | 6.31 | 5.87 | < 0.001 | 62 | -2 | -2 |
|  | L | 126 | 6.10 | 5.70 | < 0.001 | -62 | -4 | 0 |

**AIS, Contrast: IS < NP 3-6s**

|  |  |  |  |  |  |  |  |  |
| --- | --- | --- | --- | --- | --- | --- | --- | --- |
| **STC** | R | 481 | 9.59 | Inf | < 0.001 | 62 | -4 | 0 |
|  | L | 340 | 8.25 | 7.35 | < 0.001 | -62 | -12 | 4 |
| **PPC** | R | 25 | 3.97 | 3.85 | 0.001 | 6 | -80 | 42 |
|  | L | 16 | 3.54 | 3.45 | 0.004 | -4 | -78 | 34 |

**AIS, Contrast: Post 3s (IS < NP) – Pre 3s (IS < NP)**

|  |  |  |  |  |  |  |  |  |
| --- | --- | --- | --- | --- | --- | --- | --- | --- |
| **DLPFC** | L | 62 | 3.93 | 3.80 | 0.007 | -46 | 24 | 42 |
|  | R | 51 | 3.77 | 3.66 | 0.010 | 54 | 26 | 34 |
|  | L | 98 | 3.62 | 3.52 | 0.013 | -54 | 28 | 30 |
| **PPC** | L | 113 | 4.52 | 4.34 | 0.003 | -28 | -42 | 68 |
|  | L | 16 | 3.19 | 3.12 | 0.026 | -42 | -64 | 50 |
|  | L | 12 | 2.96 | 2.90 | 0.038 | -12 | -58 | 66 |
| **STC** | R | 53 | 4.99 | 4.75 | 0.003 | 50 | -16 | 6 |
|  | L | 29 | 4.85 | 4.64 | 0.003 | -48 | -18 | 6 |
|  | R | 123 | 4.29 | 4.14 | 0.005 | 58 | -10 | 2 |
| **VIS, Contrast: IS > NP 0-3s** |  |  |  |  |  |  |  |  |
| **DLPFC** | R | 889 | 8.07 | 5.66 | <0.001 | 38 | 6 | 38 |
|  | R | 229 | 7.83 | 5.56 | <0.001 | 6 | 30 | 38 |
|  | L | 810 | 7.38 | 5.37 | <0.001 | -48 | 16 | 28 |
|  | L | 151 | 4.80 | 4.03 | 0.001 | -2 | 32 | 38 |
|  | L | 13 | 2.83 | 2.62 | 0.018 | -42 | 42 | 10 |
| **PPC** | L | 2287 | 6.68 | 5.05 | <0.001 | -34 | -62 | 48 |
|  | R | 14 | 2.51 | 2.35 | 0.032 | 4 | -42 | 52 |
|  | R | 14 | 2.39 | 2.25 | 0.039 | 10 | -52 | 52 |
|  | R | 21 | 2.26 | 2.14 | 0.048 | 14 | -46 | 74 |
| **VC** | L | 75 | 6.55 | 4.99 | <0.001 | -32 | -96 | -12 |
|  | R | 302 | 6.02 | 4.72 | <0.001 | 42 | -68 | 48 |
|  | L | 91 | 4.44 | 3.80 | 0.001 | -36 | -88 | -14 |
| **VIS, Contrast: IS > NP 3-6s** |  |  |  |  |  |  |  |  |
| **DLPFC** | L | 631 | 9.81 | 6.29 | <0.001 | -44 | 18 | 26 |
|  | R | 87 | 6.73 | 5.08 | <0.001 | 8 | 30 | 38 |
|  | R | 441 | 6.50 | 4.97 | <0.001 | 44 | 22 | 34 |
|  | L | 130 | 4.95 | 4.12 | 0.001 | -8 | 26 | 36 |
|  | L | 13 | 3.9 | 3.48 | 0.004 | -50 | 38 | 2 |
|  | R | 18 | 3.57 | 3.19 | 0.008 | 48 | 20 | 28 |
| **PPC** | L | 407 | 6.41 | 4.92 | <0.001 | -26 | -54 | 44 |
|  | R | 221 | 5.48 | 4.43 | <0.001 | 32 | -46 | 64 |
|  | R | 48 | 4.30 | 3.70 | 0.002 | 10 | -66 | 48 |
|  | L | 53 | 3.82 | 3.37 | 0.005 | -32 | -48 | 62 |
|  | L | 21 | 3.21 | 2.92 | 0.015 | -30 | -74 | 42 |
| **VC** | R | 186 | 10.05 | 6.37 | <0.001 | 4 | -70 | 0 |
|  | L | 195 | 8.24 | 5.73 | <0.001 | -4 | -74 | 2 |
|  | L | 42 | 4.48 | 3.82 | 0.001 | -32 | -96 | -12 |
| **VIS, Contrast: Post 3s (IS > NP) – Pre 3s (IS > NP)** |  |  |  |  |  |  |  |  |
| **DLPFC** | L | 59 | 7.25 | 6.50 | < 0.001 | -12 | 56 | 42 |
|  | L | 22 | 4.56 | 4.35 | 0.003 | -60 | 6 | 28 |
|  | R | 23 | 4.11 | 3.95 | 0.011 | 32 | -46 | 62 |
| **VC** | R | 15 | 4.00 | 3.84 | 0.014 | 6 | -70 | 2 |
| **VIS, Contrast: IS < NP 0-3s** |  |  |  |  |  |  |  |  |
| **VC** | L | 122 | 8.34 | 7.28 | < 0.001 | -26 | -48 | -8 |
|  | R | 87 | 6.26 | 5.75 | < 0.001 | 30 | -50 | -8 |
| **VIS, Contrast: IS < NP 3-6s** |  |  |  |  |  |  |  |  |
| **VC** | L | 4467 | 9.59 | Inf | < 0.001 | -12 | -82 | 38 |
|  | R | 34 | 6.85 | 6.21 | < 0.001 | 48 | -64 | 12 |
|  | L | 36 | 5.77 | 5.36 | < 0.001 | -46 | -64 | 12 |
|  | R | 37 | 5.63 | 5.25 | < 0.001 | 28 | -64 | -4 |
|  | R | 44 | 5.12 | 4.82 | < 0.001 | 50 | -70 | 6 |
|  | L | 10 | 4.98 | 4.71 | < 0.001 | -14 | -54 | 2 |
|  | R | 15 | 3.11 | 3.03 | 0.004 | 50 | -62 | -6 |
|  | R | 13 | 2.97 | 2.90 | 0.006 | 44 | -74 | -8 |
|  | R | 11 | 2.65 | 2.60 | 0.012 | 38 | -82 | 20 |
| **PPC** | L | 648 | 6.76 | 6.14 | < 0.001 | -14 | -38 | 78 |
|  | R | 375 | 6.66 | 6.06 | < 0.001 | 14 | -42 | 50 |
| **DLPFC** | R | 92 | 5.67 | 5.28 | < 0.001 | 20 | 32 | 32 |
|  | R | 88 | 5.43 | 5.09 | < 0.001 | 50 | 16 | 26 |
|  | R | 52 | 4.83 | 4.57 | < 0.001 | 20 | 36 | 42 |
|  | L | 13 | 4.26 | 4.08 | < 0.001 | -28 | 34 | 40 |
|  | L | 59 | 4.17 | 4.00 | < 0.001 | -18 | 32 | 30 |
|  | R | 123 | 4.14 | 3.97 | < 0.001 | 38 | 28 | 42 |
|  | R | 84 | 4.04 | 3.89 | < 0.001 | 50 | 28 | 18 |
|  | L | 25 | 3.95 | 3.80 | < 0.001 | -24 | 38 | 40 |
|  | R | 16 | 3.88 | 3.74 | < 0.001 | 12 | 48 | 34 |
|  | R | 14 | 3.45 | 3.35 | 0.002 | 18 | 52 | 36 |
| **VIC, Contrast: Post 3s (IS < NP) – Pre 3s (IS < NP)** |  |  |  |  |  |  |  |  |
| **VC** | L | 833 | 6.23 | 5.73 | < 0.001 | -12 | -80 | 42 |
|  | L | 194 | 5.58 | 5.21 | < 0.001 | -26 | -88 | -26 |
|  | R | 199 | 5.21 | 4.90 | < 0.001 | 24 | -88 | -26 |
|  | R | 492 | 4.98 | 4.70 | < 0.001 | 24 | -80 | 40 |
|  | L | 24 | 4.80 | 4.55 | < 0.001 | -16 | -44 | -6 |
|  | L | 37 | 4.68 | 4.45 | 0.001 | -42 | -68 | 44 |
|  | R | 17 | 4.65 | 4.42 | 0.001 | 18 | -44 | -6 |
|  | L | 41 | 3.99 | 3.84 | 0.002 | -46 | -80 | 10 |
|  | R | 16 | 3.79 | 3.66 | 0.004 | 34 | -46 | -10 |
|  | L | 149 | 3.76 | 3.63 | 0.004 | -30 | -98 | 0 |
|  | R | 21 | 3.52 | 3.42 | 0.006 | 50 | -66 | 12 |
|  | R | 46 | 3.51 | 3.40 | 0.006 | 18 | -88 | 12 |
|  | R | 33 | 3.44 | 3.34 | 0.007 | 40 | -66 | 52 |
|  | L | 14 | 3.38 | 3.28 | 0.008 | 2 | -90 | -4 |
|  | L | 11 | 2.85 | 2.79 | 0.020 | -2 | -88 | -4 |
| **PPC** | L | 382 | 5.02 | 4.74 | < 0.001 | -14 | -38 | 78 |
|  | R | 173 | 4.43 | 4.23 | 0.001 | 14 | -42 | 50 |
| **DLPFC** | R | 52 | 4.26 | 4.08 | 0.001 | 52 | 16 | 26 |
|  | R | 50 | 4.22 | 4.04 | 0.001 | 20 | 32 | 32 |
|  | R | 38 | 3.59 | 3.47 | 0.006 | 20 | 36 | 42 |
|  | L | 11 | 3.10 | 3.02 | 0.013 | -18 | 32 | 30 |
|  | R | 29 | 3.07 | 3.00 | 0.014 | 38 | 28 | 42 |
|  | R | 14 | 3.00 | 2.93 | 0.016 | 50 | 28 | 18 |
|  | R | 20 | 2.80 | 2.75 | 0.021 | 50 | 16 | 40 |

Data are thresholded at $p(\text{FDR}) < 0.05$ , with a minimum cluster size of 10 voxels (For all analyses involving multiple
comparisons, the $p$ -values were further adjusted using an additional FDR correction). R: right. L: left.

## Extended data 6: Descriptive statistics of the MVPA results

**Extended data Table 5.**
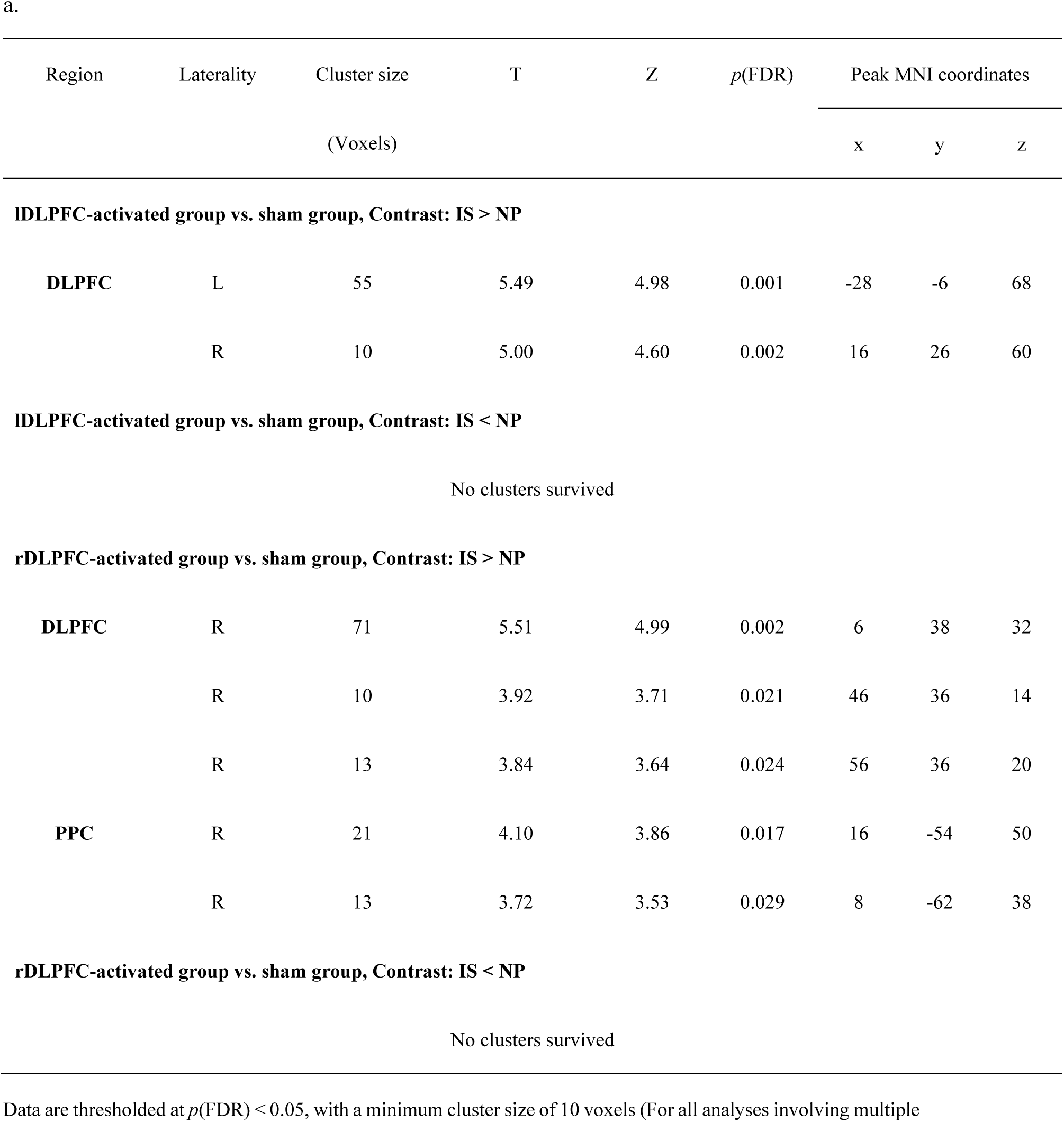
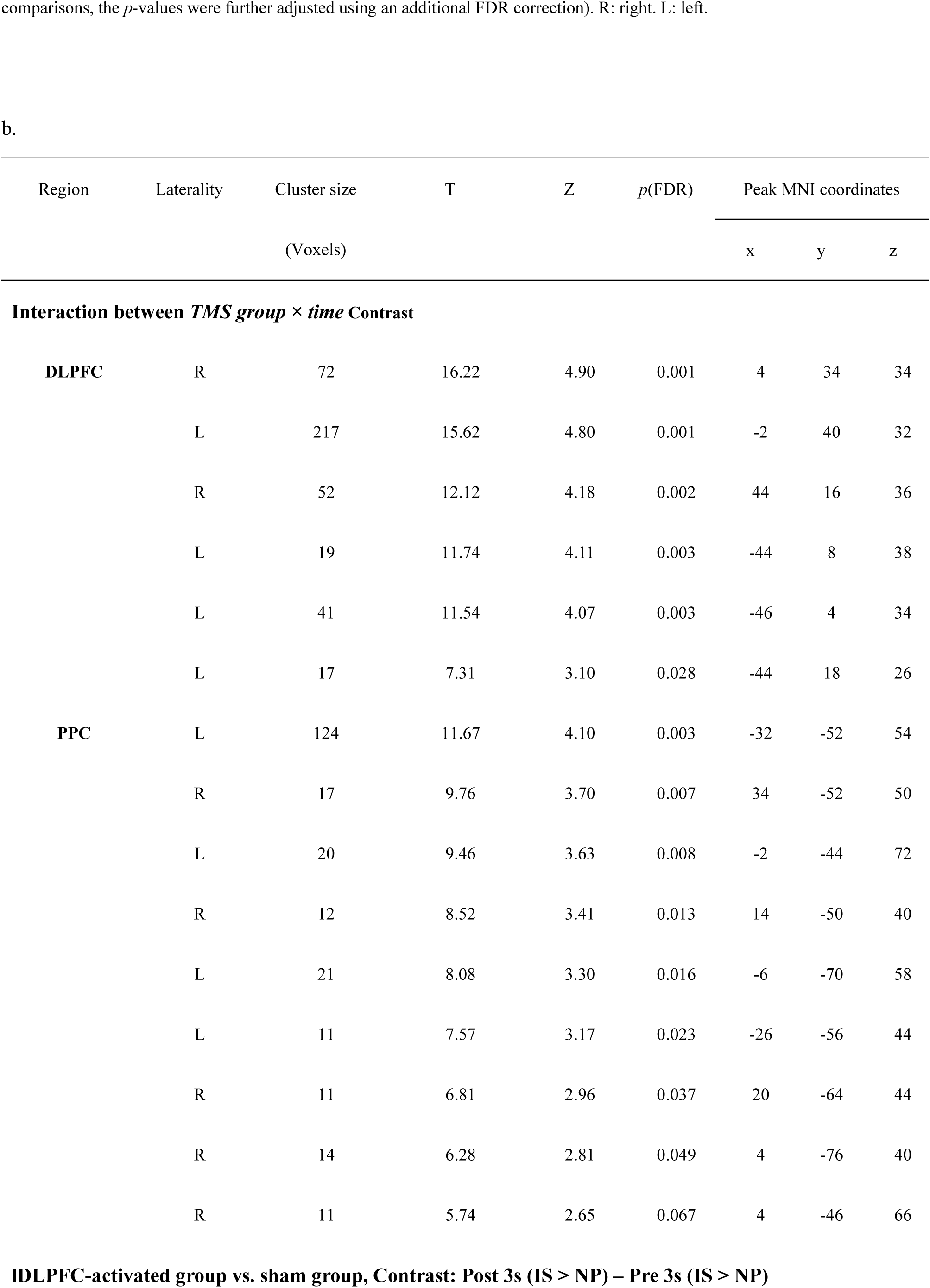
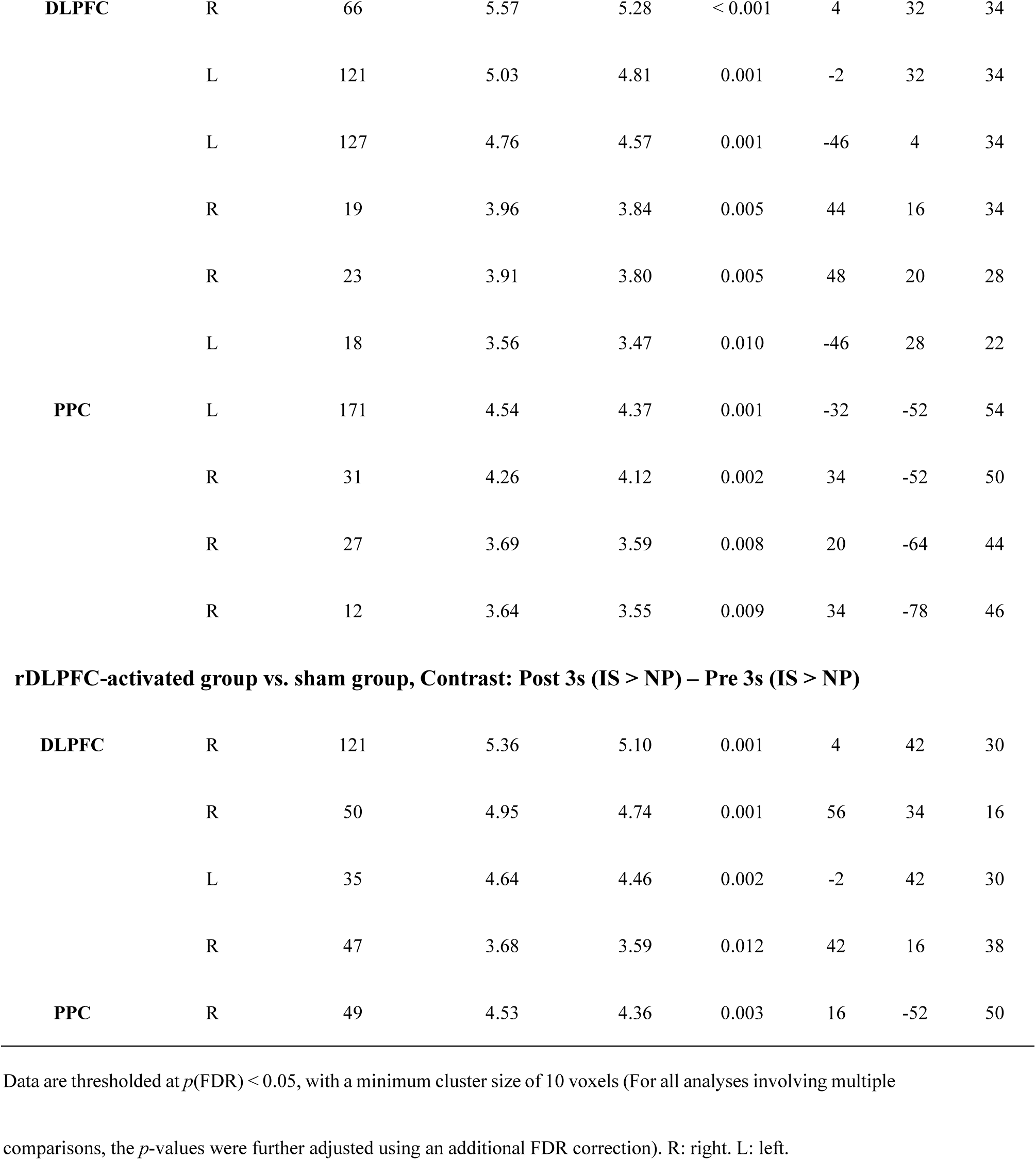
Significant activation clusters within the combined ROI (DLPFC, PPC, STC) in Experiment 3: Analysis of voice exposure periods **(a)** 0-6 seconds, and **(b)** difference of post-3s vs pre-3s period.

| Region | Laterality | Cluster size<br>(Voxels) | T | Z | p(FDR) | Peak MNI coordinates |  |  |
| --- | --- | --- | --- | --- | --- | --- | --- | --- |
|  |  |  |  |  |  | x | y | z |
| IDL PFC-activated group vs. sham group, Contrast: IS > NP |  |  |  |  |  |  |  |  |
| DLPFC | L | 55 | 5.49 | 4.98 | 0.001 | -28 | -6 | 68 |
|  | R | 10 | 5.00 | 4.60 | 0.002 | 16 | 26 | 60 |
| IDL PFC-activated group vs. sham group, Contrast: IS < NP |  |  |  |  |  |  |  |  |
| No clusters survived |  |  |  |  |  |  |  |  |
| rDLPFC-activated group vs. sham group, Contrast: IS > NP |  |  |  |  |  |  |  |  |
| DLPFC | R | 71 | 5.51 | 4.99 | 0.002 | 6 | 38 | 32 |
|  | R | 10 | 3.92 | 3.71 | 0.021 | 46 | 36 | 14 |
|  | R | 13 | 3.84 | 3.64 | 0.024 | 56 | 36 | 20 |
| PPC | R | 21 | 4.10 | 3.86 | 0.017 | 16 | -54 | 50 |
|  | R | 13 | 3.72 | 3.53 | 0.029 | 8 | -62 | 38 |
| rDLPFC-activated group vs. sham group, Contrast: IS < NP |  |  |  |  |  |  |  |  |
| No clusters survived |  |  |  |  |  |  |  |  |

| Region | Laterality | Cluster size<br>(Voxels) | T | Z | $p(\text{FDR})$ | Peak MNI coordinates | | |
| --- | --- | --- | --- | --- | --- | --- | --- | --- |
|  |  |  |  |  |  | x | y | z |
| Interaction between *TMS group* × *time* Contrast | | | | | | | | |
| DLPFC | R | 72 | 16.22 | 4.90 | 0.001 | 4 | 34 | 34 |
|  | L | 217 | 15.62 | 4.80 | 0.001 | -2 | 40 | 32 |
|  | R | 52 | 12.12 | 4.18 | 0.002 | 44 | 16 | 36 |
|  | L | 19 | 11.74 | 4.11 | 0.003 | -44 | 8 | 38 |
|  | L | 41 | 11.54 | 4.07 | 0.003 | -46 | 4 | 34 |
|  | L | 17 | 7.31 | 3.10 | 0.028 | -44 | 18 | 26 |
| PPC | L | 124 | 11.67 | 4.10 | 0.003 | -32 | -52 | 54 |
|  | R | 17 | 9.76 | 3.70 | 0.007 | 34 | -52 | 50 |
|  | L | 20 | 9.46 | 3.63 | 0.008 | -2 | -44 | 72 |
|  | R | 12 | 8.52 | 3.41 | 0.013 | 14 | -50 | 40 |
|  | L | 21 | 8.08 | 3.30 | 0.016 | -6 | -70 | 58 |
|  | L | 11 | 7.57 | 3.17 | 0.023 | -26 | -56 | 44 |
|  | R | 11 | 6.81 | 2.96 | 0.037 | 20 | -64 | 44 |
|  | R | 14 | 6.28 | 2.81 | 0.049 | 4 | -76 | 40 |
|  | R | 11 | 5.74 | 2.65 | 0.067 | 4 | -46 | 66 |
| **IDLPFC-activated group vs. sham group, Contrast: Post 3s (IS > NP) – Pre 3s (IS > NP)** |  |  |  |  |  |  |  |  |
| **DLPFC** | R | 66 | 5.57 | 5.28 | < 0.001 | 4 | 32 | 34 |
|  | L | 121 | 5.03 | 4.81 | 0.001 | -2 | 32 | 34 |
|  | L | 127 | 4.76 | 4.57 | 0.001 | -46 | 4 | 34 |
|  | R | 19 | 3.96 | 3.84 | 0.005 | 44 | 16 | 34 |
|  | R | 23 | 3.91 | 3.80 | 0.005 | 48 | 20 | 28 |
|  | L | 18 | 3.56 | 3.47 | 0.010 | -46 | 28 | 22 |
| **PPC** | L | 171 | 4.54 | 4.37 | 0.001 | -32 | -52 | 54 |
|  | R | 31 | 4.26 | 4.12 | 0.002 | 34 | -52 | 50 |
|  | R | 27 | 3.69 | 3.59 | 0.008 | 20 | -64 | 44 |
|  | R | 12 | 3.64 | 3.55 | 0.009 | 34 | -78 | 46 |
| **rDLPFC-activated group vs. sham group, Contrast: Post 3s (IS > NP) – Pre 3s (IS > NP)** |  |  |  |  |  |  |  |  |
| **DLPFC** | R | 121 | 5.36 | 5.10 | 0.001 | 4 | 42 | 30 |
|  | R | 50 | 4.95 | 4.74 | 0.001 | 56 | 34 | 16 |
|  | L | 35 | 4.64 | 4.46 | 0.002 | -2 | 42 | 30 |
|  | R | 47 | 3.68 | 3.59 | 0.012 | 42 | 16 | 38 |
| **PPC** | R | 49 | 4.53 | 4.36 | 0.003 | 16 | -52 | 50 |

Data are thresholded at $p(\text{FDR}) < 0.05$ , with a minimum cluster size of 10 voxels (For all analyses involving multiple comparisons, the $p$ -values were further adjusted using an additional FDR correction). R: right. L: left.

**Extended data Table 6.**
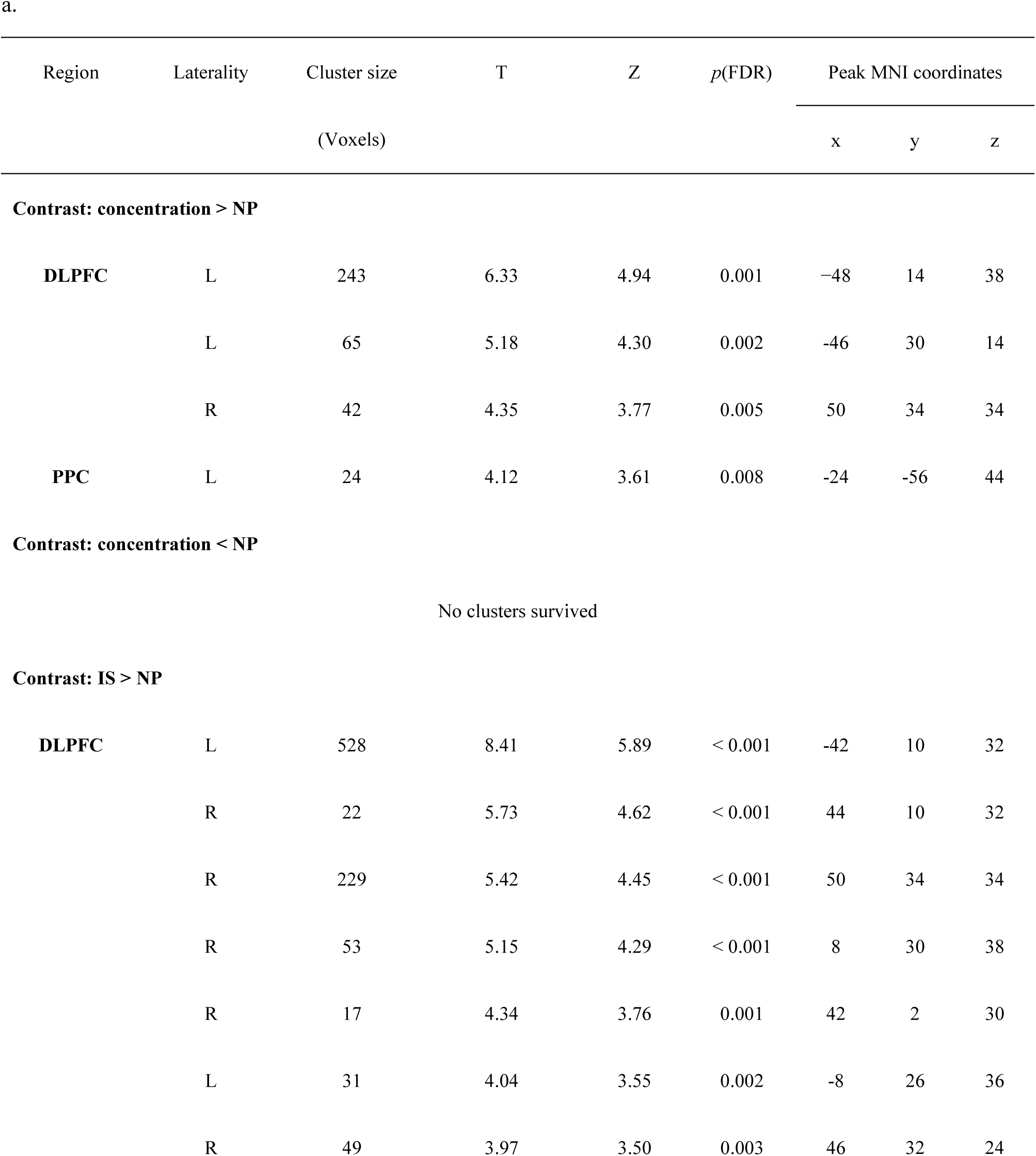
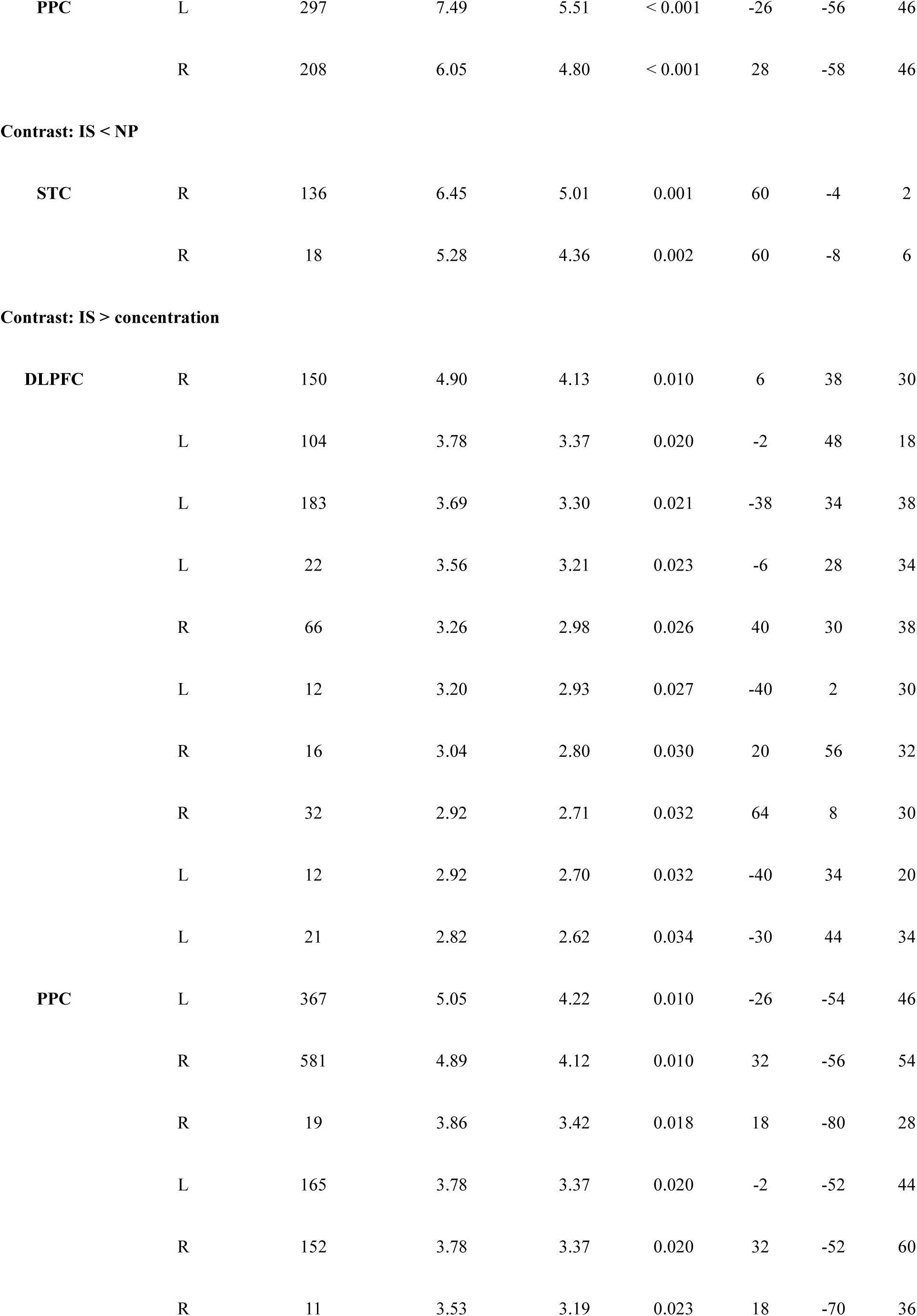
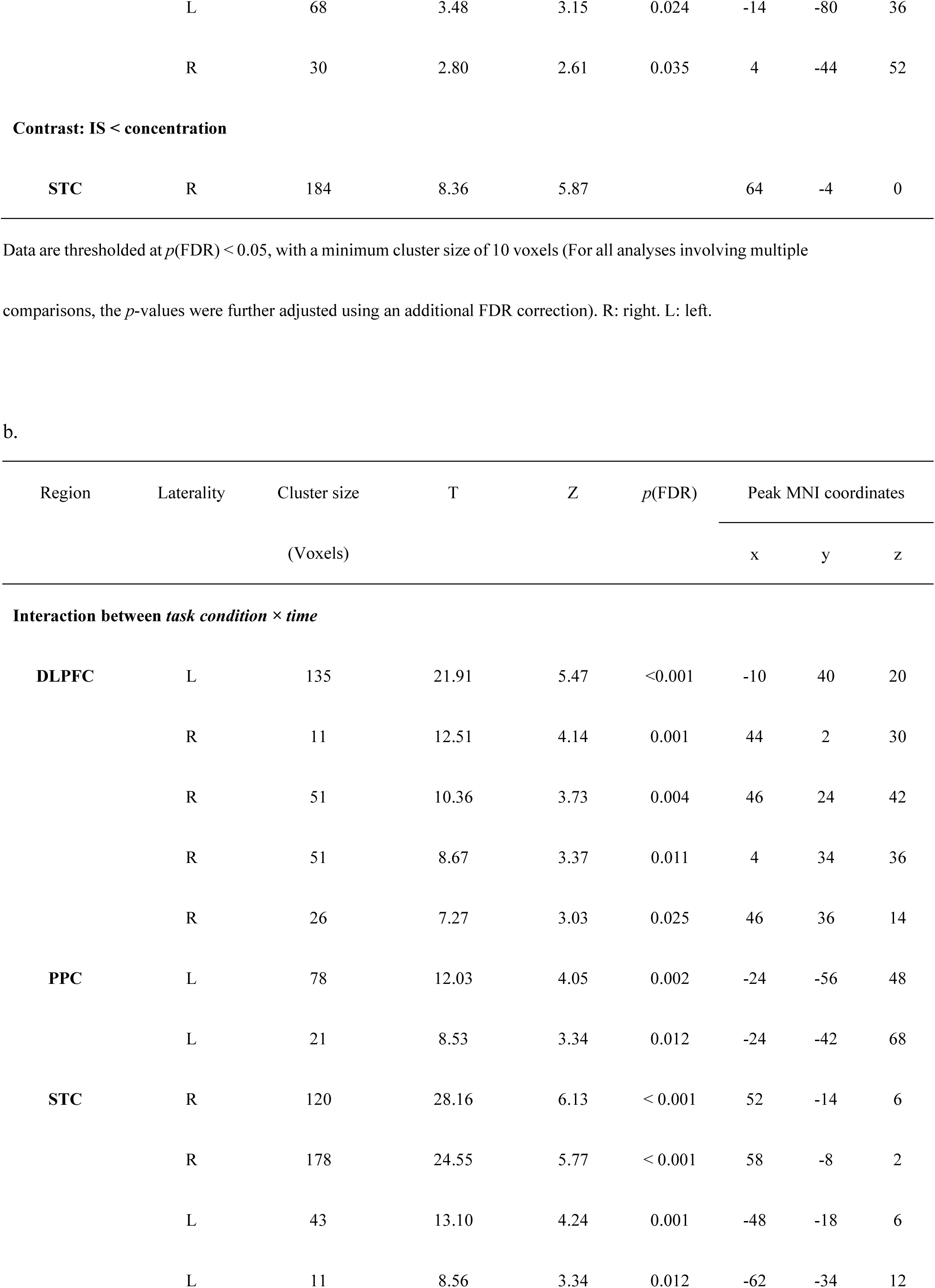
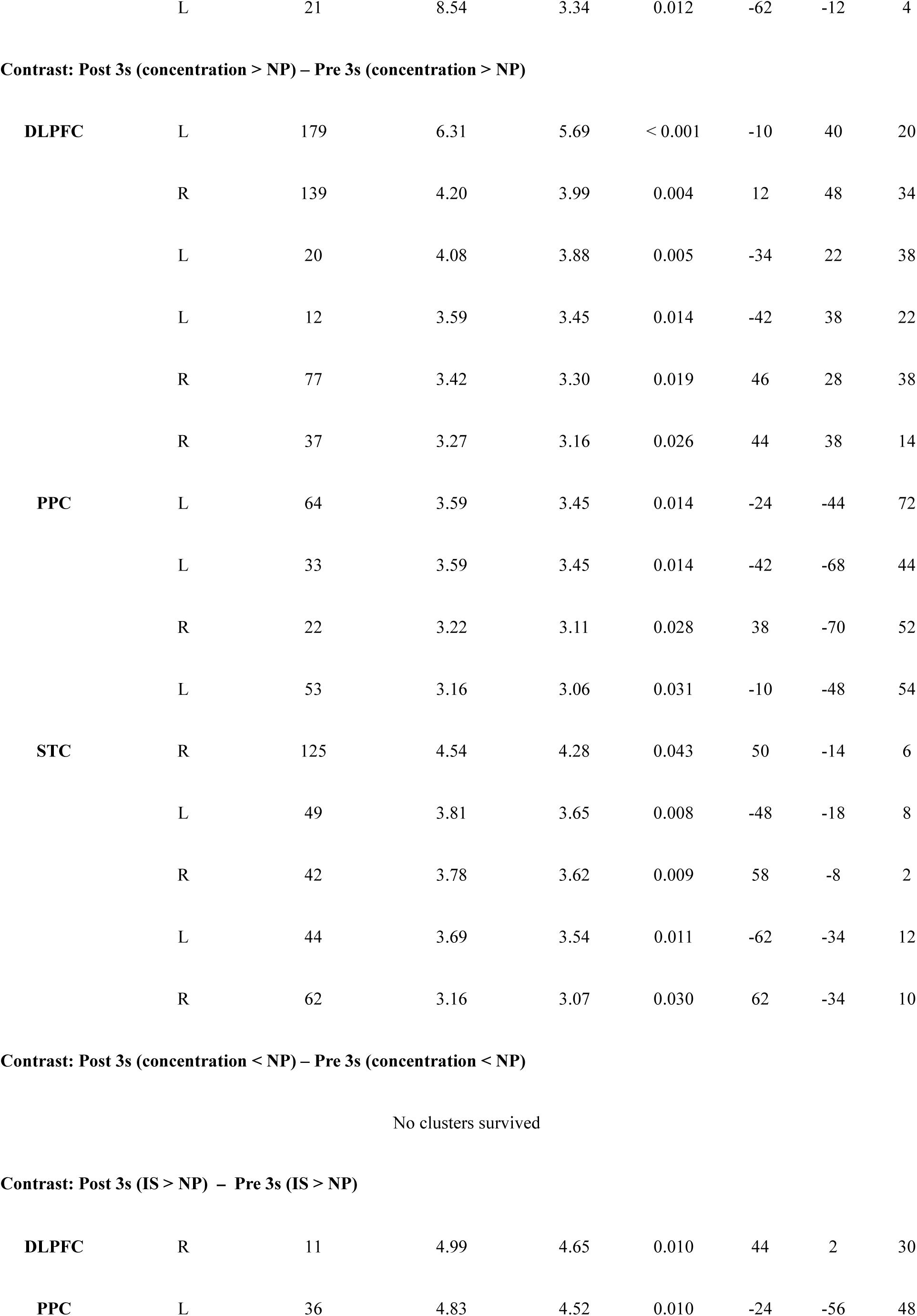
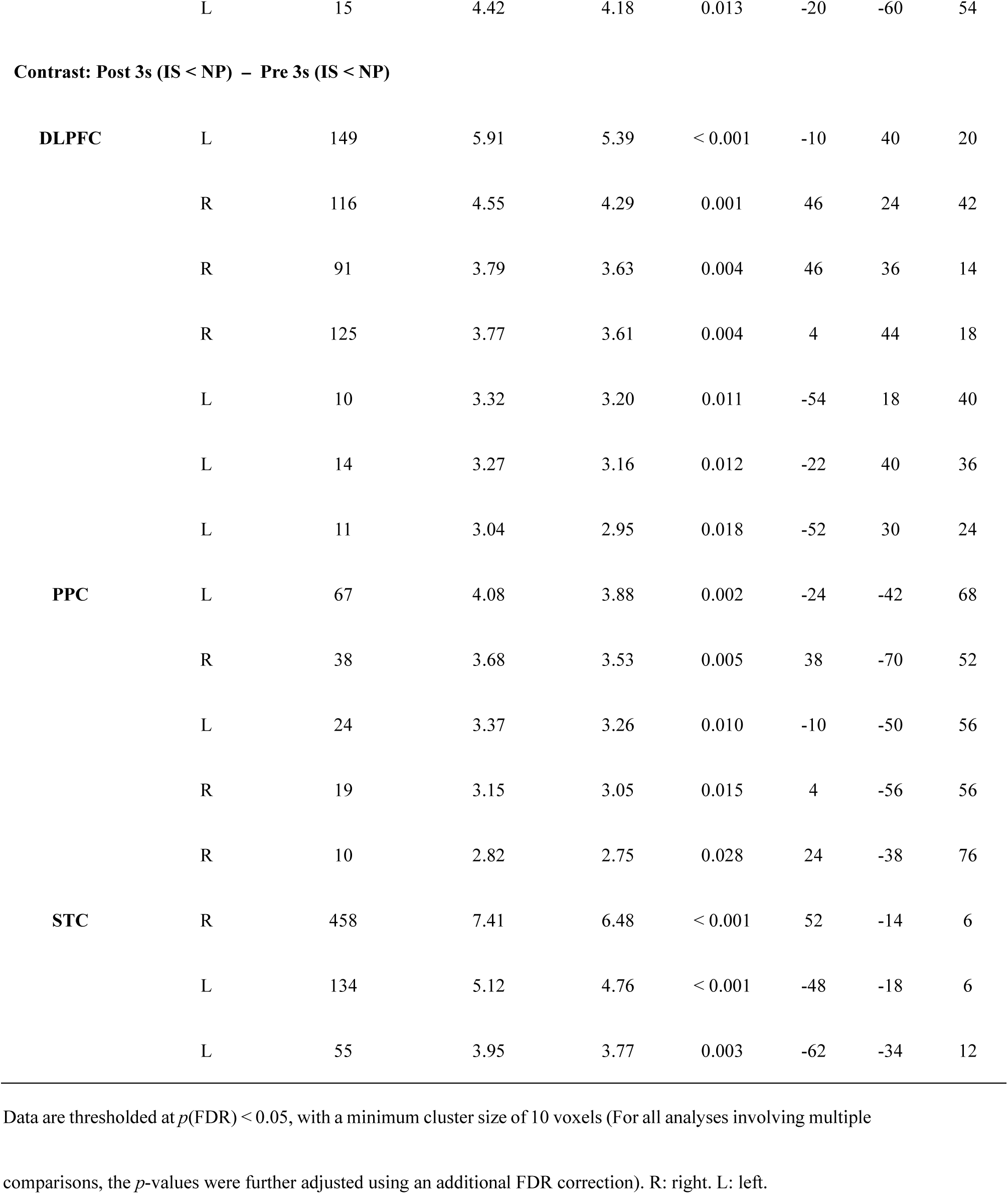
Significant activation clusters within the combined ROI (DLPFC, PPC, STC) in the “Concentration model” of Experiment 4: Analysis of voice exposure periods **(a)** 0-6 seconds, and **(b)** difference of post-3s vs. pre-3s period.

**Extended data Table 7.**
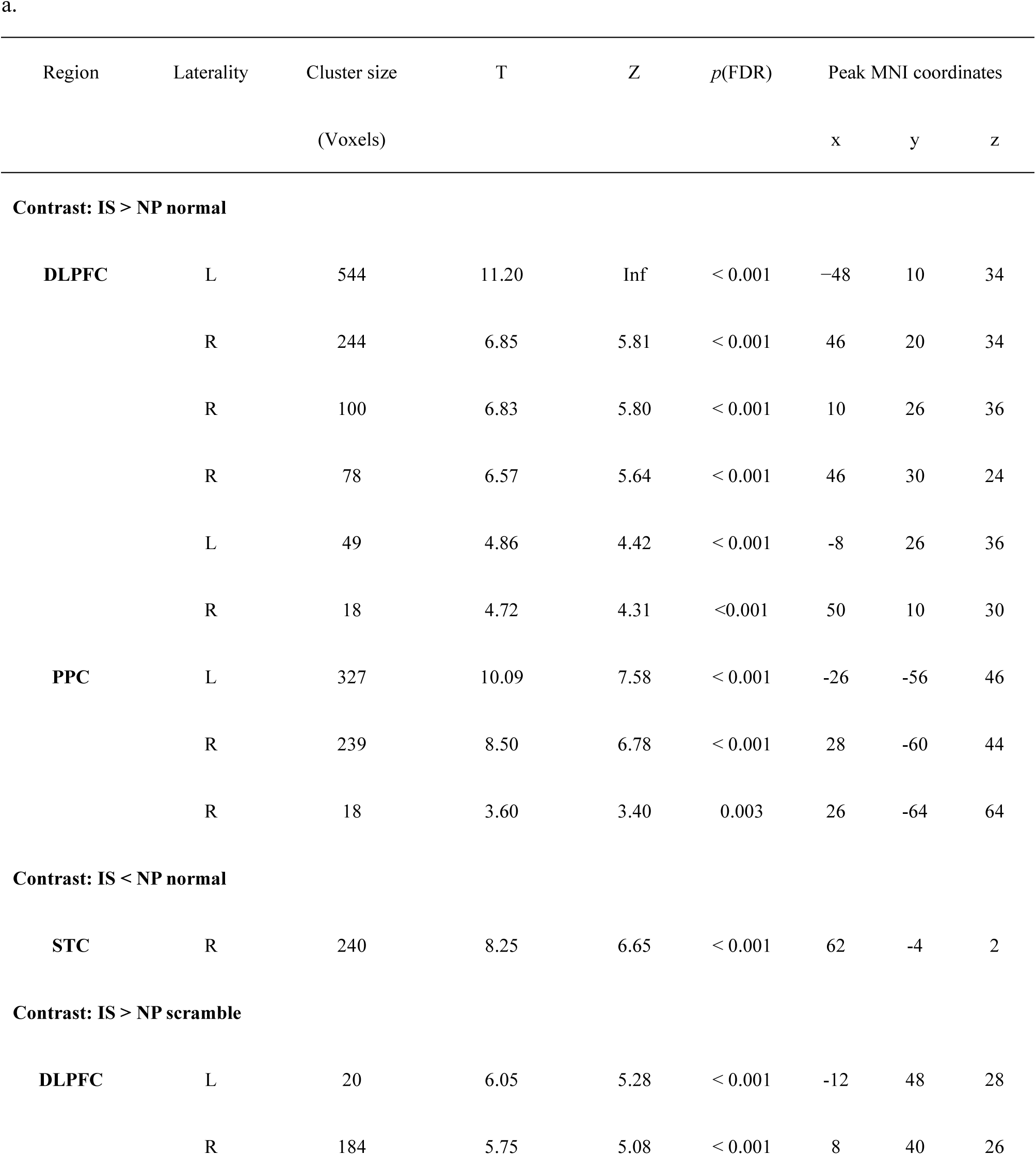
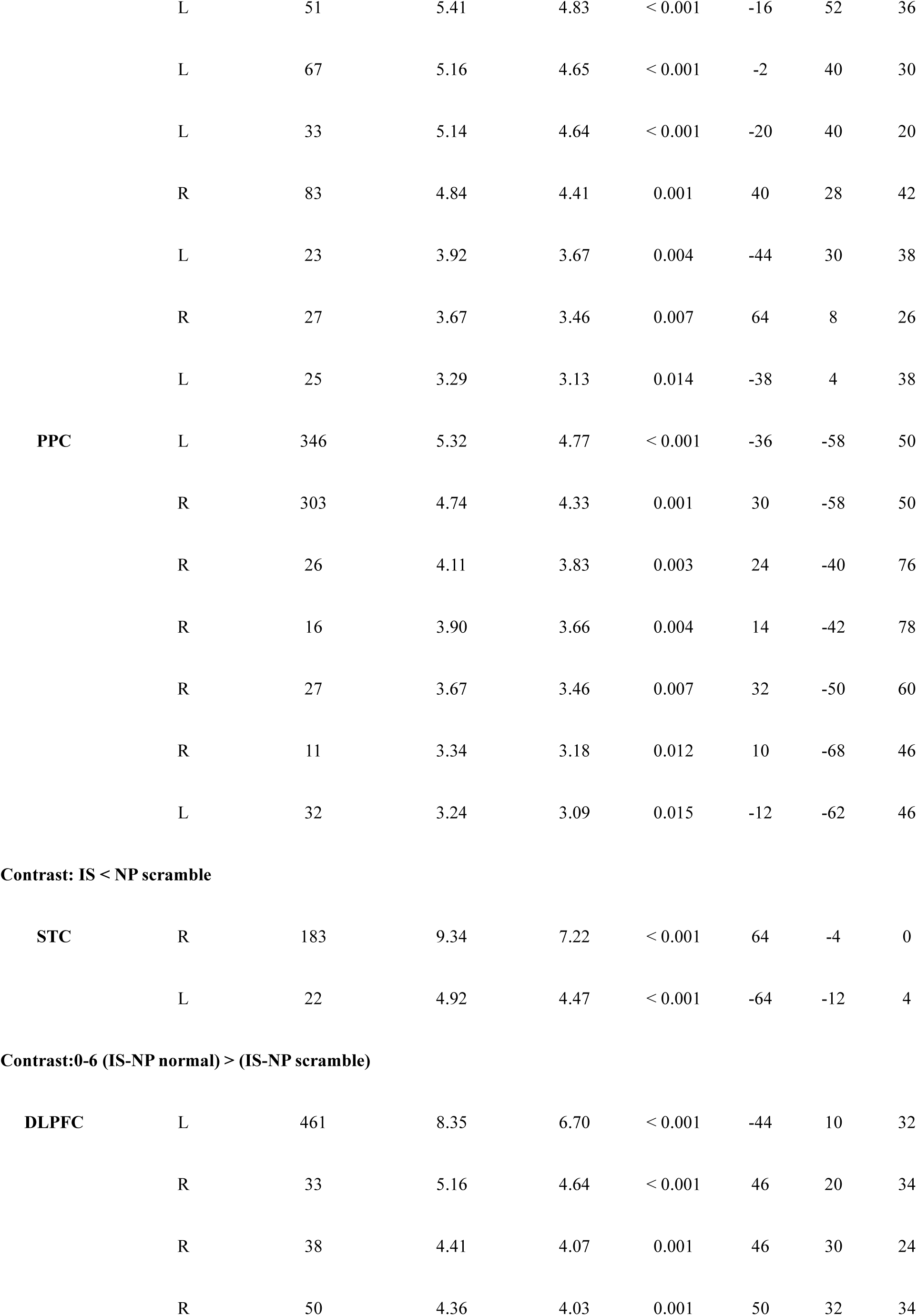
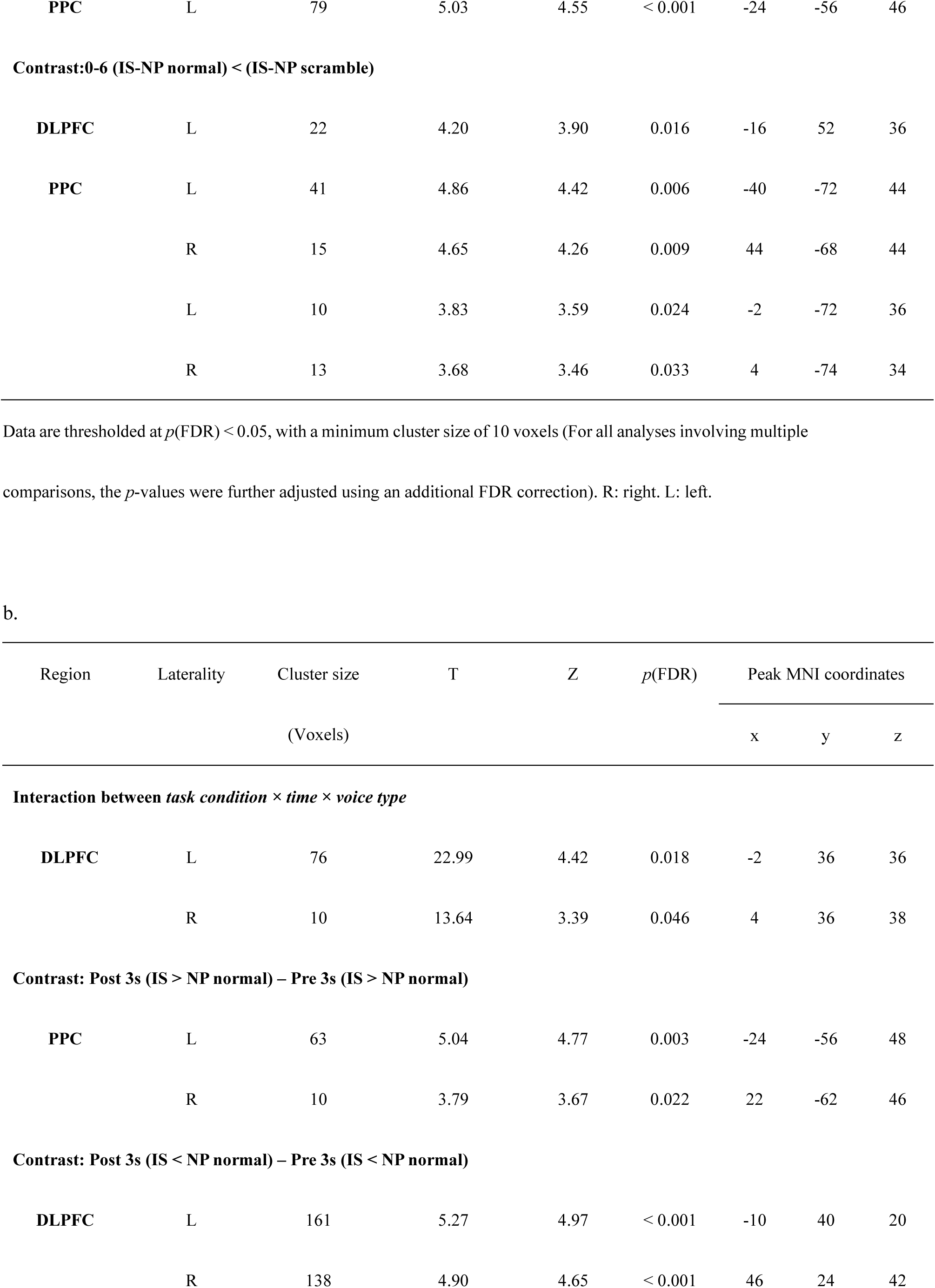
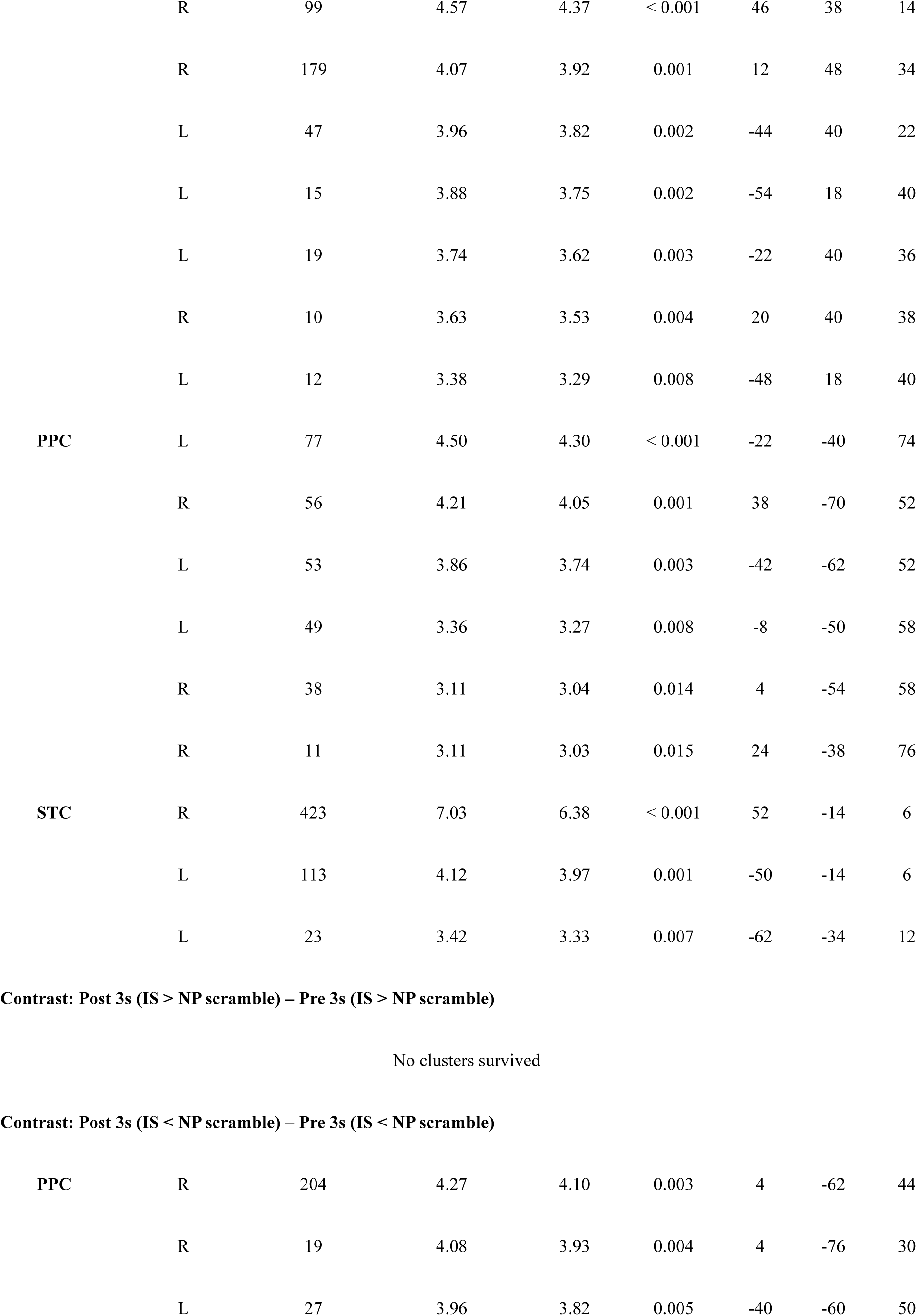
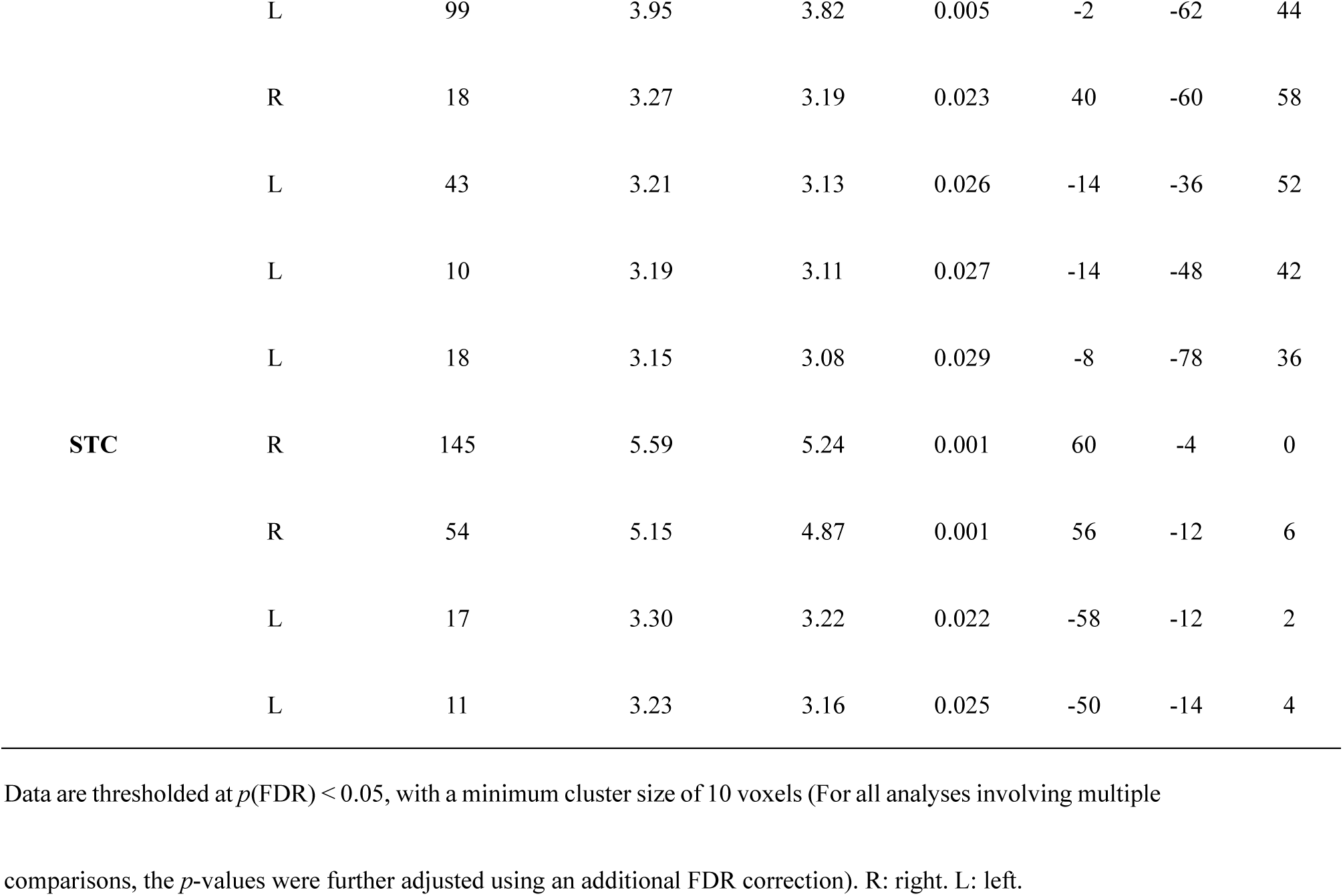
Significant activation clusters within the combined ROI (DLPFC, PPC, STC) in the “Scramble model” of Experiment 4: Analysis of voice exposure periods **(a)** 0-6 seconds, and **(b)** difference of post-3s vs. pre-3s period.

| Region | Laterality | Cluster size<br>(Voxels) | T | Z | *p*(FDR) | Peak MNI coordinates |  |  |
| --- | --- | --- | --- | --- | --- | --- | --- | --- |
|  |  |  |  |  |  | x | y | z |
| **Contrast: IS > NP normal** |  |  |  |  |  |  |  |  |
| **DLPFC** | L | 544 | 11.20 | Inf | < 0.001 | -48 | 10 | 34 |
|  | R | 244 | 6.85 | 5.81 | < 0.001 | 46 | 20 | 34 |
|  | R | 100 | 6.83 | 5.80 | < 0.001 | 10 | 26 | 36 |
|  | R | 78 | 6.57 | 5.64 | < 0.001 | 46 | 30 | 24 |
|  | L | 49 | 4.86 | 4.42 | < 0.001 | -8 | 26 | 36 |
|  | R | 18 | 4.72 | 4.31 | < 0.001 | 50 | 10 | 30 |
| **PPC** | L | 327 | 10.09 | 7.58 | < 0.001 | -26 | -56 | 46 |
|  | R | 239 | 8.50 | 6.78 | < 0.001 | 28 | -60 | 44 |
|  | R | 18 | 3.60 | 3.40 | 0.003 | 26 | -64 | 64 |
| **Contrast: IS < NP normal** |  |  |  |  |  |  |  |  |
| **STC** | R | 240 | 8.25 | 6.65 | < 0.001 | 62 | -4 | 2 |
| **Contrast: IS > NP scramble** |  |  |  |  |  |  |  |  |
| **DLPFC** | L | 20 | 6.05 | 5.28 | < 0.001 | -12 | 48 | 28 |
|  | R | 184 | 5.75 | 5.08 | < 0.001 | 8 | 40 | 26 |
|  | L | 51 | 5.41 | 4.83 | < 0.001 | -16 | 52 | 36 |
|  | L | 67 | 5.16 | 4.65 | < 0.001 | -2 | 40 | 30 |
|  | L | 33 | 5.14 | 4.64 | < 0.001 | -20 | 40 | 20 |
|  | R | 83 | 4.84 | 4.41 | 0.001 | 40 | 28 | 42 |
|  | L | 23 | 3.92 | 3.67 | 0.004 | -44 | 30 | 38 |
|  | R | 27 | 3.67 | 3.46 | 0.007 | 64 | 8 | 26 |
|  | L | 25 | 3.29 | 3.13 | 0.014 | -38 | 4 | 38 |
| **PPC** | L | 346 | 5.32 | 4.77 | < 0.001 | -36 | -58 | 50 |
|  | R | 303 | 4.74 | 4.33 | 0.001 | 30 | -58 | 50 |
|  | R | 26 | 4.11 | 3.83 | 0.003 | 24 | -40 | 76 |
|  | R | 16 | 3.90 | 3.66 | 0.004 | 14 | -42 | 78 |
|  | R | 27 | 3.67 | 3.46 | 0.007 | 32 | -50 | 60 |
|  | R | 11 | 3.34 | 3.18 | 0.012 | 10 | -68 | 46 |
|  | L | 32 | 3.24 | 3.09 | 0.015 | -12 | -62 | 46 |
| **Contrast: IS < NP scramble** |  |  |  |  |  |  |  |  |
| **STC** | R | 183 | 9.34 | 7.22 | < 0.001 | 64 | -4 | 0 |
|  | L | 22 | 4.92 | 4.47 | < 0.001 | -64 | -12 | 4 |
| **Contrast:0-6 (IS-NP normal) > (IS-NP scramble)** |  |  |  |  |  |  |  |  |
| **DLPFC** | L | 461 | 8.35 | 6.70 | < 0.001 | -44 | 10 | 32 |
|  | R | 33 | 5.16 | 4.64 | < 0.001 | 46 | 20 | 34 |
|  | R | 38 | 4.41 | 4.07 | 0.001 | 46 | 30 | 24 |
|  | R | 50 | 4.36 | 4.03 | 0.001 | 50 | 32 | 34 |
| PPC | L | 79 | 5.03 | 4.55 | < 0.001 | -24 | -56 | 46 |
| **Contrast:0-6 (IS-NP normal) < (IS-NP scramble)** |  |  |  |  |  |  |  |  |
| DLPFC | L | 22 | 4.20 | 3.90 | 0.016 | -16 | 52 | 36 |
| PPC | L | 41 | 4.86 | 4.42 | 0.006 | -40 | -72 | 44 |
|  | R | 15 | 4.65 | 4.26 | 0.009 | 44 | -68 | 44 |
|  | L | 10 | 3.83 | 3.59 | 0.024 | -2 | -72 | 36 |
|  | R | 13 | 3.68 | 3.46 | 0.033 | 4 | -74 | 34 |

| Region | Laterality | Cluster size<br>(Voxels) | T | Z | $p(\text{FDR})$ | Peak MNI coordinates | | |
| --- | --- | --- | --- | --- | --- | --- | --- | --- |
|  |  |  |  |  |  | x | y | z |
| Interaction between *task condition* × *time* × *voice type* | | | | | | | | |
| DLPFC | L | 76 | 22.99 | 4.42 | 0.018 | -2 | 36 | 36 |
|  | R | 10 | 13.64 | 3.39 | 0.046 | 4 | 36 | 38 |
| Contrast: Post 3s (IS > NP normal) – Pre 3s (IS > NP normal) |  |  |  |  |  |  |  |  |
| PPC | L | 63 | 5.04 | 4.77 | 0.003 | -24 | -56 | 48 |
|  | R | 10 | 3.79 | 3.67 | 0.022 | 22 | -62 | 46 |
| Contrast: Post 3s (IS < NP normal) – Pre 3s (IS < NP normal) |  |  |  |  |  |  |  |  |
| DLPFC | L | 161 | 5.27 | 4.97 | < 0.001 | -10 | 40 | 20 |
|  | R | 138 | 4.90 | 4.65 | < 0.001 | 46 | 24 | 42 |
|  | R | 99 | 4.57 | 4.37 | < 0.001 | 46 | 38 | 14 |
|  | R | 179 | 4.07 | 3.92 | 0.001 | 12 | 48 | 34 |
|  | L | 47 | 3.96 | 3.82 | 0.002 | -44 | 40 | 22 |
|  | L | 15 | 3.88 | 3.75 | 0.002 | -54 | 18 | 40 |
|  | L | 19 | 3.74 | 3.62 | 0.003 | -22 | 40 | 36 |
|  | R | 10 | 3.63 | 3.53 | 0.004 | 20 | 40 | 38 |
|  | L | 12 | 3.38 | 3.29 | 0.008 | -48 | 18 | 40 |
| **PPC** | L | 77 | 4.50 | 4.30 | < 0.001 | -22 | -40 | 74 |
|  | R | 56 | 4.21 | 4.05 | 0.001 | 38 | -70 | 52 |
|  | L | 53 | 3.86 | 3.74 | 0.003 | -42 | -62 | 52 |
|  | L | 49 | 3.36 | 3.27 | 0.008 | -8 | -50 | 58 |
|  | R | 38 | 3.11 | 3.04 | 0.014 | 4 | -54 | 58 |
|  | R | 11 | 3.11 | 3.03 | 0.015 | 24 | -38 | 76 |
| **STC** | R | 423 | 7.03 | 6.38 | < 0.001 | 52 | -14 | 6 |
|  | L | 113 | 4.12 | 3.97 | 0.001 | -50 | -14 | 6 |
|  | L | 23 | 3.42 | 3.33 | 0.007 | -62 | -34 | 12 |
| **Contrast: Post 3s (IS < NP scramble) – Pre 3s (IS < NP scramble)** |  |  |  |  |  |  |  |  |
| **PPC** | R | 204 | 4.27 | 4.10 | 0.003 | 4 | -62 | 44 |
|  | R | 19 | 4.08 | 3.93 | 0.004 | 4 | -76 | 30 |
|  | L | 27 | 3.96 | 3.82 | 0.005 | -40 | -60 | 50 |
|  | L | 99 | 3.95 | 3.82 | 0.005 | -2 | -62 | 44 |
|  | R | 18 | 3.27 | 3.19 | 0.023 | 40 | -60 | 58 |
|  | L | 43 | 3.21 | 3.13 | 0.026 | -14 | -36 | 52 |
|  | L | 10 | 3.19 | 3.11 | 0.027 | -14 | -48 | 42 |
|  | L | 18 | 3.15 | 3.08 | 0.029 | -8 | -78 | 36 |
| STC | R | 145 | 5.59 | 5.24 | 0.001 | 60 | -4 | 0 |
|  | R | 54 | 5.15 | 4.87 | 0.001 | 56 | -12 | 6 |
|  | L | 17 | 3.30 | 3.22 | 0.022 | -58 | -12 | 2 |
|  | L | 11 | 3.23 | 3.16 | 0.025 | -50 | -14 | 4 |

Data are thresholded at $p(\text{FDR}) < 0.05$ , with a minimum cluster size of 10 voxels (For all analyses involving multiple comparisons, the $p$ -values were further adjusted using an additional FDR correction). R: right. L: left.

**Extended data Table 8.** Descriptive statistics of the unimodal (auditory) MVPA for the decoding accuracy (*M* ± *SD*) of each condition across all participants in (a) Experiments 2 and (b) Experiment 3. The decoding accuracy for each condition was statistically tested against chance level (0.5) using one-sample *t*-test.

| Brain region | 0-6 s |  | 0-3 s |  | 3-6 s |  |
| --- | --- | --- | --- | --- | --- | --- |
|  | IS | NP | IS | NP | IS | NP |
| **AIS** |  |  |  |  |  |  |
| DLPFC | $0.61 \pm 0.16^{**}$ | $0.54 \pm 0.13$ | $0.64 \pm 0.14^{***}$ | $0.52 \pm 0.14$ | $0.58 \pm 0.16^*$ | $0.56 \pm 0.17$ |
| PPC | $0.58 \pm 0.14^{**}$ | $0.57 \pm 0.15^*$ | $0.6 \pm 0.16^{**}$ | $0.58 \pm 0.13^{**}$ | $0.57 \pm 0.15^*$ | $0.56 \pm 0.17^*$ |
| STC | $0.57 \pm 0.12^{**}$ | $0.62 \pm 0.13^{***}$ | $0.58 \pm 0.13^{**}$ | $0.58 \pm 0.12^{**}$ | $0.55 \pm 0.16$ | $0.66 \pm 0.16^{***}$ |
| **VIS** |  |  |  |  |  |  |
| DLPFC | $0.70 \pm 0.11^{***}$ | $0.66 \pm 0.14^{***}$ | $0.70 \pm 0.16^{***}$ | $0.63 \pm 0.18^{***}$ | $0.70 \pm 0.16^{***}$ | $0.63 \pm 0.18^{***}$ |
| PPC | $0.70 \pm 0.11^{***}$ | $0.66 \pm 0.15^{***}$ | $0.69 \pm 0.16^{***}$ | $0.66 \pm 0.17^{***}$ | $0.66 \pm 0.14^{***}$ | $0.69 \pm 0.17^{***}$ |
| VC | $0.67 \pm 0.16^{***}$ | $0.70 \pm 0.16^{***}$ | $0.67 \pm 0.19^{***}$ | $0.68 \pm 0.18^{***}$ | $0.66 \pm 0.16^{***}$ | $0.74 \pm 0.17^{***}$ |

**b.**
| Brain<br>region | Left-DLPFC activated group |  |  |  |  |  | Right-DLPFC activated group |  |  |  |  |  | Sham group |  |  |  |  |  |
| --- | --- | --- | --- | --- | --- | --- | --- | --- | --- | --- | --- | --- | --- | --- | --- | --- | --- | --- |
|  | 0-6 s |  | 0-3 s |  | 3-6 s |  | 0-6 s |  | 0-3 s |  | 3-6 s |  | 0-6 s |  | 0-3 s |  | 3-6 s |  |
|  | IS | NP | IS | NP | IS | NP | IS | NP | IS | NP | IS | NP | IS | NP | IS | NP | IS | NP |
| DLPFC | $0.66 \pm 0.11^{**}$ | $0.58 \pm 0.13^*$ | $0.71 \pm 0.12^{**}$ | $0.55 \pm 0.13$ | $0.58 \pm 0.15$ | $0.61 \pm 0.13^*$ | $0.67 \pm 0.08^{***}$ | $0.56 \pm 0.10$ | $0.65 \pm 0.11^{**}$ | $0.53 \pm 0.10$ | $0.58 \pm 0.08^*$ | $0.62 \pm 0.11^*$ | $0.62 \pm 0.10^{**}$ | $0.61 \pm 0.11^*$ | $0.65 \pm 0.14^*$ | $0.62 \pm 0.14^*$ | $0.59 \pm 0.12^*$ | $0.60 \pm 0.09^*$ |
| PPC | $0.60 \pm 0.12^*$ | $0.57 \pm 0.10^*$ | $0.62 \pm 0.12^*$ | $0.59 \pm 0.12^*$ | $0.56 \pm 0.10$ | $0.60 \pm 0.15^*$ | $0.59 \pm 0.08^{**}$ | $0.57 \pm 0.08^*$ | $0.62 \pm 0.11^*$ | $0.57 \pm 0.11$ | $0.57 \pm 0.08^*$ | $0.59 \pm 0.13^*$ | $0.59 \pm 0.13^*$ | $0.62 \pm 0.16^*$ | $0.58 \pm 0.09^*$ | $0.63 \pm 0.08^{**}$ | $0.63 \pm 0.20$ | $0.72 \pm 0.18^*$ |
| STC | $0.56 \pm 0.09^*$ | $0.62 \pm 0.11^{**}$ | $0.59 \pm 0.11^*$ | $0.58 \pm 0.13$ | $0.58 \pm 0.12^*$ | $0.59 \pm 0.14$ | $0.60 \pm 0.14^*$ | $0.61 \pm 0.09^{**}$ | $0.63 \pm 0.14^*$ | $0.57 \pm 0.09^*$ | $0.56 \pm 0.13$ | $0.61 \pm 0.10^*$ | $0.61 \pm 0.10^{**}$ | $0.61 \pm 0.15^*$ | $0.64 \pm 0.18^*$ | $0.62 \pm 0.19$ | $0.59 \pm 0.12^*$ | $0.65 \pm 0.12^*$ |

$*p(\text{FDR}) < 0.05$ , $**p(\text{FDR}) < 0.01$ , $***p(\text{FDR}) < 0.001$

**Extended data Table 9.** Descriptive statistics of feature MVPA decoding accuracies (*M* ± *SD*) in Experiment 1. The decoding accuracy for each condition was statistically tested against chance level (0.5) using one-sample *t*-test.

| Items | IS |  | NP |  |
| --- | --- | --- | --- | --- |
|  | high | low | high | low |
| Valence | $0.34 \pm 0.08^{***}$ | $0.34 \pm 0.09^{***}$ | $0.41 \pm 0.11^{***}$ | $0.36 \pm 0.12^{***}$ |
| Arousal | $0.34 \pm 0.08^{***}$ | $0.31 \pm 0.07^{***}$ | $0.41 \pm 0.08^{***}$ | $0.42 \pm 0.11^{***}$ |
| Information load | $0.44 \pm 0.13^{***}$ | $0.47 \pm 0.14^{***}$ | $0.35 \pm 0.08^{***}$ | $0.31 \pm 0.07^{***}$ |
| Difficulty | $0.33 \pm 0.09^{***}$ | $0.35 \pm 0.10^{***}$ | $0.36 \pm 0.09^{***}$ | $0.30 \pm 0.08^{**}$ |

$^{**}p(\text{FDR}) < 0.01$ , $^{***}p(\text{FDR}) < 0.001$

**Extended data Table 10.** Descriptive statistics of the cross-modal (auditory and visual) MVPA for the decoding accuracy (*M* ± *SD*) of each condition across all participants in Experiments 2. The decoding accuracy for each condition was statistically tested against chance level (0.5) using one-sample *t*-test.

| Brain region | 0-6 s |  | 0-3 s |  | 3-6 s |  |
| --- | --- | --- | --- | --- | --- | --- |
|  | IS | NP | IS | NP | IS | NP |
| **AIS** |  |  |  |  |  |  |
| CEN | $0.65 \pm 0.18^{***}$ | $0.45 \pm 0.16$ | $0.62 \pm 0.14^{***}$ | $0.49 \pm 0.12$ | $0.67 \pm 0.13^{***}$ | $0.42 \pm 0.12$ |
| **VIS** |  |  |  |  |  |  |
| CEN | $0.56 \pm 0.16^{\dagger}$ | $0.49 \pm 0.14$ | $0.54 \pm 0.11$ | $0.53 \pm 0.12$ | $0.58 \pm 0.13^{**}$ | $0.49 \pm 0.11$ |

$^{\dagger}p(\text{FDR})$ marginally significant, $^{**}p(\text{FDR}) < 0.01$ , $^{***}p(\text{FDR}) < 0.001$

